# Using evolutionary constraint to define novel candidate driver genes in medulloblastoma

**DOI:** 10.1101/2022.11.02.514465

**Authors:** Ananya Roy, Sharadha Sakthikumar, Sergey V. Kozyrev, Jessika Nordin, Raphaela Pensch, Mats Pettersson, Zoonomia Consortium, Elinor Karlsson, Kerstin Lindblad-Toh, Karin Forsberg-Nilsson

## Abstract

Current knowledge of cancer genomics is biased against non-coding mutations. Here, we use whole genome sequencing data from pediatric brain tumors, combined with evolutionary constraint inferred from 240 mammals to identify genes enriched in non-coding constraint mutations (NCCMs). We compare medulloblastoma (MB, malignant) to pilocytic astrocytoma (PA, benign) and find drastically different NCCM frequencies between the two. In PA, a high NCCM frequency only affects the *BRAF* locus, while in MB, >500 genes have high levels of NCCMs. Intriguingly, many genes are associated with different age of onset, such as *HOXB1* in young patients and *NUAK1* in adult patients. Our analysis points to different molecular pathways in different patient groups. These novel candidate driver genes may assist patient stratification in MB and may be useful for treatment options.

**One-Sentence Summary:** Non-coding constraint mutations implicate novel candidate genes to stratify medulloblastoma by age and subgroups.

## MAIN TEXT

Systematic and genome-wide interrogations of human cancers have enabled characterization of their mutational landscapes (*1*) and generated comprehensive pan-cancer multi-omics resources, intersecting mutational profiles with gene expression data (*2*). However, cancer driver discovery has mostly focused on coding mutations, with less attention paid to mutations in the non-coding parts, which make up >98% of the genome. Although most non-coding mutations are expected to be passenger events, mutations in regulatory regions should be examined for their contribution to disease initiation and progression (*3*).

Positions under evolutionary constraint in the genome can be leveraged to identify functional sequences and identify deleterious mutations. In a proof-of-concept study we applied GERP scores, a widely used conservation metric, to discover non-coding constraint mutations (NCCMs) in glioblastoma (GBM) (*4*). Here, for the first time, we apply phyloP, a constraint metric from a sequence alignment of 240 mammals (*5*) to identify genes with increased NCCM burden and potential function in cancer.

We downloaded variant calls from whole-genome sequencing (WGS) data from the International Cancer Genome Consortium (ICGC) (*2*) (*6*) for the two most common primary brain tumors in children, pilocytic astrocytoma (PA, 89 cases, (table S1 (*7*), which is benign, and medulloblastoma (MB,146 cases (table S2) (*6*), which is malignant. Total mutational counts, when normalized by patient numbers, was significantly more frequent for MB than PA, reflecting their different grades, (fig. S1A). To investigate mutations in the non-coding part of the genome for potential functional impact, somatic alterations in introns, 5’ and 3’ UTRs, as well as intergenic regions ±100 *kbp* upstream and downstream of each gene (excluding positions in protein-coding sequence), were analyzed for NCCMs.

We first examined PA, where *BRAF* fusions are the main events (*8*). Our NCCM analysis showed only one gene, *NDUFB2* with ≥2 NCCMs/100 *kbp*, a previously used cutoff (*4*). *NDUFB2* is located in the same locus as *BRAF* and *ADCK2*, both with ≥1 NCCMs/100 *kbp*, sharing these NCCMs (fig. S2 A-B). We found 4 NCCMs in genes previously reported to be significantly mutated in PA (fig. S2C, table S3). Examining 9 additional genes with ≥1 NCCMs/100 *kbp* (table S4), we note the nutrient sensor OGT, a kinase which is overexpressed in PA (*9*). Significantly mutated genes that have alterations in protein coding sequence (table S3) and the genes carrying ≥1 NCCMs/100 *kbp* (table S4) were jointly subjected to GSE analysis, which revealed enrichment for ‘diseases of signal transduction’ (*p*-value = 1.8×10^-4^) and ‘oncogenic MAPK signaling’ (*p*-value = 1.7×10^-3^) (table S5), supporting the relevance of NCCM enrichment for PA.

In the MB cohort, 114 genes had high NCCM occurrence rate, i.e., ≥ 2 NCCMs/100 *kbp* (fig. S3A, table S6-7). To complement this weighted analysis, we also performed a simplified analysis identifying 525 genes with ≥ 5 NCCMs (fig. S3B, table S8). These sets together (n=530, fig. S3C) were enriched for ‘nervous system development’ (*p*-value =1.3×10^-26^) and ‘generation of neurons’ (*p*-value =1.7×10^-22^) as shown by GSE analysis (table S9). In the majority of the genes with ≥ 2 NCCMs/100 kbp, NCCMs were evenly distributed across patients, with one NCCM per patient.

In 19 % of the 114 genes with highest NCCM rates, up to two NCCMs were found in the same patient (table S7). There were few coding mutations in the top 114 (table S10), and the genes with protein-coding mutations had low NCCM rates (table S11). Thus, NCCMs and coding mutations generally do not affect the same set of genes, and our analysis reveals a large number of new mutations, and genes of interest.

In line with the total mutational burden being higher for MB than for PA, cohort-wide normalization of NCCMs showed that MB had greater NCCM accumulation rates than PA. This was the case both when comparing all genes (fig S4A) and the top 0.5 % of genes with NCCMs, in each cohort (fig S4B).

Querying the number of NCCMs in known MB driver pathways revealed 29 NCCMs in 9 genes of the SHH pathway (of which 9 were in SHH group patients), 28 NCCMs in 10 out of 12 genes of the WNT pathway and 22 NCCMs in 10 genes in the MYC pathway (table S12, see supplementary information for description of MB subgroups). Many of these mutations were distributed across the different subgroups, showing a considerable overlap of pathways. SHH tumors had the highest overall count of these NCCMs, but in Group 4 the rate per patient was highest (fig. S5A-B). We also note a demarcation between NCCM-containing genes occurring predominantly in children (<18 years), or adults (≥18 years) (fig. S6). There are distinct early and late-onset genes suggesting different driver genes in the different age groups (Fig. 1A. There is a highly significant negative correlation between adult and pediatric NCCMs/100 Kbp per patient (Fig, 1B). In addition, we find a positive correlation when comparing adult patient NCCMs and SHH group NCCMs (Fig. 1C, fig S7).

**Figure 1.**
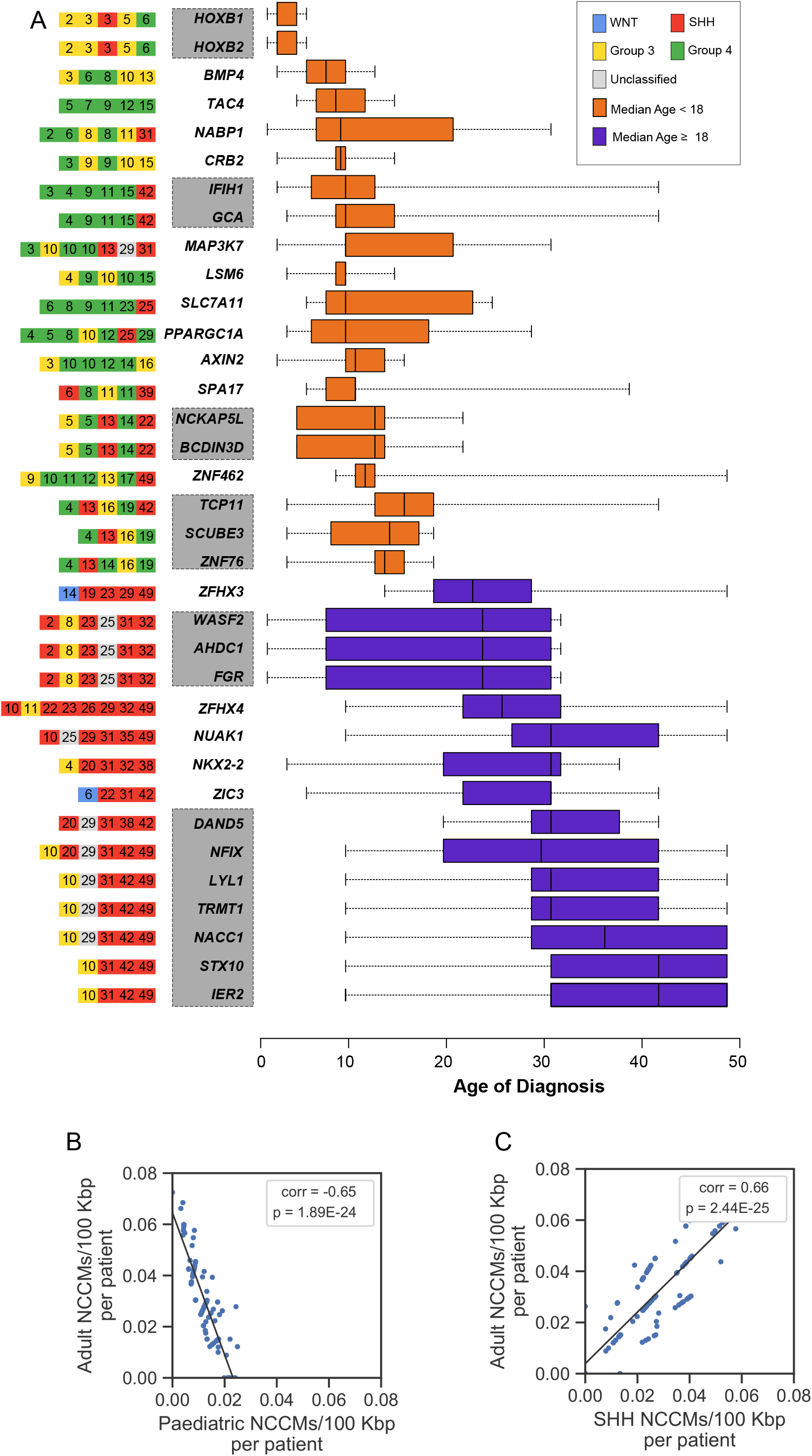
Candidate driver genes for MB have ≥ 2 NCCMs per 100 *kbp* and are differently distributed across pediatric and adult MB. **A)** Left, subgroup (by color code), and age (denoted by the number in the squares) for each patient. Right, Age of onset distribution. Orange, median age <18, purple, median age >18. Grey boxes denote genes in the same locus (See figure S5 for all 114 genes). **B)** Negative correlation in NCCMs/100 Kbp per patient between adult and pediatric patients. **C)** Positive correlation between NCCMs in adult and SHH group.

The vast majority, 81%, of pediatric MB patients had at least one NCCM, and 39% had NCCMs in more than one locus (Fig. S8). In the paediatric cohort, most genes with NCCMs are associated with Group 3 and Group 4 (Fig. 1). These subgroups remain difficult to distinguish because few group-specific recurrent protein-coding mutations have been reported (*10*) (*6*). We therefore examined the NCCM-containing genes for possible preponderance towards either of the subgroups. *BMP4, CRB2, LSM6* and *AXIN2* had NCCMs exclusively in Group 3 and 4 patients, while no NCCMs were private to Group 3. *TAC4* was exclusive to Group 4, and the loci for *IFIH1/GCA* and *SLC7A11* to Group 4 and one SHH patient. *SLC7A11* is linked to seizures in glioma (*11*), and intriguingly, four of the *SLC7A11* NCCMs are shared with its corresponding *SLC7A11* antisense transcript, showing regulatory potential. Because 24% of Group 3 and 18% of Group 4 cases lack potential coding driver mutations (*6*), the above NCCM information could identify new candidates for these subgroups.

Mutations in the promoter region of the telomerase reverse transcriptase (TERT) gene have been described as a common recurrent somatic mutation in MB (*12*) with predominant enrichment for SHH and WNT groups in adult patients. In the current cohort, 5 out of the 37 patients (14%), all belonging to the SHH group have mutations at the chr5:1,295,228 position but none at the chr5:1,295,250 position. There were no other mutated positions that are close to the promoter region in this dataset.

Among the youngest patients we find NCCMs in the loci for the *HOXB* gene family, and *BMP4*. BMP4 is a pivotal regulator of neural development that induces differentiation of cerebellar progenitors (*13*) and has been proposed to suppress MB in mice (*14*). Of five *BMP4* NCCMs (Fig. 2A), we selected NCCM3 (Fig. 2B) for further studies because this position shows high conservation in multiple transcription factor binding sites, e.g. SOX10 (Fig. 2C) and because sTRAP analysis predicted increased affinity, compared to the wild-type sequence (Fig. 2D). This prediction was confirmed by reporter assay (Fig. 2E) and electrophoretic mobility shift assay (EMSA) using nuclear extracts from the MB cell line MB002 (Fig. 2F). Next, we queried a published dataset (*15*) for expression of *BMP4*, comparing MB and normal cerebellum. Group 3, Group 4, and SHH patients had lower *BMP4* expression than normal cerebellum, while it was higher in the WNT group (Fig. 2G). Interestingly, NCCMs in the *TCP11/SCUBE3/ZNF76* locus (Fig. 1) supports the notion of the BMP4 pathway as a potential MB candidate gene, because SCUBE3, a glycoprotein anchored to the cell surface, is a coreceptor for BMP2/BMP4 (*16*).

**Figure 2.**
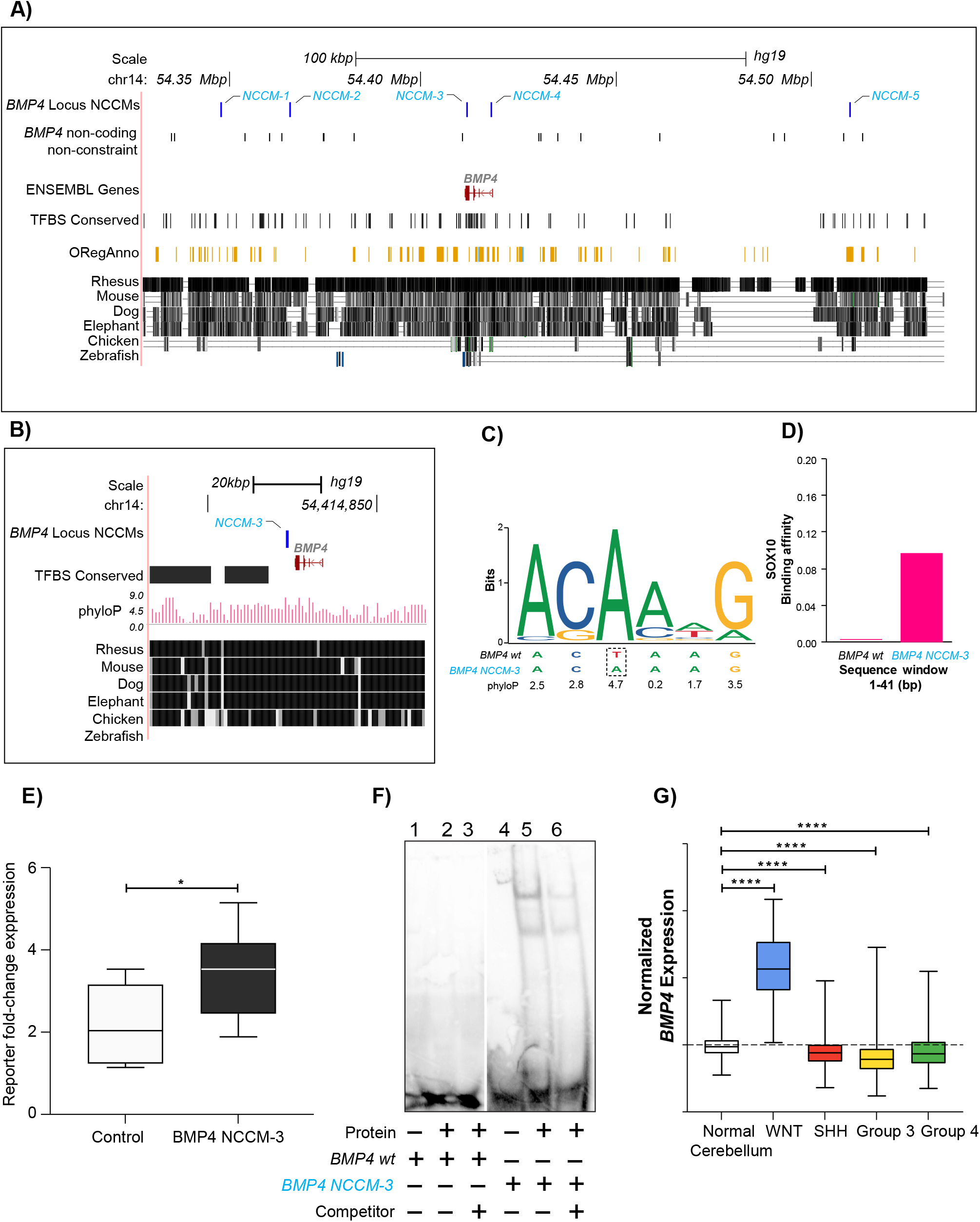
*BMP4*, a possible candidate gene for young MB patients. **A)** The five *BMP4*-NCCMs (blue) all lie in regions of high constraint and has at least one regulatory annotation. Coding mutations and non-coding mutations are show on individual rows. **B)** Zoomed-in view of *BMP4*-NCCM3 shows high phyloP score. **C)** SOX10 binding logo shows *BMP4*-NCCM3 in a highly conserved position. **D)** Bioinformatic prediction is that SOX10 affinity for the mutated versus wild-type sequence. **E**) Reporter assay shows significantly increased expression of the reporter gene for the mutant allele for NCCM3 compared to the wild type in MB002 cells. **F)** EMSA with nuclear protein extract from MB002 cells shows increased binding to the mutant (lane 5-6) compared to wild-type sequence (lane 2-3). Unlabeled dsDNA was used as competitor (lane 3,6). **G)** Expression of *BMP4* in normal cerebellum and MB patients. Boxes min-max range, horizontal lines median value, and whiskers extend to extreme values.

HOXB proteins are instrumental in hind brain development (*17*) where they spatially restrict neural progenitors to ensure segmental ordering of the rhombomeres. Five patients had NCCMS in the *HOXB* locus (Fig. 3A) and these are also shared with several *HOXB*-antisense transcripts. sTRAP analysis predicted that *HOXB*-NCCM3 increase binding of FOXD1, FOXD3 and FOXF2, and *HOXB*-NCCM5 of PAX6 and POU5F1/OCT4 (table S13). We confirmed differential binding to the regions of NCCM1, NCCM3 and NCCM5 using EMSAs (Fig. 3B). Altered reporter transcript expression for the mutant alleles of NCCM1, NCCM3 and NCCM5 were also shown (Fig. 3C). Of the *HOXB* locus genes, *HOXB2* stands out since SHH and Group 4 patients have lower *HOXB2* expression compared to normal cerebellum (*15*) (Fig. 3D), and patients with low *HOXB2* expression have shorter survival (p=3.4×10^-4^) (Fig. 3E). Examining *HOXB2* in the literature, we noted that DDX3X has tumor suppressor capacity in the hindbrain by regulating expression of e.g. *Hoxb2* (*18*). *DDX3X* is frequently mutated in WNT and SHH MB (*18*), which supports the importance of *HOXB* mutations in MB, but until now it has not been described for Group 3 and Group 4 patients. To further characterize the NCCM effects on the *HOXB* locus, we generated CRISPR/Cas9-edited DAOY MB cells and found that introduction of NCCM3, but not NCCM1, increased the expression of antisense transcripts *HOXB*-AS2 and *HOXB*-AS3 (Fig 3F). In addition, introduction of NCCM1 and NCCM3 both resulted in reduced expression of *HOXB2, HOXB5* and *HOXB9* (Fig. 3G). Proliferation assays reflect a lower proliferation rate of DAOY cells with NCCM3, whereas for NCCM1 it was comparable to control, non-edited, cells (Fig. 3H). Thus, a complex regulatory pattern for the *HOXB* locus with multiple genes affected by NCCMs was demonstrated, including alterations in proliferation.

**Figure 3.**
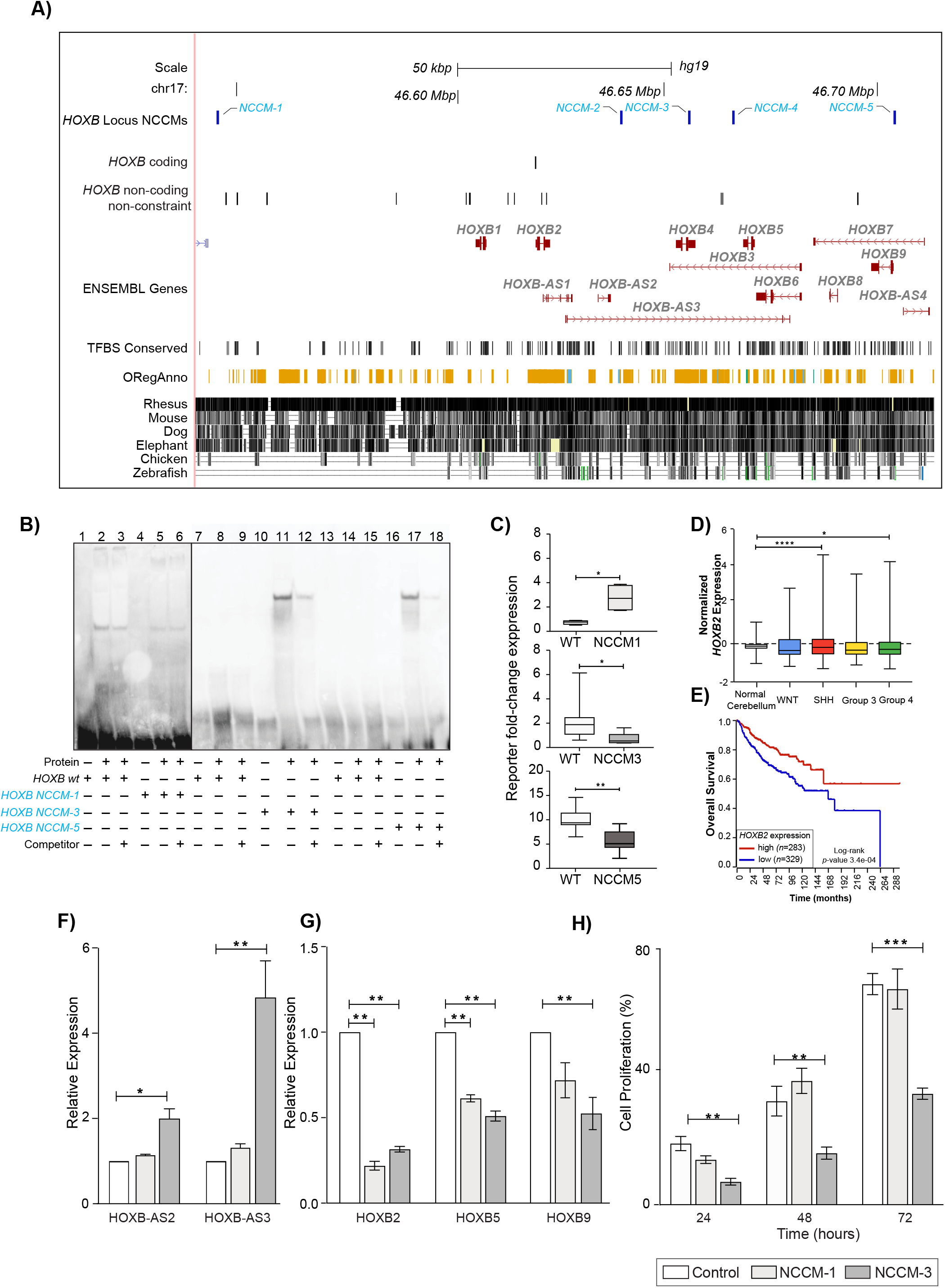
The *HOXB* gene cluster contains five highly constraint NCCMs. **A)** Five NCCMs (blue) are found in the *HOXB* locus, which contains nine protein-coding genes and multiple anti-sense transcripts. These NCCMs are highly constraint and have at least one regulatory annotation each. **B)** EMSAs with nuclear protein extracts from MB002 cells show slightly decreased binding for NCCM1 (lanes 5-6) compared to wild-type (lane 2-3). In contrast, an increased binding to *HOXB*-NCCM3 (lane 11-12) and *HOXB*-NCCM5 (lane 17-18), compared to wild-type sequences (lane 8-9 and 14-15 respectively) is seen. Unlabeled dsDNA was used as competitor (lane 3,6,9,12,15,18). **C)** Reporter assay shows increased expression of the mutant allele for NCCM1 (upper panel), and decreased expression of the mutant alleles for NCCM3 (middle panel) and NCCM5 (lower panel). **D)** Expression of *HOXB2* in normal cerebellum and MB patients. Boxes min-max range, horizontal lines median value, and whiskers extend to extreme values. **E)** Kaplan-Meier plot showing that patients with lower *HOXB2* expression have shorter survival than those with high expression. **F)** Upon CRISPR/Cas9-editing, the expression of *HOXB* antisense transcript 2 (*HOXBAS2*) and *HOXBAS3*, is increased as a consequence of inserting NCCM3 in DAOY cells. The NCCM1 did not show such an effect. **G)** DAOY cells edited to contain NCCM1 or NCCM3 show reduced expression of *HOXB2, HOXB5* and *HOXB9*. **H)** Proliferation assay comparing CRISPR/Cas9 cell edits for NCCM1 or NCCM3 to control-edited DAOY cells, showing reduced proliferation by NCCM3 but not by NCCM1.

Most NCCMs associated with adult-onset are found in genes expressed in the brain, many with relatively high expression in cerebellum: *ZFHX4, WASF2AHDC1/FGR, NUAK, ZIC3, DAND5/NFIX/STX10* (https://www.gtexportal.org/home/). There were NCCMs in at least one of these loci (Fig. S9) in 50% of SHH patients. *ZFHX4*, with nine NCCMs in seven patients, either in introns or upstream of the gene, and overlapping a *ZFHX4*-antisense transcript, has been reported to be downregulated in MB (*19*). *ZFHX3*, on the other hand, is normally not expressed in brain, but has been associated with multiple cancers where it acts as a tumor suppressor by downregulating MYC (*20*). Cancers with high MYC levels depend on the Ser/Thr kinase, NUAK1, to promote spliceosome activity, revealing a MYC-sensitive feedback control of transcription (*21*). This is intriguing as *NUAK1* has six NCCMs in five patients, and suggests that *NUAK1* inhibitors (*22*) may be useful for MB patients with high *MYC* or *NUAK1* expression, or low *ZFHX3* expression.

Six NCCMs were found in the *NFIX* locus (Fig. S10A) with seven genes, showing expression in the brain, and specifically the cerebellum (https://www.gtexportal.org/home/). These fall within the same topologically associated domain (TAD), suggesting potential coregulation (Fig. S10B), and data mining showed that *NFIX* expression is higher in SHH MB compared to normal cerebellum (fig. S10C). Because NFIX upregulates EZR in GBM (*23*), and EZR is similarly upregulated in MB where it confers increased aggressiveness (*24*), NCCMs in *NFIX* might alter EZR levels in MB. Furthermore, *DAND5*, a secreted BMP antagonist (*25*) in the same locus, could be tumor-promoting by maintaining low BMP levels. Recreating NCCM2 and NCCM4 of this locus in DAOY MB cells with CRISPR/Cas knock-in editing show differences in expression for multiple genes when compared to the non-edited control cells. *NFIX* is down-regulated by NCCM2, while *NACC1* show increased expression by NCCM4*. LYL1* was up-regulated both by NCCM2 and NCCM4 while *TRMT1* was unchanged (fig. S10D). While reporter assays and proliferation pattern for the edited variants did not reach significance (fig S10E-G), several oncogenic roles have been ascribed to genes of this locus, which merits further investigations of possible phenotypes associated with these NCCMs.

The locus containing *WASF2AHDC1/FGR*, with six NCCMs (Fig. 4A), is highly conserved, and microdeletions/duplications have been linked to neurodevelopmental disorders (*26*). A reporter assay showed that the NCCM2 mutation increased reporter expression close to significant levels (p=0.05, Fig. 4B) in the MB cell line MB002. To investigate the functional potential of NCCM2 in this locus, we used CRISPR/Cas9 to insert NCCM2 in DAOY cells. While expression of *AHDC1* did not markedly change, expression of the neighbouring gene, *FGR*, which shares the same TAD (fig. S11) increased upon gene editing (p=0.02, Fig. 4C). *FGR* is a non-receptor tyrosine-protein kinase of the SRC family, with oncogenic capacity in AML (*27*) but not previously implicated in MB. DAOY cells with NCCM2 showed a significantly increased proliferation rate (Fig. 4D) compared to control cells. Both CRISPR/Cas9-edited and control DAOY cells were growth-arrested by cisplatin, a chemotherapeutic agent used widely as a treatment of MB (*28*) (*29*) while dasatinib, a SRC family kinase inhibitor (*30*), reduced the proliferation of NCCM2-edited DAOY cells more efficiently than controls (Fig. 4E). This suggests that NCCMs could alter drug response of MB cells.

**Figure 4.**
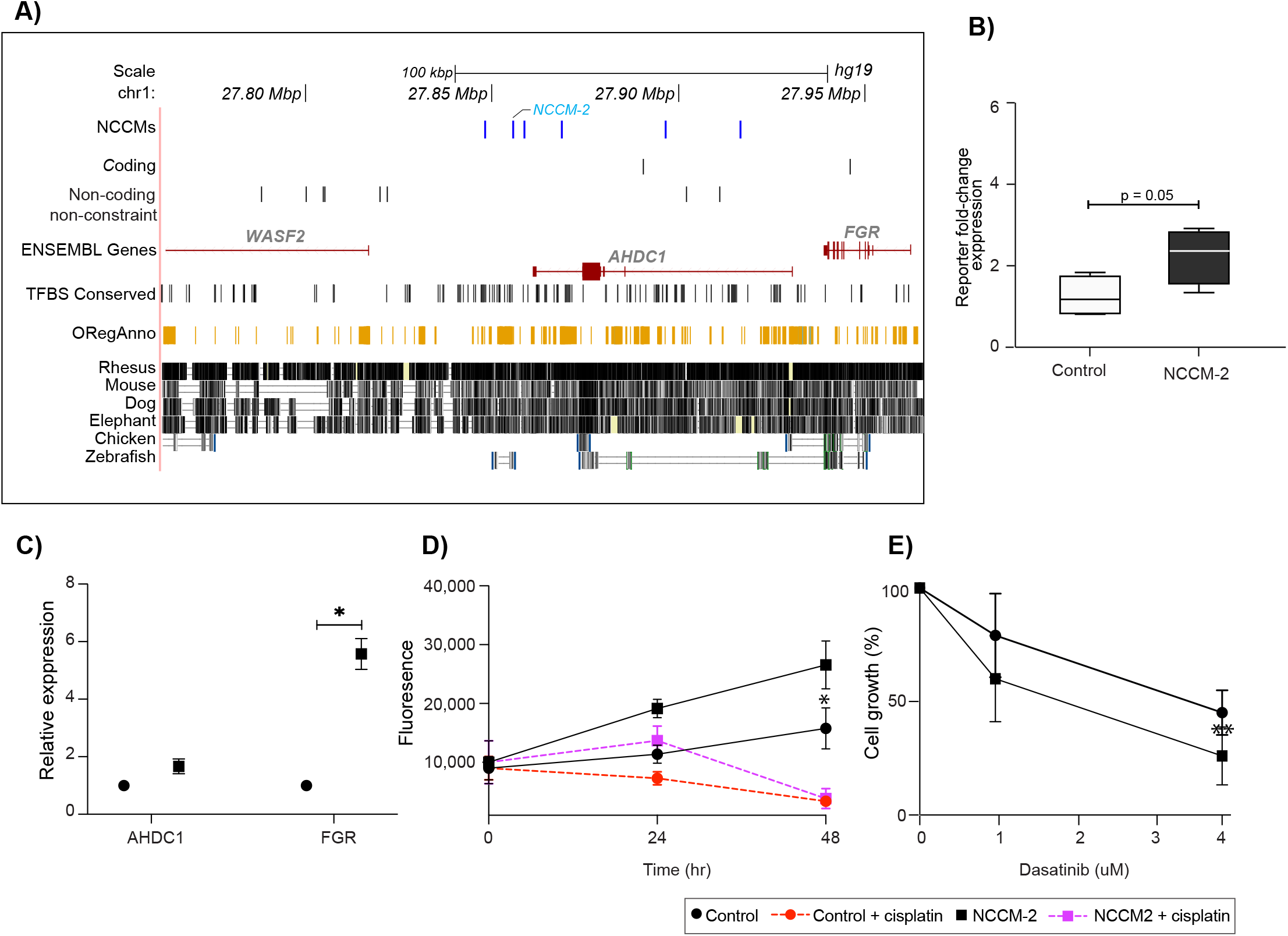
NCCMs in the *ADHC1, WASF2* and *FGR* genes mainly in adult patients. **A)** Six NCCMs (blue), with regulatory potential in the introns of the *ADHC1* gene are close to *WASF2* and *FGR*, and found in areas of high mammalian conservation. **B)** Allelic effects of the *AHDC1* NCCM2 on reporter transcript expression shows near significant (p=0.05) fold change relative to the level of the wild type reporter vector in the MB002 cells. **C)** CRISPR/Cas9 editing of NCCM2 in DAOY cells leads to increased expression of *FGR*. **D)** DAOY cells with NCCM2 show faster growth than control DAOY cells, and both are growth-inhibited by cisplatin (1μg/ml) treatment. **E)** Dasatinib, a SRC kinase inhibitor, more efficiently decrease growth of the NCCM2 CRISPR/Cas9 DAOY cells, compared to control cells after exposure for 72 hours.

In conclusion, using phyloP constraint scores from 240 mammals, we have shown that genes with NCCMs are associated with different age of onset and MB subgroups. Several novel candidate driver genes have plausible roles in MB development and NCCMs could therefore be of diagnostic and/or therapeutic potential.

## Supporting information

Roy et al, Supplementary material

## Acknowledgments

We thank the International Cancer Genome Consortium for controlled access to WGS for PA and MB. The MB002 cell line was a kind gift from Dr. Y-J. Cho, OHSU, Portland, Oregon. Computations were performed on resources provided by the Swedish National Infrastructure for Computing (SNIC) through Uppsala Multidisciplinary Center for Advanced Computational Science (UPPMAX) under project sens2020579.

## Funding

Swedish Childhood Cancer Fund PR2020-0005 (KFN)

Swedish Research Council 2018-02477_VR (KFN) and 541-2013-8161 (KLT)

Swedish Cancer Society PR190206Pj (KFN) and PR200832PjF (KLT)

## Author contributions

Conceptualization: KLT, KFN

Methodology: AR, SS, SVK, RP, MP, KLT, KFN

Investigation: AR, SS, JN, SVK, RP, EKK, KLT, KFN

Visualization: AR, SS, SVK

Funding acquisition: KLT, KFN

Supervision: KLT, KFN

Writing – original draft: KLT, KFN

Writing – review & editing: AR, SS, JN, SVK, RP, KLT, KFN

## Data and materials availability

All data are available in the main text or the supplementary materials.

