## Supplementary material for "Using evolutionary constraint to define novel candidate driver genes in medulloblastoma": Roy et al, Supplementary material

#### Materials and Methods

##### *Cohort information and data access*

All WGS data used in this project are a part of the International Cancer Genome Consortium (ICGC) project PBCA-DE. Access to the controlled data-set for analysis in this project was granted to the authors by the ICGC Data Access Compliance Office (DACO). Somatic mutation files in variant calling format (vcf) corresponding to 89 pilocytic astrocytoma (PA) and 146 medulloblastoma (MB) patient samples were downloaded from the ICGC data portal (https://dcc.icgc.org/releases/PCAWG/consensus_snv_indel/) (*31*) (*6, 7*). The raw sequence data corresponding to these analysis-ready files was aligned to the human genome build hg19. Consequently, all downstream analyses performed on the data use this version of the human genome. The cohort comprised of 47 females and 41 males for the PA set, and the median age of diagnosis was 8 years (range: 1-50). Similarly, among the MB cohort of 66 females and 90 males, the median age of diagnosis was 9 years (range: 1-49). The molecular subgroups for the MB tumors include: WNT ($n$ = 3), SHH ($n$ = 37), Group 3 ($n$ = 43), and Group 4 ($n$ = 60). For 3 MB samples there was no information available for the molecular histological classification.

##### *Molecular classification of Medulloblastoma (MB)*

Molecular profiling divides MB into four subgroups; WNT (Wingless), SHH (Sonic Hedgehog), Group 3, and Group 4 (*32*). The WNT subgroup has the best prognosis of > 90% survival outcome. This group carries germline mutations of *APC*, a WNT pathway inhibitor, as well as somatic mutations in *CTNNB1* and high protein expression of ß-catenin. The SHH sub-group patients have a bimodal age distribution into infant and adolescent groups. These tumors have mutations in the

SHH signaling pathways, including *PTCH*, *SMO*, and *SUFU*, as well as amplifications of *GL1* and *GL2* and have similarities to Group 3 MB which is exclusive to infants and has the worst clinical outcome. These tumors are genetically characterized by *MYC* amplifications, with additional *MYC*, *PVT1*, *OTX2*, *MLL2*, *SMARCA4* and *CHD7* mutations being reported. Group 4 MB can affect both infants and adolescents and display 75% five-year survival rate. Group 4 accounts for almost 35% MB patients and is characterized by isochromosome 17q, loss of chr X, 17p and *MYCN* and *CDK6* amplifications. For both groups 3 and 4 >30% of the patients have metastatic disease at diagnosis.

***Classification of variant calls and identification of significantly mutated genes based on protein-coding mutations***

GATK's Funcotator module (GATK 4.1.4.) was used to functionally classify the MB and PA somatic point mutation (SPM) and somatic indel mutations (SIM) as being either coding or non-coding changes. The coding changes were subsequently inputs to the algorithm MutSigCV1.41 to identify significantly mutated genes (SMG), genes with a statistically higher rate of somatic changes than expected by random chance.

***Identification of non-coding constraint mutations (NCCMs)***

All of the SPM and SIM calls were annotated with phyloP scores to identify NCCMs. The human genome hg38-phyloP scores (a measure of evolutionary constraint, based on the alignment of 240 mammals) were first mapped to the hg19 genome using the UCSC genome browser utility 'liftOver', after which the annotation was applied to the variant calls. The phyloP scores range from a minimum of -20 to a maximum of +9, corresponding to regions of accelerated evolution to highly conserved regions in the genome. Prior estimates of the fraction of the human genome that is functional have been between 6-13%. Hence, we used a conservative estimate of the top 8%

constraint positions previously used by us to examine NCCMs in glioblastoma (*4*). This corresponds to phyloP values ≥1.2. The non-coding variants that met the selection criteria for constraint, henceforth referred to as non-coding constraint mutations or NCCMs were then chosen for additional study. First, NCCMs located within 3′- or 5′-UTRs, introns, or ±100 *kbp* intergenic flanking regions of protein-coding genes were extracted. Next, the rate of NCCMs, normalized to the length of the queried genomic region around each gene was computed, and those genes that had ≥2 NCCMs per 100 *kbp* were delineated as candidate driver genes (*n*=114). We also normalized the NCCM rate per patient and still found a higher rate of NCCMs in MB. In addition, we identified all genes in MB with ≥5 NCCMs in the target area (±100 *kbp*) for further analysis (*n*=525). The union of the two data sets was *n*=530 genes.

***Comparison of NCCM rates between MB and PA***

To compare the NCCM rates in the MB and PA cohort, the rate for each gene was calculated as described above, and the resulting rate was then subsequently divided by the number of patients in the respective cohort. The NCCM rates of all genes in the genome, as well as the top 0.5 % of genes in each cohort were compared between MB and PA using unpaired Student's t-tests.

***NCCM rate correlation analysis between MB subgroups***

The NCCM rate was calculated separately for MB patients of the SHH, Group 3 and Group 4 subtypes as well as for paediatric (< 18 years) and adult (≥ 18 years) patients. To determine whether the NCCM rate around certain genes could be used to separate patient subgroups, the subgroup-specific NCCM rates of the 114 candidate genes were further correlated pairwise using the Kendall rank correlation coefficient.

***Evaluation of the putative functional impact of the NCCMs***

To verify if the NCCMs surrounding the candidate genes have regulatory changes, their coordinates were annotated with regulatory annotations corresponding to the hg19 reference assembly, downloaded from either the UCSC genome browser (Genome Reference Consortium GRCh37 version); or ENCODE portal (v2, source data version: ENCODE Jan 2011), or both. These included, among others, information from tracks of transcription factor binding sites (TFBS), regulatory markers (ORegAnno), transcription start sites (TSS), enhancer information, and DNase I hypersensitive sites (DHS). The topological associated domains (TADs) were visualised with Hi-C data on the 3D genome browser. For the visualization of the *NFIX* region a TAD reaching from approximately 13.0 Mb to 13.8 Mb using raw tissue Fetal_Brain_CP and Fetal_Brain_GZ at a 10 kb resolution, and Cortex_DLPFC at a 40 kb resolution was applied.

The tool sTRAP (*33*) was used to predict if the NCCMs were likely to alter the binding affinity of transcription factors to mutated versus wild-type sequences. For every NCCM allele and the corresponding wild-type, 20 *bp* upstream and downstream were used as input sequences. The matrices for the analysis were set to the JASPAR vertebrates database (*34*), and for the background model, the 'human promoter' option was selected. The JASPAR database (http://jaspar.genereg.net) was used to obtain information about transcription factor (TF) binding matrices and motifs of interest. Affinity profiles for the top-ranking matrices with $p$-values $\leq 0.05$ were recorded as having significantly more TF binding for the input sequences than what could be expected from a random sequence.

***Pathway analysis for candidate genes***

To gain mechanistic insights for the candidate genes, enrichment analysis was performed using the tool g:GOSt from the online utility g:Profiler (https://biit.cs.ut.ee/gprofiler/gost. First, the options for gene ontology analysis were set to GO- 'molecular function', 'cellular component', and 'biological process'. Then, for the pathway analysis, the data sources of KEGG and Reactome pathway databases were selected. Finally, the functional term with enrichment of *p*-values ≤0.5, the total number of genes for the term, and the overlap from the query set were tabulated.

***Data visualization***

To visualize the somatic alterations, oncoprints (brick plots) were generated using tools available on the web-based utility cBioPortal (*35*). Histograms were plotted using either R or Matlab plotting tools. Custom UCSC genome-browser views were generated by building NCCM tracks for a specified genomic locus, followed by selection of annotation tracks that included TFBS and OregAnno (from 'Regulation' track), and 'Multiz' alignments of 100 vertebrates. These specific track combinations, were saved in a session and reproduced as needed.

***Cell cultures***

The MB cell line MB002 representing Group 3 (*36*) was a kind gifts from Dr. Yoon-Jae Cho (Dept. of Neurology and Neurological Science, Stanford University, USA) and was cultured in 1:1 mixture of Neurobasal without vitamin A (Life Technologies, Stockholm, Sweden) and DMEM/F12 (Life Technologies) supplemented with 1% non-essential amino acids (Life Technologies), 1 mM sodium pyruvate (Life Technologies), 250 mM Hepes (Life Technologies), 1% glutaMAX, B27 (Life Technologies), heparin (Stemcell Technologies, Grenoble, France), leukemia inhibitory factor (Merck Millipore, Billerica, MA, USA), 20 ng/mL of fibroblast growth factor 2 (FGF2) and 20 ng/mL epidermal growth factor (EGF). The MB cell line DAOY representing the SHH Group was grown in Dulbecco's Modified Eagle Medium supplemented with 10% fetal bovine serum (FBS).

***Electrophoretic mobility shift assay (EMSA)***

EMSA was run as per the established protocol described previously (*35*). Briefly, oligos (100 μM) were first annealed in equimolar amounts in 1× annealing buffer (50 mM NaCl, 10 mM Tris-HCl, 10 mM MgCl2, 100 μg/ml BSA, pH 7.9 at 25 °C) in a thermocycler by heating to 95 °C for 5 min and gradual cooling at 1 °C/min to 4 °C. Nuclear proteins were extracted from the medulloblastoma cell line MB002 using the NucBuster™ Protein Extraction Kit (Millipore). The subsequent binding reaction of the dsDNA with nuclear protein extract was processed using LightShift™ Chemiluminescent EMSA Kit (Thermo Scientific) as per the manufacturer's protocol, and the binding reaction was conducted by incubation of the dsDNA-nuclear protein extract on ice for 40 min. The reaction was stopped by the addition of 5 μl of loading buffer to each binding reaction. A total of 20 μl per reaction was then immediately loaded per well onto a Bio-Rad Criterion gel (Bio-Rad) and electrophoresed for 90 min at 200 V. The gel was transferred to a GeneScreen Plus nylon hybridization transfer membrane (PerkinElmer) for 1 h at 45 V followed by UV crosslinking for 15 min with the membrane facing down on a transilluminator and additional 1 more minute with the membrane turned over. The membrane was developed using the Chemiluminescent Nucleic Acid Detection Module Kit (Thermo Scientific) and visualized on the Bio-Rad CCD camera (Bio-Rad). All the sequence probes were designed using the GRCh37/hg19 assembly, and the observed mutations for the NCCMs were inserted in the oligo sequence as required (supplementary table S14). All oligos were ordered in sets of three for each wt/NCCM allele pair from Integrated DNA Technologies, with one 5′ biotin-labeled and the same as unlabeled forward strand DNA oligo and one reverse complementary unlabeled strand (HPLC-purified).

***Reporter assays***

gBlock DNA fragments (IDT) of about 250 bp centered on NCCM and including either reference or mutant allele were cloned into a modified pGL4.26 vector (Promega) with a 9-bp barcode sequence inserted before the poly(A) site. The plasmids were transformed into TOP10 competent cells (Invitrogen), and individual clones were validated by Sanger sequencing. The selected plasmids were pooled together and with a control vector without insert, re-transformed and purified as a pool with EndoFree Plasmid Maxi Kit (Qiagen). Transfection of the cells (MB002) was performed using plasmids pooled together (reporter constructs and control empty vector) using Lipofectamine 3000 according to the manufacturer's protocol. Cells were harvested 48 h post transfection in TRIZOL (Thermo Fisher Scientific) and total RNA and DNA were purified according to the manufacturer's protocol. One µg of RNA/sample was treated with RQ1 RNase-free DNase (Promega) and converted into cDNA using 1 U of M-MuLV reverse transcriptase, oligo(dT) primers, 1 mM dNTPs and RNase inhibitor (all from Thermo Fisher Scientific). Reporter transcript levels of each allelic construct and empty vector were measured by quantitative real-time reverse transcription PCR (qRT-PCR) using cDNA, and independently the same fragments were analysed by qPCR on genomic DNA. The forward primer was common for all plasmids, and reverse primers were specific for plasmid barcodes, generating amplicons of 124 bp that were detected using SYBR green (Thermo Fisher Scientific) by QuantStudio 6 Flex (Applied Biosystems) real-time PCR system. Reporter transcript levels were normalized to the levels of the plasmid DNA using the standard curve method. The experiment was performed three times with five replicates per plasmid pool per experiment. Statistical analyses of all transfections were performed using unpaired Student's t-test.

***CRISPR/Cas9 gene editing***

CRISPR/Cas9 SNV knock-in cell pools were produced by Synthego (https://www.synthego.com/help/advanced-cell-projects) as per their standardized protocol. In brief, the homology of the sequence ~1 kb of the targeted genomic region was determined for the target cells (DAOY). Multiple chemically modified synthetic single guide RNA (sgRNA) with different PAMs were designed. The donor sequences were selected to be between 115-250 bp long. Cells were transfected with ribonucleoprotein (RNPs) complexes consisting of sgRNA and spCas9 protein through nucleofection and knock-in efficiency was analysed using the free online tool ICE (Inference of CRISPR edits) following Sanger sequencing of the region of interest. The sgRNAs that had the highest knock-in efficiency were chosen for further expansion and cell pools for the various SNVs with the best sgRNA was subsequently used for the experiments. Positive control sgRNA (RELA) were always transfected at the same time and edit efficiency evaluated. The negative controls (mock transfected with Cas9 protein only) were used as WT or experimental control cells as matched compare for the downstream experiments.

***Expression and survival analysis using public databases***

To compare gene expression levels between normal cerebellum and MB or between the MB subgroups, processed gene expression data was downloaded from the Gene Expression Omnibus (GEO) database repository. The dataset (GSE124814) used for expression analysis is a batch-normalized resource comprising transcription profiles of 1350 MB samples and 291 normal cerebellum samples (*15*). All statistical analysis was performed on GraphPad software Prism version 9.1.1 (a commercial proprietary scientific 2D graphing and statistics software, San Diego, CA, USA). The significance of differences in the gene expression between individual subgroups was determined using Student's unpaired *t*-test with Welch's correction.

The expression dataset (GSE85217) containing 763 MB samples, including 70 WNT, 223 SHH, 144 Group 3, and 326 Group 4 cases (*10*), was used for survival analysis. The Kaplan-Meier survival analysis was performed on the R2 (http://r2.amc.nl) genomics platform. Briefly, the dataset samples were divided into two groups based on the gene expression for a given gene. In the order of expression, every increasing expression value is used as a cut-off to create the two groups, and the significance was tested using a log-rank test. The most significant expression cut-off for survival analysis was plotted on the Kaplan Meier curve based on the log-rank test.

##### *Expression analysis*

RNA was extracted from cultured cells using the RNeasy kit from Qiagen. 200 ng of RNA was used for cDNA synthesis using High-Capacity RNA-to-cDNA™ Kit (ThermoFisher). Quantitative PCR was performed using SYBR Green master mix (Applied Biosystems, Foster City, CA, USA) on a StepOnePlus real time PCR system (Applied Biosystems). Samples were amplified in triplicate and data analyzed using the ΔΔCT method. All statistical analysis was performed and graphs plotted on GraphPad software Prism version 9.1.1 (a commercial proprietary scientific 2D graphing and statistics software, San Diego, CA, USA).

##### *Proliferation assay and drug treatment of MB cells*

Cell proliferation was assessed in 96 well format for 0, 24, 48 and 72 hours using the CyQuant Direct Cell Proliferation Kit (ThermoFisher). Percentage of proliferation compared to the 0 hr-reading was calculated and plotted. For treatments, cisplatin (1μg/ml) and dasatinib (1-4μM) were administered at the beginning of the assay and cell numbers monitored as mentioned above. All *in vitro* experiments were statistically analysed and plotted using the GraphPad software Prism.

#### SUPPLEMENTARY FIGURE LEGENDS

**Figure S1**

**Mutational counts for MB are higher than what is observed for PA**. **A)** Boxplot depicting the total number of mutations in MB versus PA. Median, the middle data point is represented as a line in the middle of the boxplot and the upper whiskers represent the maximum value within 1.5 * interquartile range of the upper quartile. Plus signs denote average mutational burden per cohort. Un-paired student t-test was used to analyse the cohorts and the p values for the statistical differences are indicated in the figure.

**Figure S2**

**Genes in the *BRAF* locus harbor non-coding constraint mutations that may have driver roles in PA. A)** Eleven genes with NCCMs ≥ 1 per 100 *kbp* were observed, with three of them in the *BRAF* locus. *BRAF* is known to have putative driver functions in PA. **B)** A UCSC genome browser view shows that the top gene *NDUFB2* shares NCCMs with the genes *BRAF* and *ADCK2*. NCCMs are shown in dark blue, genes in red, transcription factor binding sites in black, and ORegAnno in orange. Lastly, the MultiZ track, which displays a measure of evolutionary conservation, shows that the *BRAF* locus NCCMs are found in regions of high mammalian conservation. **C)** Oncoplot shows the somatic changes in the *BRAF* locus and in key protein-coding genes for the PA cohort. (Key genes are genes that have been previously implicated in PA: *BRAF*, *FGFR1*, *NF1*, *KRAS*, *AHNAK2*, *AMBP*, *FLG*, *IL4R*, *KIAA1549*, *PTPN11*, and *SETD2*).

**Figure S3**

**NCCM raw counts versus normalized for length. A)** Rates of non-coding constraint mutation for MB genes. Altogether 114 genes had ≥2.0 NCCMs/ 100*kbp*. **B)** Number of non-coding constraint mutation per gene in MB samples. Altogether 525 genes had ≥ 5 NCCMs within ±100 *kbp*. **B)** Venn diagram showing overlap of data sets in **A)** and **B)**. The union is 530 genes.

**Figure S4**

**Cohort-wide normalization of NCCMs showed that MB had greater NCCM accumulation rates than PA. A)** The NCCM rate per patient of all the genes in the genome and, **B)** The top 0.5 % of these genes with the highest NCCM rates in each cohort were compared between MB and PA. Unpaired Student t-test was used to analyse the cohorts, and the p values for the statistical difference is indicated in the figure. (*** = p << 0.0001).

**Figure S5**
**Distribution of NCCMs among the molecular subgroups**. **A)** Distribution of NCCMs per molecular subgroup. **B)** Distribution of NCCMs per patient per molecular subgroup.

**Figure S6**
**Candidate driver genes for MB (all 114 genes ≥ 2 NCCMs per 100 *kbp*) are differently distributed across pediatric and adult MB.** Orange, median age <18, purple, median age >18.

**Figure S7**
**Correlation between NCCM rates in subgroups and per age distribution of patients**
Correlation between NCCMs/100 *kbp* per patient in genes with ≥2 NCCMs/100 *kbp* for each combination of subgroups, and between age group. SHH Group and group 3 were highly correlated while young and old groups were negatively correlated.

**Figure S8**
**Oncoplots for MB patients**
Oncoplot of genes with ≥2 NCCMs/100 *kbp* that are mainly found in pediatric MB patients. Blue box denotes a patient with NCCM(s).

**Figure S9**

**NCCMs predominantly found in adult patients**

Oncoplot of somatic changes observed among the seven loci with NCCMs mainly found in adults, >50% of patients have ≥1 NCCM. Blue box denotes an NCCM.

**Figure S10**

**NCCMs in the *NFIX* locus in adult patients**

**A)** In the *NFIX* locus six NCCMs (blue) are shared across seven candidate genes. **B)** Topologically associated domains for the NFIX gene cluster in fetal brain. **C)** Expression of *NFIX* is higher in WNT and SHH MB subgroups, compared to normal cerebellum. Boxes min-max range, horizontal lines median value, and whiskers extend to extreme values. The significance of differences in the gene expression between individual subgroups was determined using Student's unpaired *t*-test with Welch's correction. **D)** CRISPR/Cas9-editing of NCCM2 and NCCM4 in DAOY cells results in decreased expression of *NFIX* for NCCM2, and increased expression of *NACC1* by NCCM4, while *LYL1* was up-regulated both by NCCM2 and NCCM4. **E)** Allelic effects of NCCM2 and NCCM4 on reporter transcript expression did not reach a significant difference (p=0.53, p=0.15) relative to the levels of the wild type reporter vector in the MB002 cells. **F)** Proliferation assay of edited NCCM2, NCCM4 and control DAOY cells did not reach a significant difference.

**Figure S111**

**Topologically associated domains for the *AHDC1* gene cluster in fetal brain.**

**Zoonomia Consortium of collaborators**

Gregory Andrews[1], Joel C. Armstrong[2], Matteo Bianchi[3], Bruce W. Birren[4], Kevin R. Bredemeyer[5], Ana M. Breit[6], Matthew J. Christmas[3], Hiram Clawson[2], Joana Damas[7], Federica Di Palma[8,9], Mark Diekhans[2], Michael X. Dong[3], Eduardo Eizirik[10], Kaili Fan[1], Cornelia Fanter[11], Nicole M. Foley[5], Karin Forsberg-Nilsson[12,13], Carlos J. Garcia[14], John Gatesy[15], Steven Gazal[16], Diane P. Genereux[4], Linda Goodman[17], Jenna Grimshaw[14], Michaela K. Halsey[14], Andrew J. Harris[5], Glenn Hickey[18], Michael Hiller[19,20,21], Allyson G. Hindle[11], Robert M. Hubley[22], Graham M. Hughes[23], Jeremy Johnson[4], David Juan[24], Irene M. Kaplow[25,26], Elinor K. Karlsson[1,4,27], Kathleen C. Keough[17,28,29], Bogdan Kirilenko[19,20,21], Klaus-Peter Koepfli[30,31,32], Jennifer M. Korstian[14], Amanda Kowalczyk[25,26], Sergey V. Kozyrev[3], Alyssa J. Lawler[4,26,33], Colleen Lawless[23], Thomas Lehmann[34], Danielle L. Levesque[6], Harris A. Lewin[7,35,36], Xue Li[1,4,37], Abigail Lind[28,29], Kerstin Lindblad-Toh[3,4], Ava Mackay-Smith[38], Voichita D. Marinescu[3], Tomas Marques-Bonet[39,40,41,42], Victor C. Mason[43], Jennifer R. S. Meadows[3], Wynn K. Meyer[44], Jill E. Moore[1], Lucas R. Moreira[1,4], Diana D. Moreno-Santillan[14], Kathleen M.

Morrill[1,4,37], Gerard Muntané[24], William J. Murphy[5], Arcadi Navarro[39,41,45,46], Martin Nweeia[47,48,49,50], Sylvia Ortmann[51], Austin Osmanski[14], Benedict Paten[2], Nicole S. Paulat[14], Andreas R. Pfenning[25,26], BaDoi N. Phan[25,26,52], Katherine S. Pollard[28,29,53], Henry E. Pratt[1], David A. Ray[14], Steven K. Reilly[38], Jeb R. Rosen[22], Irina Ruf[54], Louise Ryan[23], Oliver A. Ryder[55,56], Pardis C. Sabeti[4,57,58], Daniel E. Schäffer[25], Aitor Serres[24], Beth Shapiro[59,60], Arian F. A. Smit[22], Mark Springer[61], Chaitanya Srinivasan[25], Cynthia Steiner[55], Jessica M. Storer[22], Kevin A. M. Sullivan[14], Patrick F. Sullivan[62,63], Elisabeth Sundström[3], Megan A. Supple[59], Ross Swofford[4], Joy-El Talbot[64], Emma Teeling[23], Jason Turner-Maier[4], Alejandro Valenzuela[24], Franziska Wagner[65], Ola Wallerman[3], Chao Wang[3], Juehan Wang[16], Zhiping Weng[1], Aryn P. Wilder[55], Morgan E. Wirthlin[25,26,66], James R. Xue[4,57], Xiaomeng Zhang[4,25,26]

Affiliations:
[1]Program in Bioinformatics and Integrative Biology, UMass Chan Medical School; Worcester, MA 01605, USA.
[2]Genomics Institute, University of California Santa Cruz; Santa Cruz, CA 95064, USA.
[3]Department of Medical Biochemistry and Microbiology, Science for Life Laboratory, Uppsala University; Uppsala, 751 32, Sweden.
[4]Broad Institute of MIT and Harvard; Cambridge, MA 02139, USA.
[5]Veterinary Integrative Biosciences, Texas A&M University; College Station, TX 77843, USA.
[6]School of Biology and Ecology, University of Maine; Orono, ME 04469, USA.
[7]The Genome Center, University of California Davis; Davis, CA 95616, USA.
[8]Genome British Columbia; Vancouver, BC, Canada.
[9]School of Biological Sciences, University of East Anglia; Norwich, UK.
[10]School of Health and Life Sciences, Pontifical Catholic University of Rio Grande do Sul; Porto Alegre, 90619-900, Brazil.
[11]School of Life Sciences, University of Nevada Las Vegas; Las Vegas, NV 89154, USA.
[12]Department of Immunology, Genetics and Pathology, Science for Life Laboratory, Uppsala University; Uppsala, 751 85, Sweden.
[13]Biodiscovery Institute, University of Nottingham; Nottingham, UK.
[14]Department of Biological Sciences, Texas Tech University; Lubbock, TX 79409, USA.
[15]Division of Vertebrate Zoology, American Museum of Natural History; New York, NY 10024, USA.
[16]Keck School of Medicine, University of Southern California; Los Angeles, CA 90033, USA.
[17]Fauna Bio Incorporated; Emeryville, CA 94608, USA.
[18]Baskin School of Engineering, University of California Santa Cruz; Santa Cruz, CA 95064, USA.
[19]Faculty of Biosciences, Goethe-University; 60438 Frankfurt, Germany.
[20]LOEWE Centre for Translational Biodiversity Genomics; 60325 Frankfurt, Germany.
[21]Senckenberg Research Institute; 60325 Frankfurt, Germany.
[22]Institute for Systems Biology; Seattle, WA 98109, USA.
[23]School of Biology and Environmental Science, University College Dublin; Belfield, Dublin 4, Ireland.
[24]Department of Experimental and Health Sciences, Institute of Evolutionary Biology (UPF-CSIC), Universitat Pompeu Fabra; Barcelona, 08003, Spain.
[25]Department of Computational Biology, School of Computer Science, Carnegie Mellon University; Pittsburgh, PA 15213, USA.
[26]Neuroscience Institute, Carnegie Mellon University; Pittsburgh, PA 15213, USA.
[27]Program in Molecular Medicine, UMass Chan Medical School; Worcester, MA 01605, USA.

[28]Department of Epidemiology & Biostatistics, University of California San Francisco; San Francisco, CA 94158, USA.
[29]Gladstone Institutes; San Francisco, CA 94158, USA.
[30]Center for Species Survival, Smithsonian's National Zoo and Conservation Biology Institute; Washington, DC 20008, USA.
[31]Computer Technologies Laboratory, ITMO University; St. Petersburg 197101, Russia.
[32]Smithsonian-Mason School of Conservation, George Mason University; Front Royal, VA 22630, USA.
[33]Department of Biological Sciences, Mellon College of Science, Carnegie Mellon University; Pittsburgh, PA 15213, USA.
[34]Senckenberg Research Institute and Natural History Museum Frankfurt; 60325 Frankfurt am Main, Germany.
[35]Department of Evolution and Ecology, University of California Davis; Davis, CA 95616, USA.
[36]John Muir Institute for the Environment, University of California Davis; Davis, CA 95616, USA.
[37]Morningside Graduate School of Biomedical Sciences, UMass Chan Medical School; Worcester, MA 01605, USA.
[38]Department of Genetics, Yale School of Medicine; New Haven, CT 06510, USA.
[39]Catalan Institution of Research and Advanced Studies (ICREA); Barcelona, 08010, Spain.
[40]CNAG-CRG, Centre for Genomic Regulation, Barcelona Institute of Science and Technology (BIST); Barcelona, 08036, Spain.
[41]Department of Medicine and LIfe Sciences, Institute of Evolutionary Biology (UPF-CSIC), Universitat Pompeu Fabra; Barcelona, 08003, Spain.
[42]Institut Català de Paleontologia Miquel Crusafont, Universitat Autònoma de Barcelona; 08193, Cerdanyola del Vallès, Barcelona, Spain.
[43]Institute of Cell Biology, University of Bern; 3012, Bern, Switzerland.
[44]Department of Biological Sciences, Lehigh University; Bethlehem, PA 18015, USA.
[45]BarcelonaBeta Brain Research Center, Pasqual Maragall Foundation; Barcelona, 08005, Spain.
[46]CRG, Centre for Genomic Regulation, Barcelona Institute of Science and Technology (BIST); Barcelona, 08003, Spain.
[47]Department of Comprehensive Care, School of Dental Medicine, Case Western Reserve University; Cleveland, OH 44106, USA.
[48]Department of Vertebrate Zoology, Canadian Museum of Nature; Ottawa, Ontario K2P 2R1, Canada.
[49]Department of Vertebrate Zoology, Smithsonian Institution; Washington, DC 20002, USA.
[50]Narwhal Genome Initiative, Department of Restorative Dentistry and Biomaterials Sciences, Harvard School of Dental Medicine; Boston, MA 02115, USA.
[51]Department of Evolutionary Ecology, Leibniz Institute for Zoo and Wildlife Research; 10315 Berlin, Germany.
[52]Medical Scientist Training Program, University of Pittsburgh School of Medicine; Pittsburgh, PA 15261, USA.
[53]Chan Zuckerberg Biohub; San Francisco, CA 94158, USA.
[54]Division of Messel Research and Mammalogy, Senckenberg Research Institute and Natural History Museum Frankfurt; 60325 Frankfurt am Main, Germany.
[55]Conservation Genetics, San Diego Zoo Wildlife Alliance; Escondido, CA 92027, USA.
[56]Department of Evolution, Behavior and Ecology, School of Biological Sciences, University of California San Diego; La Jolla, CA 92039, USA.
[57]Department of Organismic and Evolutionary Biology, Harvard University; Cambridge, MA

02138, USA.
[58]Howard Hughes Medical Institute; Chevy Chase, MD, USA.
[59]Department of Ecology and Evolutionary Biology, University of California Santa Cruz; Santa Cruz, CA 95064, USA.
[60]Howard Hughes Medical Institute, University of California Santa Cruz; Santa Cruz, CA 95064, USA.
[61]Department of Evolution, Ecology and Organismal Biology, University of California Riverside; Riverside, CA 92521, USA.
[62]Department of Genetics, University of North Carolina Medical School; Chapel Hill, NC 27599, USA.
[63]Department of Medical Epidemiology and Biostatistics, Karolinska Institutet; Stockholm, Sweden.
[64]Iris Data Solutions, LLC; Orono, ME 04473, USA.
[65]Museum of Zoology, Senckenberg Natural History Collections Dresden; 01109 Dresden, Germany.
[66]Allen Institute for Brain Science; Seattle, WA 98109, USA

### Supplementary Figure 1

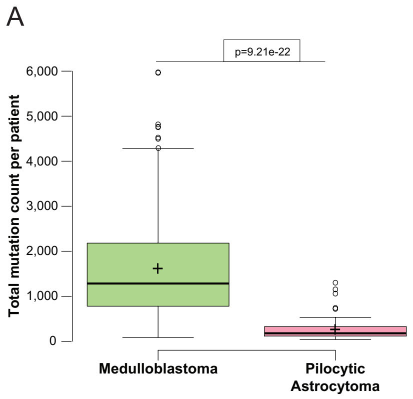

### Supplementary Figure 2

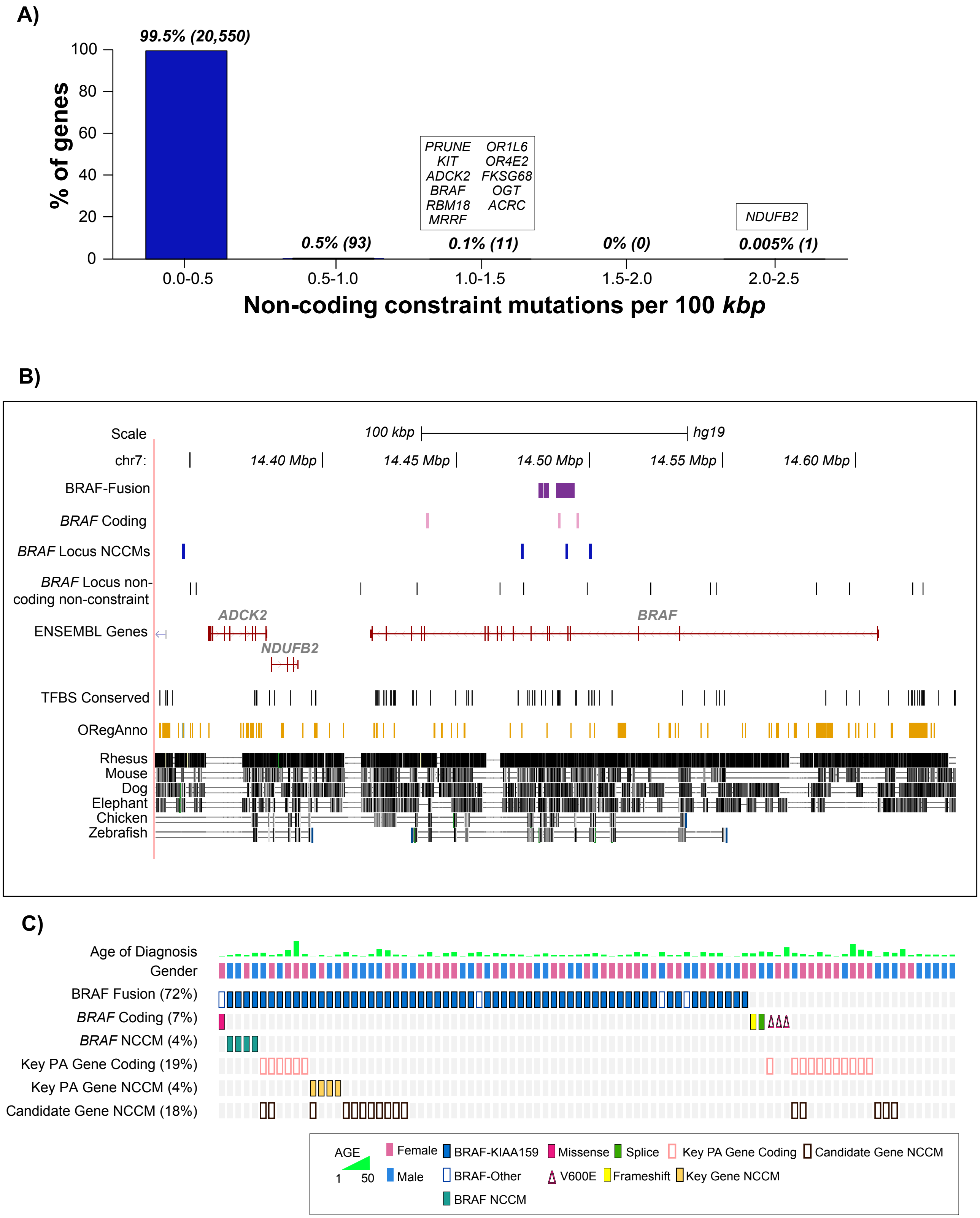

### Supplementary Figure 3

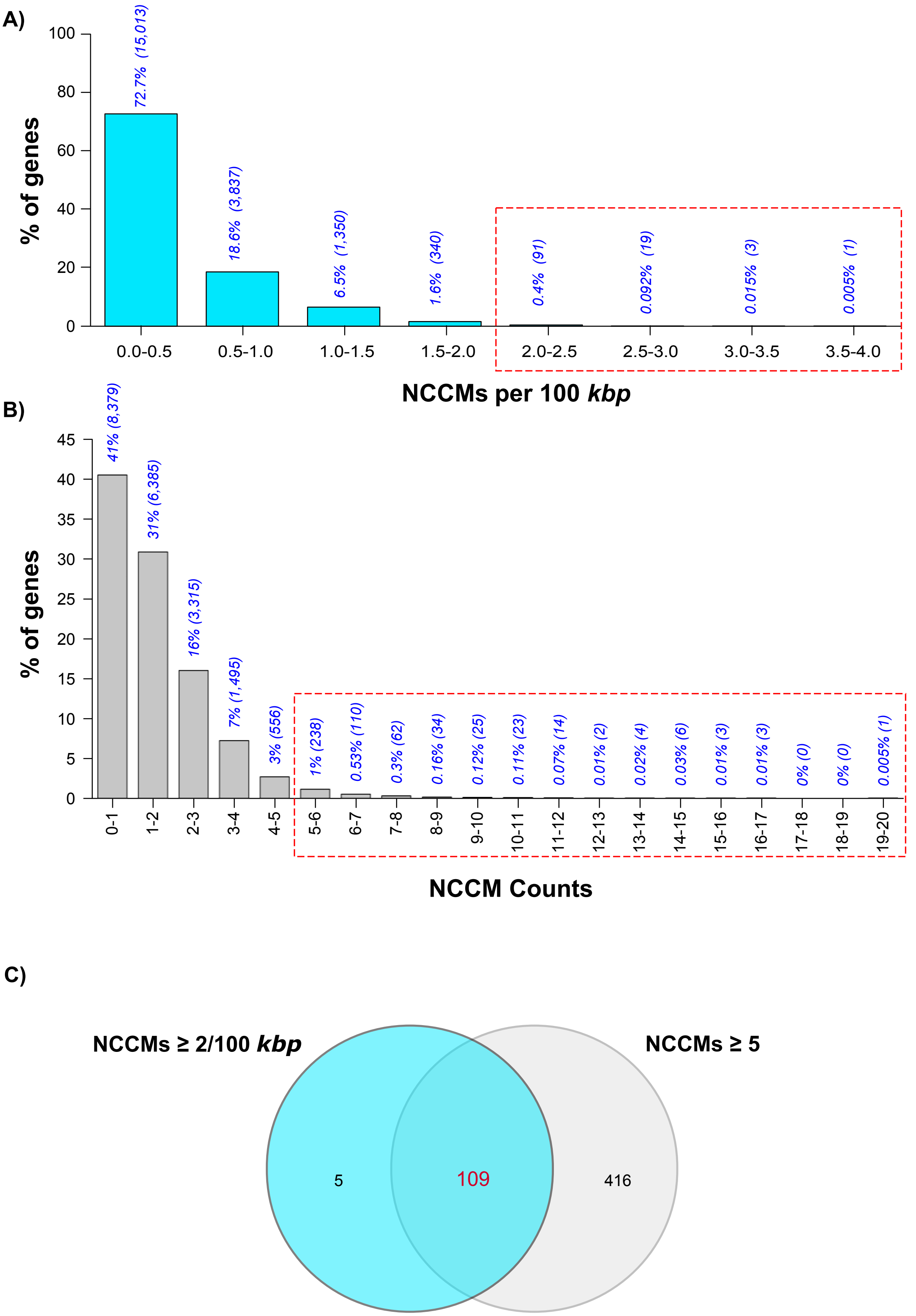

### Supplementary Figure 4

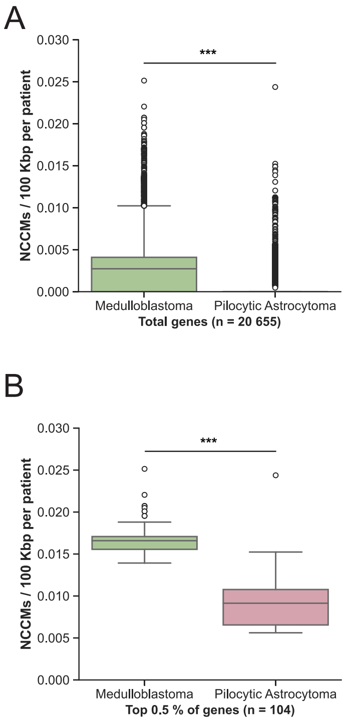

### Supplementary Figure 5

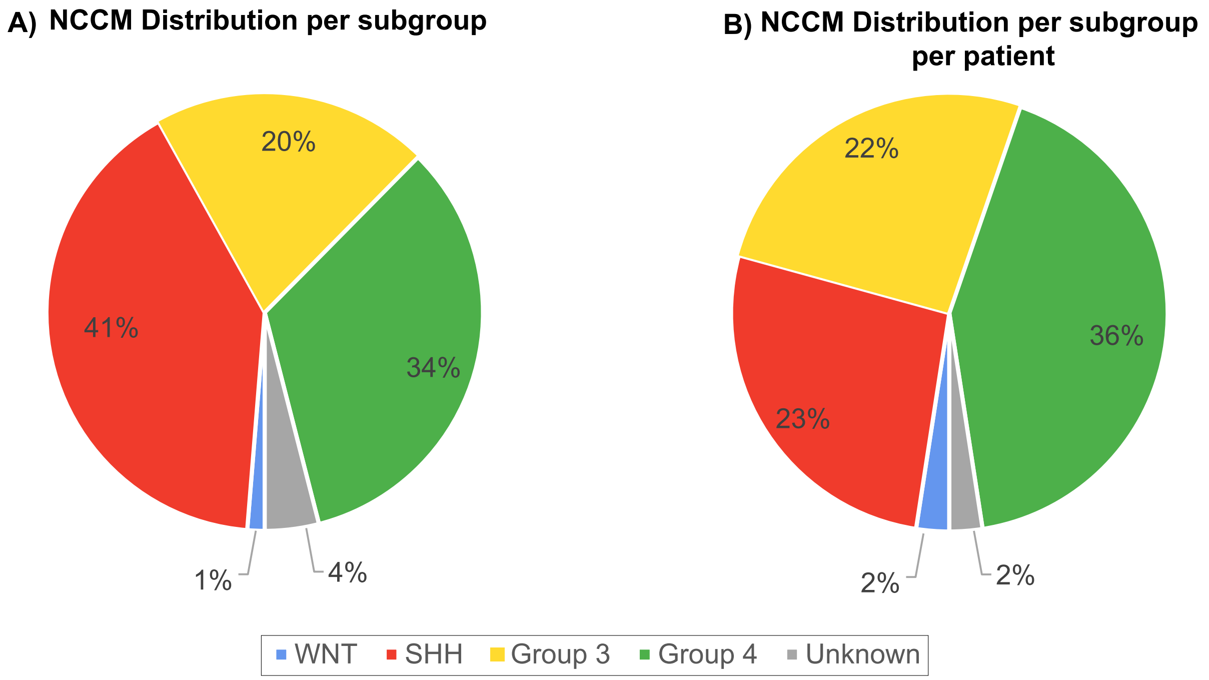

### Supplementary Figure 6

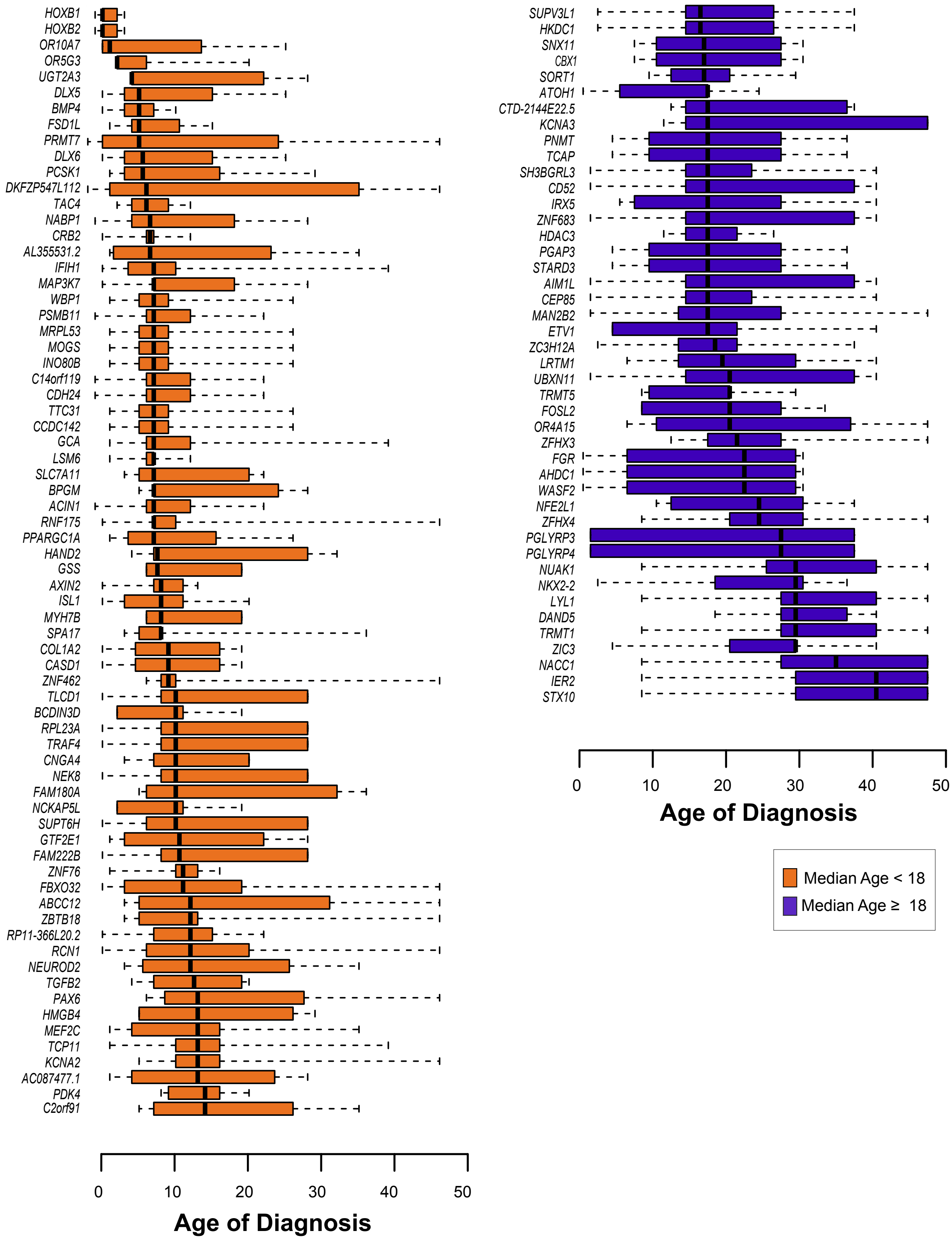

### Supplementary Figure 7

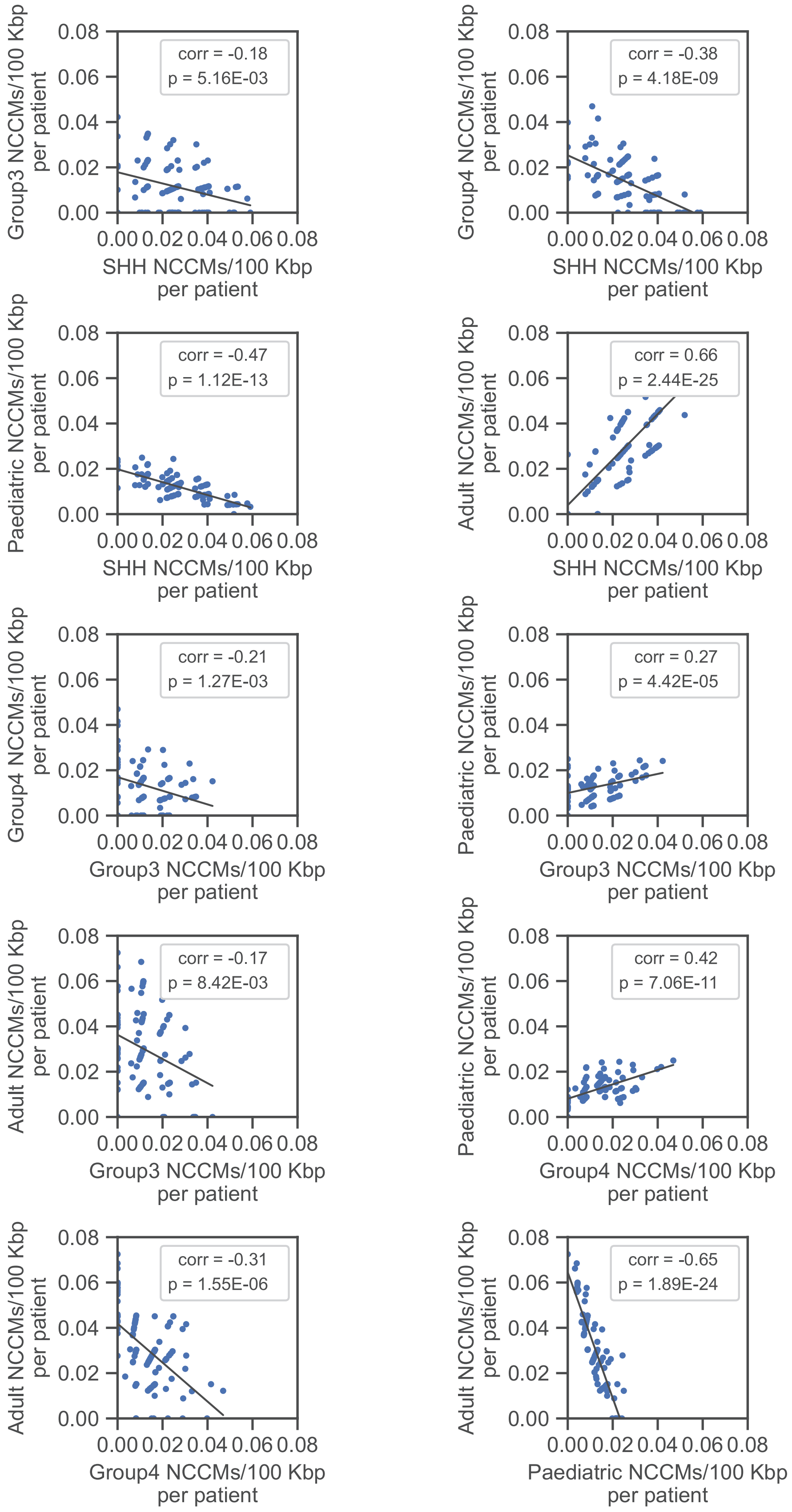

### Supplementary Figure 8

A)

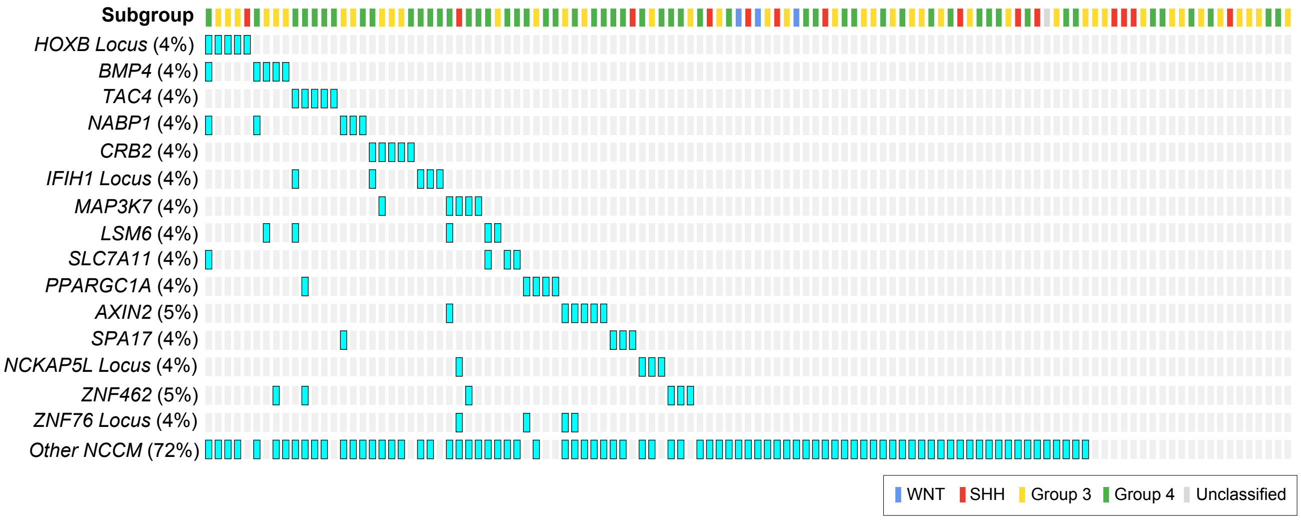

### Supplementary Figure 9

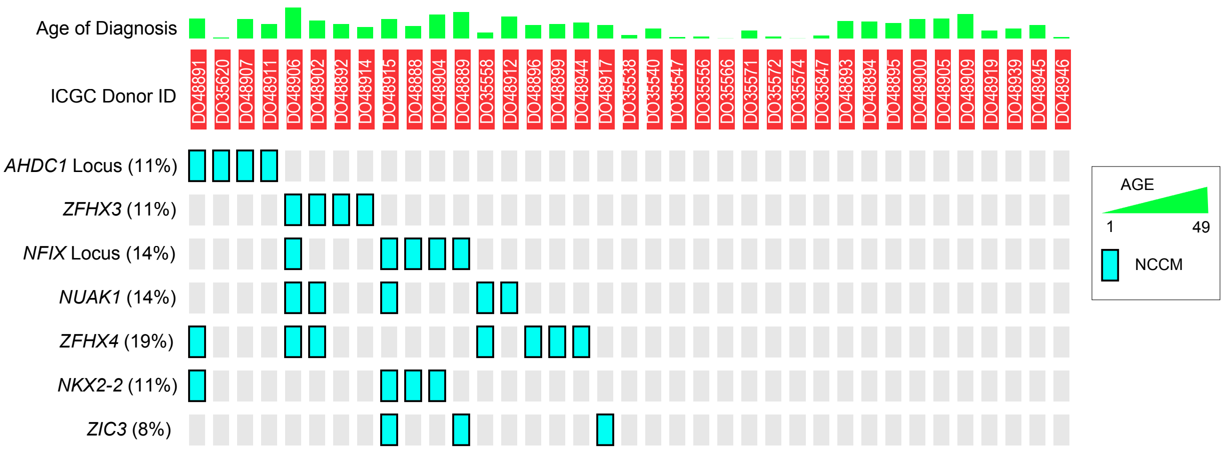

### Supplementary Figure 10

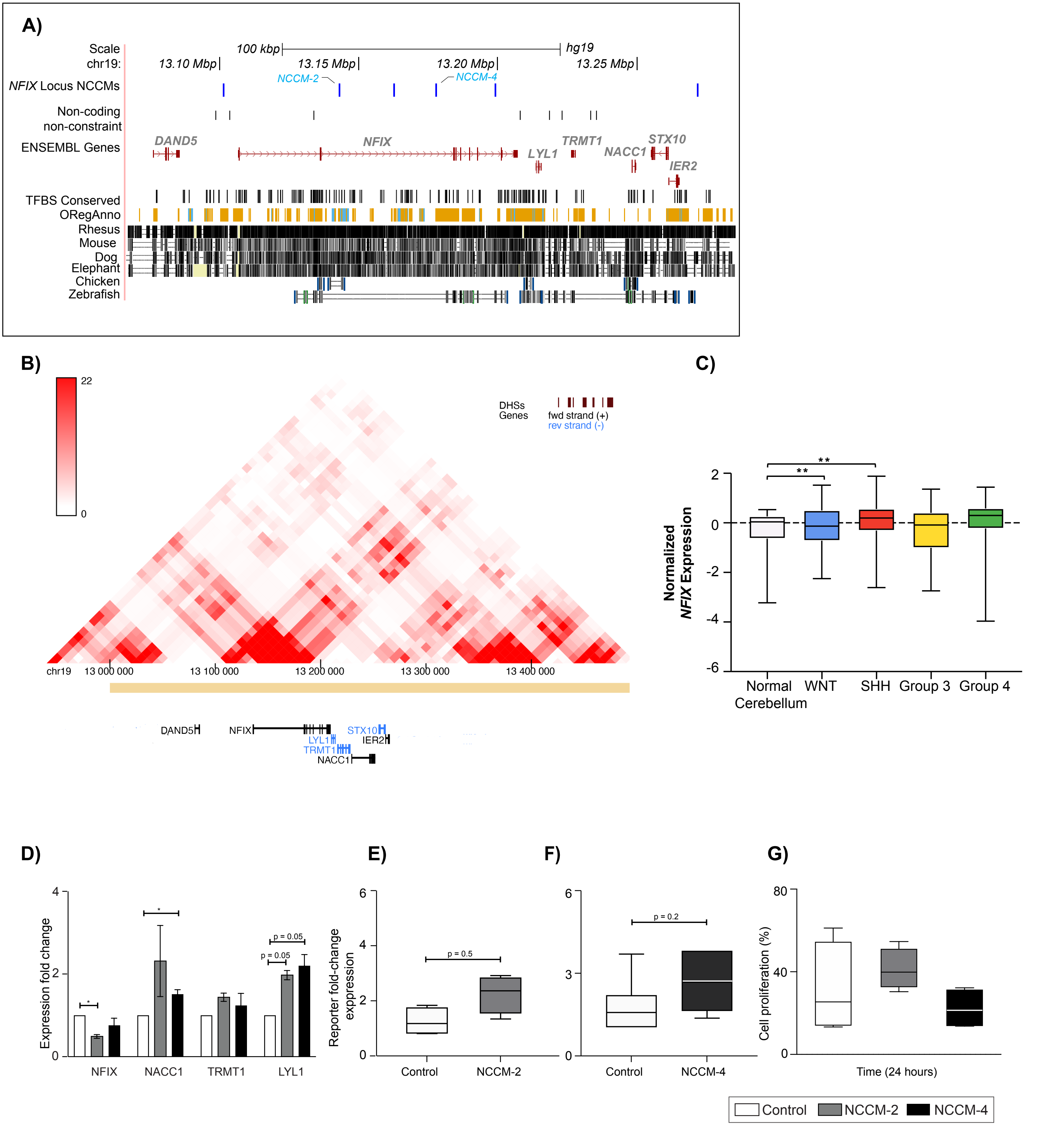

### Supplementary Figure 11

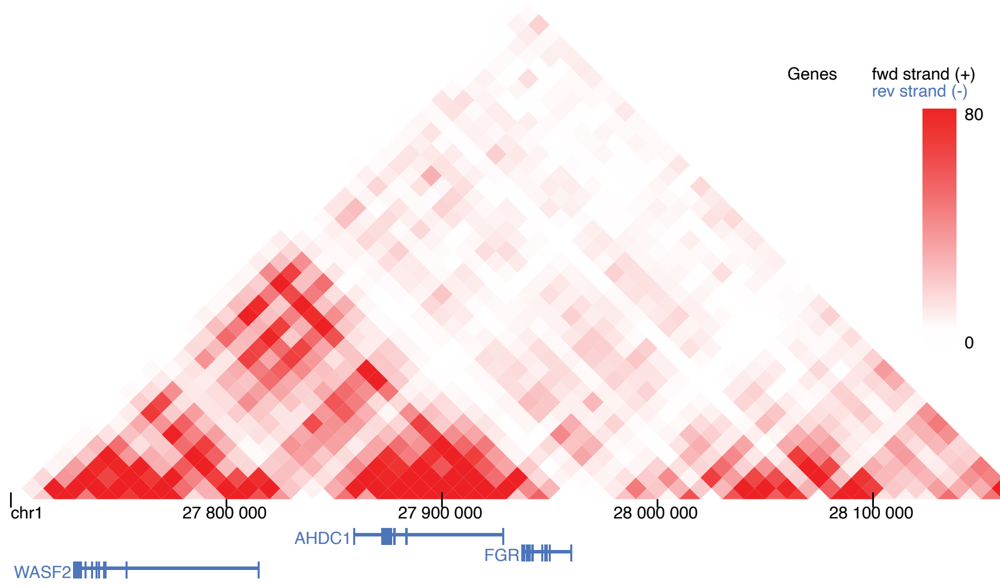

**Key for sheets in the current working document**

| Sheet Name | Table Name | Description |
|---|---|---|
| table 1 | table 1: Summary statistics for Pilocytic Astrocytoma (PA) cohort and their mutational burden | PA metadata + mutational burden summary |
| table 2 | table 2: Summary statistics for Medulloblastoma (MB) cohort and their mutational burden | MB metadata + mutational burden summary |
| table 3 | table 3: Genes with non-silent/protein-modifying variants for PA seen in ≥ 2 tumors + Significantly Mutated Genes (SMG) | PA genes with protein-modifiying changes + genes that are significantly mutated |
| table 4 | table 4: Candidate genes in PA with NCCMs ≥ 1 per 100 *kbp* | Genes in PA that have ≥ 1 NCCMs per 100 *kbp*. |
| table 5 | table 5: Functional enrichment/over- | Functional enrichment analysis was performed on gene set that is a union of protein modifying changes (table 3) and candidate genes that have ≥1 NCCMs per 100 *kbp* (table 4) |
| table 6 | table 6: Candidate gene in MB with ≥ 2 NCCMs per 100 *kbp* | Genes that have ≥ 2 NCCMs/100 kbp are shown with their rank order, and breakdown per subgroup per gene |
| table 7 | table 7: Annotation of candidate genes' associated NCCMs with data from the ENCODE portal and UCSC browser databases | NCCMs of candidate genes were annotated with regulatory data from ENCODE and UCSC Genombe browser. |
| table 8 | table 8: Genes in MB with ≥ 5 NCCMs in the target area (±100 *kbp*) | Genes that have ≥ 5 NCCMs in the ± 100 *kbp* flanking regions. |
| table 9 | table 9: Functional enrichment/over-representation analysis of MB NCCM genes (≥ 5, and ≥ 2 NCCMs per 100 *kbp*) | Functional enrichment analysis was performed on gene set that is a union of NCCM genes that have ≥ 5 NCCMs in the ± 100 kbp flanking regions (table 6) and those that have ≥ 2 NCCMs per 100 *kbp* (table 8). |
| table 10 | table 10: Coding mutations in the top 114 NCCM genes | NCCM counts in the ± 100 *kbp* flanking regions of genes known to be mutated in MB. |
| table 11 | table 11: NCCM rates in the significantly mutated genes of the MB cohort | Analysis with the tool sTRAP was implemented to to compute the binding affinity scores of related transcription factors in the context of reference and mutant sequences of transcription factor binding sites. |
| table 12 | table 12: NCCM counts for genes in previously implicated MB pathways, and those well-known to be mutated in MB | Oligos that were ordered for the EMSA analysis. |
| table 13 | table 13: sTRAP analysis for NCCMs associated with *BMP4* and *HOXB*-locus genes | |
| table 14 | table 14: oligos used for the EMSA experiments. | |

table 1: Summary statistics for Pilocytic Astrocytoma (PA) cohort and their muta

| ICGC Donor ID | Diagnosis | Gender | Age at diagnosis | SPM | SIM | Total |
|---|---|---|---|---|---|---|
| DO35869 | Pilocytic Astrocytoma | male | 1 | 120 | 12 | 132 |
| DO35871 | Pilocytic Astrocytoma | female | 4 | 145 | 4 | 149 |
| DO35874 | Pilocytic Astrocytoma | female | 7 | 99 | 7 | 106 |
| DO35877 | Pilocytic Astrocytoma | female | 15 | 121 | 5 | 126 |
| DO35880 | Pilocytic Astrocytoma | female | 3 | 101 | 12 | 113 |
| DO35883 | Pilocytic Astrocytoma | female | 11 | 453 | 25 | 478 |
| DO35886 | Pilocytic Astrocytoma | female | 14 | 364 | 15 | 379 |
| DO35889 | Pilocytic Astrocytoma | female | 5 | 90 | 3 | 93 |
| DO35892 | Pilocytic Astrocytoma | male | 4 | 131 | 7 | 138 |
| DO35895 | Pilocytic Astrocytoma | female | 23 | 464 | 25 | 489 |
| DO35898 | Pilocytic Astrocytoma | female | 15 | 283 | 22 | 305 |
| DO35901 | Pilocytic Astrocytoma | male | 4 | 514 | 16 | 530 |
| DO35904 | Pilocytic Astrocytoma | male | 2 | 137 | 8 | 145 |
| DO35907 | Pilocytic Astrocytoma | male | 3 | 90 | 4 | 94 |
| DO35916 | Pilocytic Astrocytoma | male | 12 | 199 | 9 | 208 |
| DO35919 | Pilocytic Astrocytoma | female | 14 | 709 | 33 | 742 |
| DO35922 | Pilocytic Astrocytoma | male | 4 | 101 | 3 | 104 |
| DO35925 | Pilocytic Astrocytoma | male | 2 | 95 | 7 | 102 |
| DO35928 | Pilocytic Astrocytoma | female | 11 | 402 | 17 | 419 |
| DO35931 | Pilocytic Astrocytoma | male | 9 | 240 | 30 | 270 |
| DO35934 | Pilocytic Astrocytoma | female | 9 | 219 | 11 | 230 |
| DO35937 | Pilocytic Astrocytoma | female | 15 | 225 | 5 | 230 |
| DO35940 | Pilocytic Astrocytoma | male | 3 | 67 | 2 | 69 |
| DO35943 | Pilocytic Astrocytoma | female | 6 | 288 | 18 | 306 |
| DO35946 | Pilocytic Astrocytoma | male | 9 | 266 | 11 | 277 |
| DO35949 | Pilocytic Astrocytoma | female | 3 | 161 | 9 | 170 |
| DO35952 | Pilocytic Astrocytoma | male | 6 | 136 | 7 | 143 |
| DO35955 | Pilocytic Astrocytoma | male | 10 | 231 | 7 | 238 |
| DO35959 | Pilocytic Astrocytoma | female | 5 | 166 | 2 | 168 |
| DO35961 | Pilocytic Astrocytoma | male | 9 | 141 | 11 | 152 |
| DO35964 | Pilocytic Astrocytoma | male | 13 | 358 | 12 | 370 |
| DO35967 | Pilocytic Astrocytoma | female | 5 | 126 | 12 | 138 |
| DO35976 | Pilocytic Astrocytoma | male | 4 | 129 | 14 | 143 |
| DO35979 | Pilocytic Astrocytoma | male | 16 | 1 284 | 18 | 1 302 |
| DO35982 | Pilocytic Astrocytoma | female | 6 | 362 | 32 | 394 |
| DO35985 | Pilocytic Astrocytoma | male | 6 | 122 | 10 | 132 |
| DO35988 | Pilocytic Astrocytoma | female | 2 | 172 | 4 | 176 |
| DO35991 | Pilocytic Astrocytoma | male | 9 | 478 | 18 | 496 |
| DO35994 | Pilocytic Astrocytoma | male | 14 | 702 | 19 | 721 |
| DO35997 | Pilocytic Astrocytoma | male | 6 | 118 | 4 | 122 |
| DO36000 | Pilocytic Astrocytoma | male | 14 | 206 | 7 | 213 |
| DO36003 | Pilocytic Astrocytoma | male | 12 | 264 | 18 | 282 |
| DO36006 | Pilocytic Astrocytoma | female | 8 | 322 | 6 | 328 |
| DO36009 | Pilocytic Astrocytoma | male | 9 | 286 | 17 | 303 |
| DO36012 | Pilocytic Astrocytoma | female | 4 | 107 | 4 | 111 |
| DO36015 | Pilocytic Astrocytoma | male | 11 | 190 | 7 | 197 |
| DO36018 | Pilocytic Astrocytoma | female | 4 | 176 | 6 | 182 |
| DO36021 | Pilocytic Astrocytoma | female | 6 | 109 | 4 | 113 |
| DO36024 | Pilocytic Astrocytoma | male | 8 | 57 | 1 | 58 |
| DO36027 | Pilocytic Astrocytoma | female | 3 | 180 | 3 | 183 |
| DO36030 | Pilocytic Astrocytoma | male | 12 | 1 022 | 38 | 1 060 |
| DO36033 | Pilocytic Astrocytoma | female | 21 | 1 117 | 37 | 1 154 |
| DO36036 | Pilocytic Astrocytoma | female | 4 | 90 | 7 | 97 |
| DO36039 | Pilocytic Astrocytoma | female | 7 | 182 | 10 | 192 |
| DO36042 | Pilocytic Astrocytoma | female | 15 | 120 | 6 | 126 |
| DO36048 | Pilocytic Astrocytoma | male | 9 | 107 | 11 | 118 |
| DO36051 | Pilocytic Astrocytoma | female | 1 | 128 | 6 | 134 |
| DO36054 | Pilocytic Astrocytoma | male | 15 | 146 | 1 | 147 |
| DO36057 | Pilocytic Astrocytoma | male | 7 | 58 | 5 | 63 |
| DO36062 | Pilocytic Astrocytoma | female | 4 | 104 | 11 | 115 |
| DO36065 | Pilocytic Astrocytoma | female | 5 | 104 | 7 | 111 |
| DO36068 | Pilocytic Astrocytoma | female | 5 | 198 | 9 | 207 |
| DO36071 | Pilocytic Astrocytoma | male | 12 | 195 | 3 | 198 |
| DO36074 | Pilocytic Astrocytoma | male | 14 | 224 | 11 | 235 |
| DO36077 | Pilocytic Astrocytoma | female | 50 | 513 | 11 | 524 |
| DO36080 | Pilocytic Astrocytoma | female | 40 | 516 | 16 | 532 |
| DO36083 | Pilocytic Astrocytoma | female | 9 | 129 | 4 | 133 |
| DO36087 | Pilocytic Astrocytoma | male | 25 | 430 | 29 | 459 |
| DO36091 | Pilocytic Astrocytoma | female | 20 | 489 | 33 | 522 |
| DO36095 | Pilocytic Astrocytoma | male | 7 | 84 | 4 | 88 |
| DO36099 | Pilocytic Astrocytoma | male | 3 | 77 | 3 | 80 |
| DO36103 | Pilocytic Astrocytoma | female | 14 | 348 | 18 | 366 |
| DO36107 | Pilocytic Astrocytoma | female | 11 | 126 | 6 | 132 |
| DO36111 | Pilocytic Astrocytoma | female | 10 | 364 | 21 | 385 |
| DO36115 | Pilocytic Astrocytoma | female | 22 | 323 | 12 | 335 |
| DO36119 | Pilocytic Astrocytoma | female | 17 | 374 | 14 | 388 |
| DO36123 | Pilocytic Astrocytoma | female | 2 | 159 | 4 | 163 |
| DO36127 | Pilocytic Astrocytoma | female | 7 | 141 | 11 | 152 |
| DO36131 | Pilocytic Astrocytoma | male | 12 | 268 | 10 | 278 |
| DO36135 | Pilocytic Astrocytoma | male | 1 | 94 | 5 | 99 |
| DO36139 | Pilocytic Astrocytoma | male | 2 | 90 | 8 | 98 |
| DO36143 | Pilocytic Astrocytoma | female | 11 | 95 | 8 | 103 |
| DO36147 | Pilocytic Astrocytoma | female | 5 | 100 | 2 | 102 |
| DO36151 | Pilocytic Astrocytoma | male | 5 | 199 | 25 | 224 |
| DO36155 | Pilocytic Astrocytoma | male | 1 | 163 | 16 | 179 |
| DO36159 | Pilocytic Astrocytoma | male | 7 | 86 | 13 | 99 |
| DO36163 | Pilocytic Astrocytoma | female | 29 | 40 | 1 | 41 |
| DO36167 | Pilocytic Astrocytoma | male | 5 | 237 | 18 | 255 |
| DO36171 | Pilocytic Astrocytoma | female | 8 | 391 | 11 | 402 |

**table 2: Summary statistics for Medulloblastoma (MB) cohort and their mutational burden**

| ICGC Donor ID | Diagnosis | Subgroup | Gender | Age at diagnosis | SPM | SIM | Total |
|---|---|---|---|---|---|---|---|
| DO35538 | Medulloblastoma | SHH | female | 6 | 741 | 54 | 795 |
| DO35539 | Medulloblastoma | Group 4 | male | 7 | 598 | 58 | 656 |
| DO35540 | Medulloblastoma | SHH | female | 16 | 1 170 | 180 | 1 350 |
| DO35541 | Medulloblastoma | Group 3 | female | 6 | 1 515 | 35 | 1 550 |
| DO35542 | Medulloblastoma | Group 4 | female | 14 | 1 729 | 214 | 1 943 |
| DO35543 | Medulloblastoma | Group 4 | male | 9 | 1 128 | 115 | 1 243 |
| DO35544 | Medulloblastoma | Group 4 | female | 11 | 976 | 185 | 1 161 |
| DO35545 | Medulloblastoma | Group 3 | male | 6 | 954 | 83 | 1 037 |
| DO35547 | Medulloblastoma | SHH | male | 3 | 347 | 25 | 372 |
| DO35550 | Medulloblastoma | Group 4 | male | 17 | 1 393 | 98 | 1 491 |
| DO35551 | Medulloblastoma | Group 3 | male | 3 | 606 | 53 | 659 |
| DO35552 | Medulloblastoma | Group 3 | female | 15 | 1 155 | 47 | 1 202 |
| DO35553 | Medulloblastoma | Group 3 | female | 13 | 1 729 | 275 | 2 004 |
| DO35554 | Medulloblastoma | Group 4 | female | 13 | 1 034 | 131 | 1 165 |
| DO35555 | Medulloblastoma | WNT | female | 9 | 1 279 | 195 | 1 474 |
| DO35556 | Medulloblastoma | SHH | female | 4 | 429 | 54 | 483 |
| DO35558 | Medulloblastoma | SHH | female | 10 | 2 503 | 226 | 2 729 |
| DO35559 | Medulloblastoma | Group 4 | male | 7 | 541 | 44 | 585 |
| DO35564 | Medulloblastoma | Group 4 | male | 8 | 1 349 | 133 | 1 482 |
| DO35566 | Medulloblastoma | SHH | male | 1 | 84 | 3 | 87 |
| DO35568 | Medulloblastoma | WNT | female | 14 | 2 101 | 383 | 2 484 |
| DO35569 | Medulloblastoma | Group 3 | female | 8 | 772 | 84 | 856 |
| DO35571 | Medulloblastoma | SHH | female | 13 | 4 437 | 328 | 4 765 |
| DO35572 | Medulloblastoma | SHH | female | 4 | 385 | 28 | 413 |
| DO35573 | Medulloblastoma | Group 3 | male | 4 | 1 615 | 283 | 1 898 |
| DO35574 | Medulloblastoma | SHH | female | 1 | 118 | 13 | 131 |
| DO35575 | Medulloblastoma | Group 4 | male | 6 | 1 107 | 94 | 1 201 |
| DO35576 | Medulloblastoma | Group 3 | male | 4 | 348 | 43 | 391 |
| DO35577 | Medulloblastoma | Group 4 | male | 3 | 216 | 25 | 241 |
| DO35581 | Medulloblastoma | Group 3 | male | 3 | 2 242 | 107 | 2 349 |
| DO35582 | Medulloblastoma | WNT | female | 6 | 1 244 | 115 | 1 359 |
| DO35584 | Medulloblastoma | Group 4 | male | 10 | 847 | 160 | 1 007 |
| DO35585 | Medulloblastoma | Group 3 | female | 10 | 1 556 | 210 | 1 766 |
| DO35586 | Medulloblastoma | Group 4 | male | 7 | 775 | 64 | 839 |
| DO35611 | Medulloblastoma | Group 4 | male | 19 | 2 335 | 132 | 2 467 |
| DO35612 | Medulloblastoma | Group 4 | male | 18 | 1 172 | 143 | 1 315 |
| DO35613 | Medulloblastoma | Group 4 | male | 18 | 1 115 | 85 | 1 200 |
| DO35616 | Medulloblastoma | Group 4 | female | 14 | 1 557 | 82 | 1 639 |
| DO35617 | Medulloblastoma | Group 4 | male | 12 | 1 773 | 112 | 1 885 |
| DO35618 | Medulloblastoma | Group 4 | female | 5 | 339 | 48 | 387 |
| DO35619 | Medulloblastoma | Group 4 | male | 4 | 657 | 43 | 700 |
| DO35620 | Medulloblastoma | SHH | male | 2 | 376 | 50 | 426 |
| DO35621 | Medulloblastoma | Group 3 | female | 4 | 776 | 101 | 877 |
| DO35622 | Medulloblastoma | Group 3 | male | 2 | 442 | 35 | 477 |
| DO35623 | Medulloblastoma | Group 4 | male | 15 | 1 186 | 128 | 1 314 |
| DO35624 | Medulloblastoma | Group 4 | male | 10 | 2 266 | 140 | 2 406 |
| DO35625 | Medulloblastoma | Group 4 | female | 15 | 2 734 | 287 | 3 021 |
| DO35626 | Medulloblastoma | Group 3 | male | 3 | 419 | 21 | 440 |
| DO35627 | Medulloblastoma | Group 3 | male | 3 | 507 | 51 | 558 |
| DO35628 | Medulloblastoma | Group 3 | male | 8 | 1 157 | 88 | 1 245 |
| DO35654 | Medulloblastoma | Group 3 | male | 16 | 4 193 | 562 | 4 755 |
| DO35668 | Medulloblastoma | Group 4 | male | 8 | 710 | 65 | 775 |
| DO35676 | Medulloblastoma | Group 3 | female | 9 | 458 | 99 | 557 |
| DO35678 | Medulloblastoma | Group 4 | female | 9 | 584 | 49 | 633 |
| DO35704 | Medulloblastoma | Group 3 | male | 3 | 934 | 233 | 1 167 |
| DO35715 | Medulloblastoma | Group 3 | male | 4 | 1 515 | 52 | 1 567 |
| DO35721 | Medulloblastoma | Group 4 | male | 9 | 838 | 112 | 950 |
| DO35723 | Medulloblastoma | Group 4 | female | 9 | 1 630 | 182 | 1 812 |
| DO35725 | Medulloblastoma | Group 4 | male | 11 | 658 | 38 | 696 |
| DO35729 | Medulloblastoma | Group 3 | male | 1 | 308 | 63 | 371 |
| DO35733 | Medulloblastoma | Group 3 | male | 3 | 433 | 107 | 540 |
| DO35737 | Medulloblastoma | Group 4 | male | 10 | 792 | 127 | 919 |
| DO35739 | Medulloblastoma | Group 4 | female | 10 | 4 196 | 302 | 4 498 |
| DO35741 | Medulloblastoma | Group 3 | female | 4 | 500 | 68 | 568 |
| DO35747 | Medulloblastoma | Group 4 | male | 8 | 788 | 90 | 878 |
| DO35749 | Medulloblastoma | Group 4 | male | 16 | 870 | 82 | 952 |
| DO35753 | Medulloblastoma | Group 4 | male | 11 | 1 125 | 115 | 1 240 |
| DO35755 | Medulloblastoma | Group 4 | female | 5 | 292 | 33 | 325 |
| DO35757 | Medulloblastoma | Group 4 | male | 8 | 897 | 66 | 963 |
| DO35763 | Medulloblastoma | Group 3 | female | 3 | 927 | 62 | 989 |
| DO35841 | Medulloblastoma | Group 4 | male | 16 | 1 631 | 196 | 1 827 |
| DO35845 | Medulloblastoma | Group 3 | male | 5 | 1 405 | 345 | 1 750 |
| DO35847 | Medulloblastoma | SHH | female | 5 | 402 | 19 | 421 |
| DO35857 | Medulloblastoma | Group 3 | male | 3 | 638 | 102 | 740 |
| DO48888 | Medulloblastoma | SHH | male | 20 | 2 196 | 196 | 2 392 |
| DO48889 | Medulloblastoma | SHH | female | 42 | 3 085 | 511 | 3 596 |
| DO48890 | Medulloblastoma | Group 4 | female | 23 | 1 315 | 89 | 1 404 |
| DO48891 | Medulloblastoma | SHH | female | 32 | 2 669 | 370 | 3 039 |
| DO48892 | Medulloblastoma | SHH | male | 23 | 1 498 | 310 | 1 808 |
| DO48893 | Medulloblastoma | SHH | male | 28 | 3 542 | 442 | 3 984 |
| DO48894 | Medulloblastoma | SHH | male | 27 | 2 751 | 243 | 2 994 |
| DO48895 | Medulloblastoma | SHH | male | 25 | 2 170 | 405 | 2 575 |
| DO48896 | Medulloblastoma | SHH | male | 22 | 1 145 | 149 | 1 294 |
| DO48898 | Medulloblastoma | Unclassified | female | 29 | 2 442 | 403 | 2 845 |
| DO48899 | Medulloblastoma | SHH | male | 23 | 2 717 | 555 | 3 272 |
| DO48900 | Medulloblastoma | SHH | female | 31 | 2 525 | 294 | 2 819 |
| DO48901 | Medulloblastoma | Group 4 | female | 29 | 4 081 | 198 | 4 279 |
| DO48902 | Medulloblastoma | SHH | female | 29 | 3 206 | 260 | 3 466 |
| DO48904 | Medulloblastoma | SHH | male | 38 | 3 734 | 562 | 4 296 |
| DO48905 | Medulloblastoma | SHH | male | 32 | 1 987 | 284 | 2 271 |
| DO48906 | Medulloblastoma | SHH | female | 49 | 5 223 | 751 | 5 974 |
| DO48907 | Medulloblastoma | SHH | male | 31 | 1 941 | 412 | 2 353 |
| DO48909 | Medulloblastoma | SHH | female | 39 | 5 416 | 547 | 5 963 |
| DO48911 | Medulloblastoma | SHH | female | 23 | 1 945 | 284 | 2 229 |
| DO48912 | Medulloblastoma | SHH | male | 35 | 3 327 | 363 | 3 690 |
| DO48914 | Medulloblastoma | SHH | female | 19 | 3 645 | 880 | 4 525 |
| DO48915 | Medulloblastoma | SHH | female | 31 | 4 319 | 499 | 4 818 |
| DO48916 | Medulloblastoma | Unclassified | female | 25 | 2 528 | 328 | 2 856 |
| DO48917 | Medulloblastoma | SHH | female | 22 | 2 864 | 201 | 3 065 |
| DO48918 | Medulloblastoma | Group 3 | female | 10 | 1 125 | 138 | 1 263 |
| DO48919 | Medulloblastoma | SHH | male | 13 | 2 631 | 260 | 2 891 |
| DO48920 | Medulloblastoma | Group 3 | female | 7 | 1 758 | 88 | 1 846 |
| DO48921 | Medulloblastoma | Group 4 | male | 4 | 1 960 | 287 | 2 247 |
| DO48922 | Medulloblastoma | Group 3 | male | 10 | 946 | 96 | 1 042 |
| DO48923 | Medulloblastoma | Group 3 | female | 3 | 1 740 | 168 | 1 908 |
| DO48924 | Medulloblastoma | Group 3 | male | 4 | 237 | 15 | 252 |
| DO48925 | Medulloblastoma | Group 4 | male | 6 | 707 | 83 | 790 |
| DO48927 | Medulloblastoma | Group 4 | female | 9 | 924 | 70 | 994 |
| DO48929 | Medulloblastoma | Group 4 | female | 6 | 993 | 86 | 1 079 |
| DO48930 | Medulloblastoma | Group 3 | female | 8 | 1 589 | 98 | 1 687 |
| DO48932 | Medulloblastoma | Group 4 | female | 4 | 1 119 | 89 | 1 208 |
| DO48934 | Medulloblastoma | Group 3 | male | 5 | 721 | 103 | 824 |
| DO48935 | Medulloblastoma | Group 3 | male | 7 | 970 | 69 | 1 039 |
| DO48936 | Medulloblastoma | Group 3 | male | 9 | 1 434 | 59 | 1 493 |
| DO48937 | Medulloblastoma | Group 3 | male | 21 | 1 058 | 187 | 1 245 |
| DO48938 | Medulloblastoma | Group 3 | male | 11 | 2 285 | 339 | 2 624 |
| DO48939 | Medulloblastoma | SHH | female | 16 | 733 | 112 | 845 |
| DO48940 | Medulloblastoma | Group 4 | male | 11 | 1 688 | 106 | 1 794 |
| DO48941 | Medulloblastoma | Group 4 | male | 6 | 590 | 53 | 643 |
| DO48943 | Medulloblastoma | Group 4 | female | 27 | 1 438 | 228 | 1 666 |
| DO48944 | Medulloblastoma | SHH | male | 26 | 1 976 | 312 | 2 288 |
| DO48945 | Medulloblastoma | SHH | male | 22 | 1 878 | 303 | 2 181 |
| DO48946 | Medulloblastoma | SHH | female | 3 | 1 276 | 208 | 1 484 |
| DO48948 | Medulloblastoma | Group 4 | male | 12 | 1 293 | 50 | 1 343 |
| DO48949 | Medulloblastoma | Group 4 | male | 3 | 891 | 95 | 986 |
| DO48950 | Medulloblastoma | Group 4 | male | 3 | 325 | 29 | 354 |
| DO48951 | Medulloblastoma | Unclassified | female | 16 | 948 | 148 | 1 096 |
| DO48952 | Medulloblastoma | Group 3 | female | 6 | 1 005 | 38 | 1 043 |
| DO48953 | Medulloblastoma | Group 4 | female | 8 | 1 454 | 105 | 1 559 |
| DO48955 | Medulloblastoma | Group 3 | female | 9 | 2 002 | 117 | 2 119 |
| DO48957 | Medulloblastoma | Group 3 | male | 5 | 1 258 | 145 | 1 403 |
| DO48958 | Medulloblastoma | Group 4 | male | 5 | 1 172 | 137 | 1 309 |
| DO48959 | Medulloblastoma | Group 3 | female | 9 | 774 | 141 | 915 |
| DO48961 | Medulloblastoma | Group 4 | female | 5 | 623 | 72 | 695 |
| DO48963 | Medulloblastoma | Group 4 | male | 4 | 676 | 104 | 780 |
| DO48964 | Medulloblastoma | Group 3 | male | 8 | 1 082 | 277 | 1 359 |
| DO48965 | Medulloblastoma | Group 4 | male | 4 | 423 | 58 | 481 |
| DO48966 | Medulloblastoma | Group 4 | female | 3 | 1 142 | 133 | 1 275 |
| DO48967 | Medulloblastoma | Group 4 | female | 9 | 609 | 57 | 666 |
| DO48968 | Medulloblastoma | Group 4 | male | 2 | 445 | 63 | 508 |
| DO48969 | Medulloblastoma | Group 4 | female | 18 | 572 | 72 | 644 |
| DO48970 | Medulloblastoma | Group 3 | female | 11 | 217 | 22 | 239 |
| DO48971 | Medulloblastoma | Group 4 | male | 10 | 2 133 | 276 | 2 409 |
| DO48972 | Medulloblastoma | Group 4 | female | 8 | 1 181 | 153 | 1 334 |
| DO48973 | Medulloblastoma | Group 3 | female | 10 | 1 754 | 202 | 1 956 |
| DO48974 | Medulloblastoma | Group 4 | female | 8 | 535 | 80 | 615 |

table 3: Genes with non-silent/protein-modifying variants for PA seen

| Gene | No. of non-silent changes | Observed in how many tumors | SMG? |
|---|---|---|---|
| ***BRAF*** | 6 | 6 | **Yes** |
| *FGFR1* | 6 | 5 | No |
| *NF1* | 5 | 4 | No |
| *KRAS* | 4 | 2 | No |
| *AHNAK2* | 2 | 2 | No |
| *AMBP* | 2 | 1 | No |
| *FLG* | 2 | 2 | No |
| *IL4R* | 2 | 2 | No |
| *KIAA1549* | 2 | 2 | No |
| *PTPN11* | 2 | 2 | No |
| *SETD2* | 2 | 2 | No |

table 4: Candidate genes in PA with NCCMs ≥ 1 per 100 *kbp*

| Index | Mutation Name | Mutation Name *(Shared Loci separated by ',' and colored)* | Top NCCM Gene | ICGC Donor ID | Gender | Age at diagnosis | Chrom# | Posi. | Ref. | Alt. | Mutation Classification | phyloP |
|---|---|---|---|---|---|---|---|---|---|---|---|---|
| 1 | NDUFB2.NCCM-1 | ADCK2.NCCM-1, BRAF.NCCM-1, NDUFB2.NCCM-1 | *NDUFB2* | DO36071 | male | 12 | 7 | 140347391 | C | A | RNA | 2,8 |
| 2 | NDUFB2.NCCM-2 | ADCK2.NCCM-2, BRAF.NCCM-2, NDUFB2.NCCM-2 | *NDUFB2* | DO36065 | female | 5 | 7 | 140474641 | A | C | Intron | 3,6 |
| 3 | NDUFB2.NCCM-3 | ADCK2.NCCM-3, BRAF.NCCM-3, NDUFB2.NCCM-3 | *NDUFB2* | DO35901 | male | 4 | 7 | 140491337 | T | A | Intron | 1,3 |
| 4 | NDUFB2.NCCM-4 | BRAF.NCCM-4, NDUFB2.NCCM-4 | *NDUFB2* | DO35961 | male | 9 | 7 | 140500287 | A | T | Intron | 4,7 |
| 5 | NDUFB2.NCCM-5 | BRAF.NCCM-5, NDUFB2.NCCM-5 | *NDUFB2* | DO35961 | male | 9 | 7 | 140500288 | G | T | Intron | 8,8 |
| 6 | ADCK2.NCCM-1 | ADCK2.NCCM-1, BRAF.NCCM-1, NDUFB2.NCCM-1 | *ADCK2* | DO36071 | male | 12 | 7 | 140347391 | C | A | RNA | 2,8 |
| 7 | ADCK2.NCCM-2 | ADCK2.NCCM-2, BRAF.NCCM-2, NDUFB2.NCCM-2 | *ADCK2* | DO36065 | female | 5 | 7 | 140474641 | A | C | Intron | 3,6 |
| 8 | ADCK2.NCCM-3 | ADCK2.NCCM-3, BRAF.NCCM-3, NDUFB2.NCCM-3 | *ADCK2* | DO35901 | male | 4 | 7 | 140491337 | T | A | Intron | 1,3 |
| 9 | PRUNE.NCCM-1 | PRUNE.NCCM-1 | *PRUNE* | DO36167 | male | 5 | 1 | 150888475 | T | A | IGR | 1,7 |
| 10 | PRUNE.NCCM-2 | PRUNE.NCCM-2 | *PRUNE* | DO35964 | male | 13 | 1 | 150979302 | A | C | Intron | 3,8 |
| 11 | PRUNE.NCCM-3 | PRUNE.NCCM-3 | *PRUNE* | DO36003 | male | 12 | 1 | 151094473 | A | G | 3'UTR | 3,0 |
| 12 | RBM18.NCCM-1 | MRRF.NCCM-1,RBM18.NCCM-1 | *RBM18* | DO36087 | male | 25 | 9 | 124993145 | C | T | 5'Flank | 1,5 |
| 13 | RBM18.NCCM-2 | MRRF.NCCM-2,RBM18.NCCM-2 | *RBM18* | DO35991 | male | 9 | 9 | 125070511 | G | T | Intron | 2,2 |
| 14 | RBM18.NCCM-3 | MRRF.NCCM-3,RBM18.NCCM-3 | *RBM18* | DO35892 | male | 4 | 9 | 125117238 | G | T | IGR | 1,7 |
| 15 | ACRC.NCCM-1 | ACRC.NCCM-1,OGT.NCCM-1 | *ACRC* | DO36091 | female | 20 | X | 70711926 | C | T | Intron | 4,8 |
| 16 | ACRC.NCCM-2 | ACRC.NCCM-2,OGT.NCCM-2 | *ACRC* | DO35886 | female | 14 | X | 70795136 | G | T | 3'UTR | 4,8 |
| 17 | ACRC.NCCM-3 | ACRC.NCCM-3,OGT.NCCM-3 | *ACRC* | DO36171 | female | 8 | X | 70888031 | C | T | RNA | 2,2 |
| 18 | OGT.NCCM-1 | ACRC.NCCM-1,OGT.NCCM-1 | *OGT* | DO36091 | female | 20 | X | 70711926 | C | T | Intron | 4,8 |
| 19 | OGT.NCCM-2 | ACRC.NCCM-2,OGT.NCCM-2 | *OGT* | DO35886 | female | 14 | X | 70795136 | G | T | 3'UTR | 4,8 |
| 20 | OGT.NCCM-3 | ACRC.NCCM-3,OGT.NCCM-3 | *OGT* | DO36171 | female | 8 | X | 70888031 | C | T | RNA | 2,2 |
| 21 | BRAF.NCCM-1 | ADCK2.NCCM-1, BRAF.NCCM-1, NDUFB2.NCCM-1 | *BRAF* | DO36071 | male | 12 | 7 | 140347391 | C | A | RNA | 2,8 |
| 22 | BRAF.NCCM-2 | ADCK2.NCCM-2, BRAF.NCCM-2, NDUFB2.NCCM-2 | *BRAF* | DO36065 | female | 5 | 7 | 140474641 | A | C | Intron | 3,6 |
| 23 | BRAF.NCCM-3 | ADCK2.NCCM-3, BRAF.NCCM-3, NDUFB2.NCCM-3 | *BRAF* | DO35901 | male | 4 | 7 | 140491337 | T | A | Intron | 1,3 |
| 24 | BRAF.NCCM-4 | BRAF.NCCM-4, NDUFB2.NCCM-4 | *BRAF* | DO35961 | male | 9 | 7 | 140500287 | A | T | Intron | 4,7 |
| 25 | BRAF.NCCM-5 | BRAF.NCCM-5, NDUFB2.NCCM-5 | *BRAF* | DO35961 | male | 9 | 7 | 140500288 | G | T | Intron | 8,8 |
| 26 | MRRF.NCCM-1 | MRRF.NCCM-1,RBM18.NCCM-1 | *MRRF* | DO36087 | male | 25 | 9 | 124993145 | C | T | 5'Flank | 1,5 |
| 27 | MRRF.NCCM-2 | MRRF.NCCM-2,RBM18.NCCM-2 | *MRRF* | DO35991 | male | 9 | 9 | 125070511 | G | T | Intron | 2,2 |
| 28 | MRRF.NCCM-3 | MRRF.NCCM-3,RBM18.NCCM-3 | *MRRF* | DO35892 | male | 4 | 9 | 125117238 | G | T | IGR | 1,7 |
| 29 | KIT.NCCM-1 | KIT.NCCM-1 | *KIT* | DO36018 | female | 4 | 4 | 55474554 | G | A | lincRNA | 1,6 |
| 30 | KIT.NCCM-2 | KIT.NCCM-2 | *KIT* | DO36127 | female | 7 | 4 | 55592271 | C | T | Intron | 4,0 |
| 31 | KIT.NCCM-3 | KIT.NCCM-3 | *KIT* | DO36030 | male | 12 | 4 | 55615737 | T | G | IGR | 2,2 |
| 32 | OR4E2.NCCM-1 | OR4E2.NCCM-1 | *OR4E2* | DO35925 | male | 2 | 14 | 22033814 | G | T | IGR | 2,8 |
| 33 | OR4E2.NCCM-2 | OR4E2.NCCM-2 | *OR4E2* | DO36167 | male | 5 | 14 | 22180092 | G | T | RNA | 1,7 |
| 34 | OR1L6.NCCM-1 | OR1L6.NCCM-1 | *OR1L6* | DO36030 | male | 12 | 9 | 125507513 | C | A | IGR | 1,5 |
| 35 | OR1L6.NCCM-2 | OR1L6.NCCM-2 | *OR1L6* | DO36155 | male | 1 | 9 | 125610709 | C | A | 3'UTR | 2,4 |
| 36 | FKSG68.NCCM-1 | FKSG68.NCCM-1 | *FKSG68* | DO35940 | male | 3 | 21 | 37221612 | C | T | IGR | 2,3 |
| 37 | FKSG68.NCCM-2 | FKSG68.NCCM-2 | *FKSG68* | DO36000 | male | 14 | 21 | 37357519 | C | A | IGR | 1,2 |

Key for table 4.

| Colı | Name | Description |
|---|---|---|
| A | Index | Sequential indexing for the current dataset |
| B | Mutation Name | Name of the mutation, formed by prefixing gene + consecutive numbers |
| C | Mutation Name (Shared Loci separated by ',' and colored) | If present, other shared loci are shown as well. Otherwise name is the same as col. B |
| D | Top NCCM Gene | Genes, whose regulatory potential is examined |
| E | ICGC Donor ID | ICGC Sample ID |
| F | Gender | Gender/Sex for the patient |
| G | Age at diagnosis | Age at which the disease was diagnosed. Colored cells represent pediatric cohort, i.e. age <18 |
| H | Chrom# | Chromosome number |
| I | Posi. | Position(s) where SPM/SIM is observed |
| J | Ref. | Reference allele |
| K | Alt. | Alternate allele |
| L | Mutation Classification | Type of change for the SPM/SIM. |
| M | phyloP | phyloP scores. |

**table 5: Functional enrichment/over-**

| Source Database | ID | Pathway/Ontology | *p*-value (Adjusted) | Total no. of genes in Pathway/ Ontology | No. of query genes overlapping | Genes |
|---|---|---|---|---|---|---|
| REACTOME | REAC:R-HSA-5663202 | Diseases of signal transduction by growth factor receptors and second messengers | 1,86×10^-04 | 380 | 7 | *NF1,FGFR1,KIAA1549,PTPN11,KRAS,BRAF,KIT* |
| REACTOME | REAC:R-HSA-6802957 | Oncogenic MAPK signaling | 1,70×10^-03 | 82 | 4 | *NF1,KIAA1549,KRAS,BRAF* |
| KEGG | KEGG:04014 | Ras signaling pathway | 1,79×10^-03 | 231 | 5 | *NF1,FGFR1,PTPN11,KRAS,KIT* |
| KEGG | KEGG:05224 | Breast cancer | 5,46×10^-03 | 147 | 4 | *FGFR1,KRAS,BRAF,KIT* |
| KEGG | KEGG:04010 | MAPK signaling pathway | 5,68×10^-03 | 294 | 5 | *NF1,FGFR1,KRAS,BRAF,KIT* |
| REACTOME | REAC:R-HSA-5684996 | MAPK1/MAPK3 signaling | 1,06×10^-02 | 269 | 5 | *NF1,PTPN11,KRAS,BRAF,KIT* |
| KEGG | KEGG:05221 | Acute myeloid leukemia | 1,10×10^-02 | 67 | 3 | *KRAS,BRAF,KIT* |
| KEGG | KEGG:05211 | Renal cell carcinoma | 1,15×10^-02 | 68 | 3 | *PTPN11,KRAS,BRAF* |
| KEGG | KEGG:05230 | Central carbon metabolism in cancer | 1,25×10^-02 | 70 | 3 | *FGFR1,KRAS,KIT* |
| KEGG | KEGG:05218 | Melanoma | 1,36×10^-02 | 72 | 3 | *FGFR1,KRAS,BRAF* |
| KEGG | KEGG:05220 | Chronic myeloid leukemia | 1,60×10^-02 | 76 | 3 | *PTPN11,KRAS,BRAF* |
| KEGG | KEGG:01521 | EGFR tyrosine kinase inhibitor resistance | 1,79×10^-02 | 79 | 3 | *NF1,KRAS,BRAF* |
| KEGG | KEGG:05205 | Proteoglycans in cancer | 1,97×10^-02 | 205 | 4 | *FGFR1,PTPN11,KRAS,BRAF* |
| REACTOME | REAC:R-HSA-5683057 | MAPK family signaling cascades | 2,02×10^-02 | 308 | 5 | *NF1,PTPN11,KRAS,BRAF,KIT* |
| KEGG | KEGG:04015 | Rap1 signaling pathway | 2,16×10^-02 | 210 | 4 | *FGFR1,KRAS,BRAF,KIT* |
| KEGG | KEGG:05215 | Prostate cancer | 3,28×10^-02 | 97 | 3 | *FGFR1,KRAS,BRAF* |
| GO:Biological Proc | GO:0043408 | regulation of MAPK cascade | 6,23×10^-03 | 734 | 7 | *NF1,FGFR1,PTPN11,AMBP,KRAS,BRAF,KIT* |
| GO:Biological Proc | GO:0000165 | MAPK cascade | 3,75×10^-02 | 965 | 7 | *NF1,FGFR1,PTPN11,AMBP,KRAS,BRAF,KIT* |
| GO:Biological Proc | GO:0048871 | multicellular organismal homeostasis | 1,44×10^-02 | 533 | 6 | *NF1,FLG,PTPN11,IL4R,KRAS,OGT* |
| GO:Biological Proc | GO:0043405 | regulation of MAP kinase activity | 1,85×10^-02 | 316 | 5 | *NF1,FGFR1,PTPN11,KRAS,KIT* |
| GO:Biological Proc | GO:0008542 | visual learning | 1,65×10^-04 | 45 | 4 | *NF1,KRAS,BRAF,KIT* |
| GO:Biological Proc | GO:0007632 | visual behavior | 2,54×10^-04 | 50 | 4 | *NF1,KRAS,BRAF,KIT* |
| GO:Biological Proc | GO:0008306 | associative learning | 1,63×10^-03 | 79 | 4 | *NF1,KRAS,BRAF,KIT* |
| GO:Biological Proc | GO:0007612 | learning | 1,94×10^-02 | 147 | 4 | *NF1,KRAS,BRAF,KIT* |
| GO:Biological Proc | GO:0021897 | forebrain astrocyte development | 1,55×10^-03 | 2 | 2 | *NF1,KRAS* |
| GO:Biological Proc | GO:0021896 | forebrain astrocyte differentiation | 1,55×10^-03 | 2 | 2 | *NF1,KRAS* |

**table 6: Candidate gene in MB with ≥ 2 NCCMs per 100 *kbp***

| Ranl | Candidate Gene | Description | Tota | WNT | SHH | Group 3 | Group 4 | Unclassified |
|---|---|---|---|---|---|---|---|---|
| 1 | *NABP1* | SOSS complex subunit B2 | 8 | 0 | 2 | 3 | 3 | 0 |
| 2 | *IFIH1* | Interferon-induced helicase C domain-containing protein 1 | 8 | 0 | 1 | 0 | 7 | 0 |
| 3 | *COL1A2* | Collagen alpha-2(I) chain | 7 | 0 | 2 | 3 | 2 | 0 |
| 4 | *PAX6* | Paired box protein Pax-6 | 7 | 0 | 3 | 3 | 1 | 0 |
| 5 | *DLX6* | Homeobox protein DLX-6 | 6 | 0 | 2 | 1 | 3 | 0 |
| 6 | *HAND2* | Heart- and neural crest derivatives-expressed protein 2 | 6 | 0 | 2 | 1 | 3 | 0 |
| 7 | *CASD1* | N-acetylneuraminate 9-O-acetyltransferase | 7 | 0 | 2 | 3 | 2 | 0 |
| 8 | *NFE2L1* | Nuclear factor erythroid 2-related factor 1 | 6 | 0 | 3 | 0 | 3 | 0 |
| 9 | *SNX11* | Sorting nexin-11 | 6 | 0 | 2 | 0 | 4 | 0 |
| 10 | *CRB2* | Protein crumbs homolog 2 | 6 | 0 | 0 | 4 | 2 | 0 |
| 11 | *FGR* | Tyrosine-protein kinase Fgr | 6 | 0 | 4 | 1 | 0 | 1 |
| 12 | *NACC1* | Nucleus accumbens-associated protein 1 | 6 | 0 | 4 | 1 | 0 | 1 |
| 13 | *GSS* | Glutathione synthetase | 6 | 0 | 3 | 1 | 2 | 0 |
| 14 | *ZNF462* | Zinc finger protein 462 | 9 | 0 | 1 | 2 | 6 | 0 |
| 15 | *AXIN2* | Axin-2 | 6 | 0 | 0 | 2 | 4 | 0 |
| 16 | *CBX1* | Chromobox protein homolog 1 | 6 | 0 | 2 | 0 | 4 | 0 |
| 17 | *ABCC12* | Multidrug resistance-associated protein 9 | 7 | 0 | 2 | 1 | 3 | 1 |
| 18 | *MAP3K7* | Mitogen-activated protein kinase kinase kinase 7 | 7 | 0 | 2 | 1 | 3 | 1 |
| 19 | *UBXN11* | UBX domain-containing protein 11 | 6 | 0 | 3 | 2 | 0 | 1 |
| 20 | *NUAK1* | NUAK family SNF1-like kinase 1 | 7 | 0 | 6 | 0 | 0 | 1 |
| 21 | *GTF2E1* | General transcription factor IIE subunit 1 | 6 | 1 | 2 | 1 | 2 | 0 |
| 22 | *DKFZP547L112* | Putative uncharacterized protein | 5 | 0 | 2 | 1 | 2 | 0 |
| 23 | *OR5G3* | Olfactory Receptor Family 5 Subfamily G Member 3 | 5 | 0 | 1 | 3 | 1 | 0 |
| 24 | *ATOH1* | Protein atonal homolog 1 | 5 | 0 | 3 | 1 | 1 | 0 |
| 25 | *CTD-2144E22.5* | Putative uncharacterized protein | 5 | 1 | 3 | 1 | 0 | 0 |
| 26 | *WBP1* | WW domain binding protein 1 | 5 | 0 | 1 | 1 | 2 | 1 |
| 27 | *PSMB11* | Proteasome subunit beta type-11 | 5 | 0 | 1 | 2 | 2 | 0 |
| 28 | *MRPL53* | Mitochondrial ribosomal protein L53 | 5 | 0 | 1 | 1 | 2 | 1 |
| 29 | *PCSK1* | Neuroendocrine convertase 1 | 6 | 0 | 2 | 2 | 2 | 0 |
| 30 | *HOXB1* | Homeobox protein Hox-B1 | 5 | 0 | 1 | 3 | 1 | 0 |
| 31 | *KCNA3* | Potassium voltage-gated channel subfamily A member 3 | 5 | 0 | 3 | 0 | 2 | 0 |
| 32 | *PNMT* | Phenylethanolamine N-methyltransferase | 5 | 0 | 2 | 0 | 3 | 0 |
| 33 | *MOGS* | Mannosyl-oligosaccharide glucosidase | 5 | 0 | 1 | 1 | 2 | 1 |
| 34 | *TCAP* | Telethonin | 5 | 0 | 2 | 0 | 3 | 0 |
| 35 | *HMGB4* | Canonical high mobility Group | 5 | 0 | 2 | 1 | 2 | 0 |
| 36 | *SH3BGRL3* | SH3 domain-binding glutamic acid-rich-like protein 3 | 5 | 0 | 2 | 2 | 0 | 1 |
| 37 | *CD52* | CAMPATH-1 antigen | 5 | 0 | 3 | 2 | 0 | 0 |
| 38 | *IRX5* | Iroquois-class homeodomain protein IRX-5 | 5 | 0 | 2 | 2 | 1 | 0 |
| 39 | *NKX2-2* | Homeobox protein Nkx-2.2 | 5 | 0 | 4 | 1 | 0 | 0 |
| 40 | *LYL1* | Protein lyl-1 | 5 | 0 | 3 | 1 | 0 | 1 |
| 41 | *TLCD1* | Calfacilitin | 5 | 0 | 3 | 0 | 2 | 0 |
| 42 | *DLX5* | Homeobox protein DLX-5 | 5 | 0 | 2 | 1 | 2 | 0 |
| 43 | *IER2* | Immediate early response gene 2 protein | 5 | 0 | 4 | 1 | 0 | 0 |
| 44 | *INO80B* | INO80 complex subunit B | 5 | 0 | 1 | 1 | 2 | 1 |
| 45 | *HOXB2* | Homeobox protein Hox-B2 | 5 | 0 | 1 | 3 | 1 | 0 |
| 46 | *C14orf119* | Uncharacterized protein C14orf119 | 5 | 0 | 1 | 2 | 2 | 0 |
| 47 | *BCDIN3D* | Pre-miRNA 5'-monophosphate methyltransferase | 5 | 0 | 2 | 1 | 2 | 0 |
| 48 | *RPL23A* | 60S ribosomal protein L23a | 5 | 0 | 3 | 0 | 2 | 0 |
| 49 | *ZBTB18* | Zinc finger and BTB domain-containing protein 18 | 5 | 1 | 1 | 1 | 2 | 0 |
| 50 | *TRAF4* | TNF receptor-associated factor 4 | 5 | 0 | 3 | 0 | 2 | 0 |
| 51 | *STX10* | Syntaxin-10 | 5 | 0 | 4 | 1 | 0 | 0 |
| 52 | *CDH24* | Cadherin-24 | 5 | 0 | 1 | 2 | 2 | 0 |
| 53 | *BMP4* | Bone morphogenetic protein 4 | 5 | 0 | 0 | 3 | 2 | 0 |
| 54 | *TTC31* | Tetratricopeptide repeat domain containing | 5 | 0 | 1 | 1 | 2 | 1 |
| 55 | *CNGA4* | Cyclic nucleotide-gated cation channel alpha-4 | 5 | 0 | 3 | 1 | 1 | 0 |
| 56 | *ZC3H12A* | Endoribonuclease ZC3H12A | 5 | 0 | 4 | 0 | 1 | 0 |
| 57 | *ZFHX4* | Zinc finger homeobox protein 4 | 9 | 0 | 8 | 1 | 0 | 0 |
| 58 | *TRMT5* | tRNA (guanine(37)-N1)-methyltransferase | 5 | 0 | 3 | 1 | 1 | 0 |
| 59 | *DAND5* | DAN domain family member 5 | 5 | 0 | 4 | 0 | 0 | 1 |
| 60 | *CCDC142* | Coiled-coil domain containing 142 | 5 | 0 | 1 | 1 | 2 | 1 |
| 61 | *TAC4* | Tachykinin-4 | 5 | 0 | 0 | 0 | 5 | 0 |
| 62 | *GCA* | Grancalcin | 6 | 0 | 1 | 0 | 5 | 0 |
| 63 | *TRMT1* | tRNA (guanine(26)-N(2))-dimethyltransferase | 5 | 0 | 3 | 1 | 0 | 1 |
| 64 | *ZIC3* | Zinc finger protein ZIC 3 | 5 | 1 | 4 | 0 | 0 | 0 |
| 65 | *ISL1* | Insulin gene enhancer protein ISL-1 | 5 | 0 | 1 | 3 | 1 | 0 |
| 66 | *ZNF683* | Tissue-resident T-cell transcription regulator protein ZNF683 | 5 | 0 | 3 | 2 | 0 | 0 |
| 67 | *PDK4* | [Pyruvate dehydrogenase (acetyl-transferring)] kinase isozyme 4, mitochondrial | 5 | 0 | 2 | 0 | 3 | 0 |
| 68 | *PGLYRP3* | Peptidoglycan recognition protein 3 | 5 | 0 | 3 | 1 | 1 | 0 |
| 69 | *MEF2C* | Myocyte-specific enhancer factor 2C | 9 | 1 | 4 | 1 | 3 | 0 |
| 70 | *HDAC3* | Histone deacetylase 3 | 5 | 0 | 2 | 1 | 1 | 1 |
| 71 | *NEK8* | Serine/threonine-protein kinase Nek8 | 5 | 0 | 3 | 0 | 2 | 0 |
| 72 | *FSD1L* | FSD1-like protein | 7 | 0 | 1 | 3 | 3 | 0 |
| 73 | *C16orf47* | Zinc Finger Homeobox 3 | 5 | 1 | 4 | 0 | 0 | 0 |
| 74 | *PGLYRP4* | Peptidoglycan recognition protein 4 | 5 | 0 | 3 | 1 | 1 | 0 |
| 75 | *C2orf91* | Uncharacterized protein C2orf91 | 5 | 0 | 1 | 0 | 4 | 0 |
| 76 | *FAM180A* | Protein FAM180A | 5 | 1 | 3 | 1 | 0 | 0 |
| 77 | *AHDC1* | AT-hook DNA-binding motif-containing protein 1 | 6 | 0 | 4 | 1 | 0 | 1 |
| 78 | *UGT2A3* | UDP-glucuronosyltransferase 2A3 | 5 | 0 | 1 | 2 | 1 | 1 |
| 79 | *MYH7B* | Myosin-7B | 5 | 0 | 2 | 1 | 2 | 0 |
| 80 | *PGAP3* | Post-GPI attachment to proteins factor 3 | 5 | 0 | 2 | 0 | 3 | 0 |
| 81 | *STARD3* | StAR-related lipid transfer protein 3 | 5 | 0 | 2 | 0 | 3 | 0 |
| 82 | *SPA17* | Sperm surface protein Sp17 | 5 | 0 | 2 | 1 | 2 | 0 |
| 83 | *LSM6* | U6 snRNA-associated Sm-like protein LSm6 | 5 | 0 | 0 | 2 | 3 | 0 |
| 84 | *FOSL2* | Fos-related antigen 2 | 5 | 0 | 2 | 0 | 3 | 0 |
| 85 | *SUPV3L1* | ATP-dependent RNA helicase SUPV3L1, mitochondrial | 5 | 0 | 2 | 2 | 1 | 0 |
| 86 | *TCP11* | T-complex protein 11 homolog | 5 | 0 | 2 | 1 | 2 | 0 |
| 87 | *AIM1L* | Beta/gamma crystallin domain-containing protein 2 | 5 | 0 | 3 | 2 | 0 | 0 |
| 88 | *RP11-366L20.2* | HMGA2 Antisense RNA 1 gene. | 5 | 0 | 0 | 1 | 3 | 1 |
| 89 | *SLC7A11* | Cystine/glutamate transporter | 6 | 0 | 1 | 0 | 5 | 0 |
| 90 | *BPGM* | Bisphosphoglycerate mutase | 5 | 0 | 2 | 0 | 3 | 0 |
| 91 | *ZNF76* | Zinc finger protein 76 | 5 | 0 | 1 | 1 | 3 | 0 |
| 92 | *ACIN1* | Apoptotic chromatin condensation inducer in the nucleus | 5 | 0 | 1 | 2 | 2 | 0 |
| 93 | *NCKAP5L* | Nck-associated protein 5-like | 5 | 0 | 2 | 1 | 2 | 0 |
| 94 | *SUPT6H* | Transcription elongation factor SPT6 | 5 | 0 | 3 | 0 | 2 | 0 |
| 95 | *KCNA2* | Potassium voltage-gated channel subfamily A member 2 | 5 | 0 | 2 | 0 | 3 | 0 |
| 96 | *WASF2* | Wiskott-Aldrich syndrome protein family member 2 | 6 | 0 | 4 | 1 | 0 | 1 |
| 97 | *SORT1* | Sortilin | 6 | 0 | 2 | 0 | 4 | 0 |
| 98 | *CEP85* | Centrosomal protein of 85 kDa | 5 | 0 | 2 | 2 | 0 | 1 |
| 99 | *FBXO32* | F-box only protein 32 | 5 | 0 | 2 | 2 | 1 | 0 |
| 100 | *HKDC1* | Putative hexokinase HKDC1 | 5 | 0 | 2 | 2 | 1 | 0 |
| 101 | *PRMT7* | Protein arginine N-methyltransferase 7 | 5 | 0 | 2 | 2 | 1 | 0 |
| 102 | *MAN2B2* | Epididymis-specific alpha-mannosidase | 5 | 0 | 2 | 1 | 1 | 1 |
| 103 | *RCN1* | Reticulocalbin-1 | 10 | 0 | 5 | 4 | 1 | 0 |
| 104 | *RNF175* | Ring finger protein 175 | 5 | 0 | 2 | 1 | 2 | 0 |
| 105 | *TGFB2* | Transforming growth factor beta-2 | 6 | 0 | 2 | 0 | 3 | 1 |
| 106 | *PPARGC1A* | Peroxisome proliferator-activated receptor gamma coactivator 1-alpha | 7 | 0 | 1 | 1 | 5 | 0 |
| 107 | *LRTM1* | Leucine rich repeats and transmembrane domains 1 | 5 | 0 | 2 | 2 | 1 | 0 |
| 108 | *FAM222B* | Protein FAM222B | 6 | 0 | 3 | 0 | 3 | 0 |
| 109 | *ETV1* | ETS translocation variant 1 | 6 | 1 | 4 | 0 | 1 | 0 |
| 110 | *AC087477.1* | Putative uncharacterized protein | 4 | 0 | 3 | 0 | 1 | 0 |
| 111 | *AL355531.2* | Putative uncharacterized protein | 4 | 0 | 1 | 2 | 1 | 0 |
| 112 | *NEUROD2* | Neurogenic differentiation factor 2 | 4 | 0 | 2 | 0 | 2 | 0 |
| 113 | *OR10A7* | Olfactory receptor 10A7 | 4 | 0 | 2 | 1 | 1 | 0 |
| 114 | *OR4A15* | Olfactory receptor 4A15 | 4 | 0 | 2 | 0 | 1 | 1 |

**table 7: Annotation of candidate genes' associated NCCMs with data from the ENCODE portal and UCSC browser databases**

| Index | Mutation Name | Mutation Name *(Shared Loci separated by ',' and colored)* | Top NCCM Gene | ICGC Donor ID | Gender | Age at diagnosis | Molecular classification for the tumor | Chrom# | Posi. | Ref. | Alt. | Mutation Classification | phyloP | ORegAnno | Cosnv. TFBS | CpG Islands | DNAse Clusters | Genhancers | H3K4me3 | MeDIP RawSignal | RNA RawSignal | Vista Enhancers | DHS | DRM | Promoter | TF | TSS | Ultra-conserved site | Ultra-sensitive site | UTR | No. of regulatory annotations for the NCCM | CRISPR candidates |
|---|---|---|---|---|---|---|---|---|---|---|---|---|---|---|---|---|---|---|---|---|---|---|---|---|---|---|---|---|---|---|---|---|
| 68 | FGR.NCCM-1 | FGR.NCCM-1,AHDC1.NCCM-1,WASF2.N | *FGR* | DO48911 | female | 23 | SHH | 1 | 27848112 | G | C | IGR | 1,4 | ORE | *nd* | *nd* | DNa | GH0 | H3K | MeDIP-seq Raw Signal | *nd* | *nd* | *nd* | *nd* | *nd* | *nd* | *nd* | *nd* | *nd* | *nd* | 5 | |
| 69 | FGR.NCCM-2 | FGR.NCCM-2,AHDC1.NCCM-2,WASF2.N | *FGR* | DO48916 | female | 25 | Unknown | 1 | 27855405 | CCC | C | RNA | 8,8 | ORE | V--H | *nd* | DNa | GH0 | H3K | MeDIP-seq Raw Signal | *nd* | *nd* | DHS | *nd* | *nd* | TFP. | *nd* | *nd* | *nd* | *nd* | 8 | |
| 70 | FGR.NCCM-3 | FGR.NCCM-3,AHDC1.NCCM-3,WASF2.N | *FGR* | DO35620 | male | 2 | SHH | 1 | 27858637 | C | T | IGR | 2,7 | *nd* | *nd* | *nd* | *nd* | *nd* | *nd* | MeDIP-seq Raw Signal | RNA | *nd* | *nd* | *nd* | *nd* | *nd* | *nd* | *nd* | *nd* | *nd* | 2 | |
| 71 | FGR.NCCM-4 | FGR.NCCM-4,AHDC1.NCCM-4,WASF2.N | *FGR* | DO48891 | female | 32 | SHH | 1 | 27868644 | C | T | Intron | 1,5 | ORE | *nd* | *nd* | DNa | *nd* | *nd* | *nd* | RNA | *nd* | DHS | *nd* | *nd* | TFP. | *nd* | *nd* | *nd* | *nd* | 5 | |
| 72 | FGR.NCCM-5 | FGR.NCCM-5,AHDC1.NCCM-5,WASF2.N | *FGR* | DO35628 | male | 8 | Group 3 | 1 | 27896335 | C | T | Intron | 2,4 | ORE | *nd* | *nd* | DNa | GH0 | *nd* | MeDIP-seq Raw Signal | *nd* | *nd* | DHS | *nd* | *nd* | TFP. | *nd* | *nd* | *nd* | *nd* | 6 | |
| 73 | FGR.NCCM-6 | FGR.NCCM-6,AHDC1.NCCM-6,WASF2.N | *FGR* | DO48907 | male | 31 | SHH | 1 | 27916524 | C | A | Intron | 2,1 | *nd* | V--A | *nd* | DNa | GH0 | H3K | *nd* | *nd* | *nd* | DHS | *nd* | *nd* | TFP. | *nd* | *nd* | *nd* | *nd* | 6 | |
| 121 | UBXN11.NCCM-1 | UBXN11.NCCM-1,SH3BGRL3.NCCM-1,( | *UBXN11* | DO48916 | female | 25 | Unknown | 1 | 26512473 | G | A | Intron | 1,2 | *nd* | *nd* | *nd* | DNa | *nd* | *nd* | MeDIP-seq Raw Signal | *nd* | *nd* | DHS | *nd* | *nd* | *nd* | *nd* | *nd* | *nd* | *nd* | 3 | |
| 122 | UBXN11.NCCM-2 | UBXN11.NCCM-2,SH3BGRL3.NCCM-2,( | *UBXN11* | DO48889 | female | 42 | SHH | 1 | 26597730 | C | T | Intron | 1,3 | *nd* | *nd* | *nd* | *nd* | *nd* | *nd* | MeDIP-seq Raw Signal | RNA | *nd* | *nd* | *nd* | *nd* | *nd* | *nd* | *nd* | *nd* | *nd* | 2 | |
| 123 | UBXN11.NCCM-3 | UBXN11.NCCM-3,SH3BGRL3.NCCM-3,( | *UBXN11* | DO35654 | male | 16 | Group 3 | 1 | 26599878 | C | T | Intron | 1,2 | *nd* | *nd* | *nd* | *nd* | *nd* | H3K | MeDIP-seq Raw Signal | *nd* | *nd* | *nd* | *nd* | *nd* | *nd* | *nd* | *nd* | *nd* | *nd* | 2 | |
| 124 | UBXN11.NCCM-4 | UBXN11.NCCM-4,SH3BGRL3.NCCM-4,( | *UBXN11* | DO35704 | male | 3 | Group 3 | 1 | 26663554 | C | A | Intron | 1,7 | ORE | *nd* | *nd* | *nd* | *nd* | *nd* | MeDIP-seq Raw Signal | *nd* | *nd* | DHS | *nd* | *nd* | TFP. | *nd* | *nd* | *nd* | *nd* | 4 | |
| 125 | UBXN11.NCCM-5 | UBXN11.NCCM-5,SH3BGRL3.NCCM-5,( | *UBXN11* | DO48914 | female | 19 | SHH | 1 | 26678633 | C | A | IGR | 8,9 | *nd* | V--P | *nd* | DNa | GH0 | H3K | MeDIP-seq Raw Signal | *nd* | *nd* | *nd* | *nd* | *nd* | TFP. | *nd* | *nd* | *nd* | *nd* | 6 | |
| 126 | UBXN11.NCCM-6 | UBXN11.NCCM-6,CD52.NCCM-5,ZNF68 | *UBXN11* | DO48909 | female | 39 | SHH | 1 | 26715523 | G | T | IGR | 1,8 | *nd* | *nd* | *nd* | DNa | *nd* | *nd* | MeDIP-seq Raw Signal | *nd* | *nd* | DHS | *nd* | *nd* | TFP. | *nd* | *nd* | *nd* | *nd* | 4 | |
| 186 | KCNA3.NCCM-1 | KCNA3.NCCM-1,KCNA2.NCCM-2 | *KCNA3* | DO35749 | male | 16 | Group 4 | 1 | 111125152 | G | A | IGR | 2,8 | *nd* | *nd* | *nd* | DNa | *nd* | H3K | *nd* | *nd* | *nd* | DHS | *nd* | *nd* | *nd* | *nd* | *nd* | *nd* | *nd* | 3 | |
| 187 | KCNA3.NCCM-2 | KCNA3.NCCM-2,KCNA2.NCCM-3 | *KCNA3* | DO35554 | female | 13 | Group 4 | 1 | 111157994 | C | A | Intron | 1,4 | *nd* | *nd* | *nd* | *nd* | *nd* | *nd* | MeDIP-seq Raw Signal | *nd* | *nd* | *nd* | *nd* | *nd* | *nd* | *nd* | *nd* | *nd* | *nd* | 1 | |
| 188 | KCNA3.NCCM-3 | KCNA3.NCCM-3,KCNA2.NCCM-4 | *KCNA3* | DO48914 | female | 19 | SHH | 1 | 111188148 | C | T | IGR | 1,2 | *nd* | *nd* | *nd* | DNa | *nd* | *nd* | *nd* | *nd* | *nd* | DHS | *nd* | *nd* | TFP. | *nd* | *nd* | *nd* | *nd* | 3 | |
| 189 | KCNA3.NCCM-4 | KCNA3.NCCM-4,KCNA2.NCCM-5 | *KCNA3* | DO48906 | female | 49 | SHH | 1 | 111228843 | ACC | A | IGR | 2,8 | ORE | *nd* | *nd* | *nd* | *nd* | H3K | MeDIP-seq Raw Signal | RNA | *nd* | *nd* | *nd* | *nd* | TFP. | *nd* | *nd* | *nd* | *nd* | 5 | |
| 190 | KCNA3.NCCM-5 | KCNA3.NCCM-5 | *KCNA3* | DO48906 | female | 49 | SHH | 1 | 111302196 | CTT | C | IGR | 1,8 | *nd* | *nd* | *nd* | *nd* | *nd* | *nd* | MeDIP-seq Raw Signal | *nd* | *nd* | *nd* | *nd* | *nd* | *nd* | *nd* | *nd* | *nd* | *nd* | 1 | |
| 206 | HMGB4.NCCM-1 | HMGB4.NCCM-1 | *HMGB4* | DO48953 | female | 8 | Group 4 | 1 | 34338362 | C | T | Intron | 1,4 | *nd* | *nd* | *nd* | DNa | *nd* | *nd* | MeDIP-seq Raw Signal | *nd* | *nd* | DHS | *nd* | *nd* | *nd* | *nd* | *nd* | *nd* | *nd* | 3 | |
| 207 | HMGB4.NCCM-2 | HMGB4.NCCM-2 | *HMGB4* | DO35628 | male | 8 | Group 3 | 1 | 34363877 | A | G | Intron | 6,3 | *nd* | *nd* | *nd* | *nd* | *nd* | *nd* | MeDIP-seq Raw Signal | *nd* | *nd* | *nd* | *nd* | *nd* | *nd* | *nd* | *nd* | *nd* | *nd* | 1 | |
| 208 | HMGB4.NCCM-3 | HMGB4.NCCM-3 | *HMGB4* | DO48891 | female | 32 | SHH | 1 | 34364073 | T | C | Intron | 4,7 | *nd* | V--C | *nd* | *nd* | *nd* | *nd* | *nd* | *nd* | *nd* | *nd* | *nd* | *nd* | *nd* | *nd* | *nd* | *nd* | *nd* | 1 | |
| 209 | HMGB4.NCCM-4 | HMGB4.NCCM-4 | *HMGB4* | DO48902 | female | 29 | SHH | 1 | 34368229 | C | T | Intron | 3,3 | *nd* | *nd* | *nd* | DNa | *nd* | *nd* | MeDIP-seq Raw Signal | *nd* | *nd* | DHS | *nd* | *nd* | TFP. | *nd* | *nd* | *nd* | *nd* | 4 | |
| 210 | HMGB4.NCCM-5 | HMGB4.NCCM-5 | *HMGB4* | DO35841 | male | 16 | Group 4 | 1 | 34402804 | G | C | Intron | 1,8 | *nd* | *nd* | *nd* | *nd* | *nd* | *nd* | MeDIP-seq Raw Signal | *nd* | *nd* | *nd* | *nd* | *nd* | *nd* | *nd* | *nd* | *nd* | *nd* | 1 | |
| 211 | SH3BGRL3.NCCM-1 | UBXN11.NCCM-1,SH3BGRL3.NCCM-1,( | *SH3BGRL3* | DO48916 | female | 25 | Unknown | 1 | 26512473 | G | A | Intron | 1,2 | *nd* | *nd* | *nd* | DNa | *nd* | *nd* | MeDIP-seq Raw Signal | *nd* | *nd* | DHS | *nd* | *nd* | *nd* | *nd* | *nd* | *nd* | *nd* | 3 | |
| 212 | SH3BGRL3.NCCM-2 | UBXN11.NCCM-2,SH3BGRL3.NCCM-2,( | *SH3BGRL3* | DO48889 | female | 42 | SHH | 1 | 26597730 | C | T | Intron | 1,3 | *nd* | *nd* | *nd* | *nd* | *nd* | *nd* | MeDIP-seq Raw Signal | RNA | *nd* | *nd* | *nd* | *nd* | *nd* | *nd* | *nd* | *nd* | *nd* | 2 | |
| 213 | SH3BGRL3.NCCM-3 | UBXN11.NCCM-3,SH3BGRL3.NCCM-3,( | *SH3BGRL3* | DO35654 | male | 16 | Group 3 | 1 | 26599878 | C | T | Intron | 1,2 | *nd* | *nd* | *nd* | *nd* | *nd* | H3K | MeDIP-seq Raw Signal | *nd* | *nd* | *nd* | *nd* | *nd* | *nd* | *nd* | *nd* | *nd* | *nd* | 2 | |
| 214 | SH3BGRL3.NCCM-4 | UBXN11.NCCM-4,SH3BGRL3.NCCM-4,( | *SH3BGRL3* | DO35704 | male | 3 | Group 3 | 1 | 26663554 | C | A | Intron | 1,7 | ORE | *nd* | *nd* | *nd* | *nd* | *nd* | MeDIP-seq Raw Signal | *nd* | *nd* | DHS | *nd* | *nd* | TFP. | *nd* | *nd* | *nd* | *nd* | 4 | |
| 215 | SH3BGRL3.NCCM-5 | UBXN11.NCCM-5,SH3BGRL3.NCCM-5,( | *SH3BGRL3* | DO48914 | female | 19 | SHH | 1 | 26678633 | C | A | IGR | 8,9 | *nd* | V--P | *nd* | DNa | GH0 | H3K | MeDIP-seq Raw Signal | *nd* | *nd* | *nd* | *nd* | *nd* | TFP. | *nd* | *nd* | *nd* | *nd* | 6 | |
| 216 | CD52.NCCM-1 | UBXN11.NCCM-2,SH3BGRL3.NCCM-2,( | *CD52* | DO48889 | female | 42 | SHH | 1 | 26597730 | C | T | Intron | 1,3 | *nd* | *nd* | *nd* | *nd* | *nd* | *nd* | MeDIP-seq Raw Signal | RNA | *nd* | *nd* | *nd* | *nd* | *nd* | *nd* | *nd* | *nd* | *nd* | 2 | |
| 217 | CD52.NCCM-2 | UBXN11.NCCM-3,SH3BGRL3.NCCM-3,( | *CD52* | DO35654 | male | 16 | Group 3 | 1 | 26599878 | C | T | Intron | 1,2 | *nd* | *nd* | *nd* | *nd* | *nd* | H3K | MeDIP-seq Raw Signal | *nd* | *nd* | *nd* | *nd* | *nd* | *nd* | *nd* | *nd* | *nd* | *nd* | 2 | |
| 218 | CD52.NCCM-3 | UBXN11.NCCM-4,SH3BGRL3.NCCM-4,( | *CD52* | DO35704 | male | 3 | Group 3 | 1 | 26663554 | C | A | Intron | 1,7 | ORE | *nd* | *nd* | *nd* | *nd* | *nd* | MeDIP-seq Raw Signal | *nd* | *nd* | DHS | *nd* | *nd* | TFP. | *nd* | *nd* | *nd* | *nd* | 4 | |
| 219 | CD52.NCCM-4 | UBXN11.NCCM-5,SH3BGRL3.NCCM-5,( | *CD52* | DO48914 | female | 19 | SHH | 1 | 26678633 | C | A | IGR | 8,9 | *nd* | V--P | *nd* | DNa | GH0 | H3K | MeDIP-seq Raw Signal | *nd* | *nd* | *nd* | *nd* | *nd* | TFP. | *nd* | *nd* | *nd* | *nd* | 6 | |
| 220 | CD52.NCCM-5 | UBXN11.NCCM-6,CD52.NCCM-5,ZNF68 | *CD52* | DO48909 | female | 39 | SHH | 1 | 26715523 | G | T | IGR | 1,8 | *nd* | *nd* | *nd* | DNa | *nd* | *nd* | MeDIP-seq Raw Signal | *nd* | *nd* | DHS | *nd* | *nd* | TFP. | *nd* | *nd* | *nd* | *nd* | 4 | |
| 276 | ZBTB18.NCCM-1 | ZBTB18.NCCM-1 | *ZBTB18* | DO35628 | male | 8 | Group 3 | 1 | 244118526 | G | C | lincRNA | 1,4 | *nd* | *nd* | *nd* | *nd* | *nd* | *nd* | *nd* | *nd* | *nd* | *nd* | *nd* | *nd* | *nd* | *nd* | *nd* | *nd* | *nd* | 0 | |
| 277 | ZBTB18.NCCM-2 | ZBTB18.NCCM-2 | *ZBTB18* | DO35749 | male | 16 | Group 4 | 1 | 244150884 | G | C | lincRNA | 2,4 | *nd* | *nd* | *nd* | *nd* | *nd* | H3K | MeDIP-seq Raw Signal | *nd* | *nd* | *nd* | *nd* | *nd* | *nd* | *nd* | *nd* | *nd* | *nd* | 2 | |
| 278 | ZBTB18.NCCM-3 | ZBTB18.NCCM-3 | *ZBTB18* | DO35625 | female | 15 | Group 4 | 1 | 244203195 | A | G | lincRNA | 1,6 | *nd* | *nd* | *nd* | *nd* | *nd* | *nd* | MeDIP-seq Raw Signal | *nd* | *nd* | *nd* | *nd* | *nd* | *nd* | *nd* | *nd* | *nd* | *nd* | 1 | |
| 279 | ZBTB18.NCCM-4 | ZBTB18.NCCM-4 | *ZBTB18* | DO48906 | female | 49 | SHH | 1 | 244217248 | TTC | G | NOINFO | 8,9 | *nd* | *nd* | *nd* | DNa | *nd* | H3K | MeDIP-seq Raw Signal | RNA | *nd* | DHS | *nd* | *nd* | *nd* | *nd* | *nd* | *nd* | *nd* | 5 | |
| 280 | ZBTB18.NCCM-5 | ZBTB18.NCCM-5 | *ZBTB18* | DO35582 | female | 6 | WNT | 1 | 244219476 | A | G | 3'UTR | 6,3 | *nd* | *nd* | *nd* | *nd* | *nd* | *nd* | MeDIP-seq Raw Signal | RNA | *nd* | *nd* | *nd* | *nd* | *nd* | *nd* | *nd* | *nd* | UTR | 3 | |
| 311 | ZC3H12A.NCCM-1 | ZC3H12A.NCCM-1 | *ZC3H12A* | DO35625 | female | 15 | Group 4 | 1 | 37852870 | G | GT | IGR | 1,2 | *nd* | *nd* | *nd* | DNa | *nd* | *nd* | MeDIP-seq Raw Signal | *nd* | *nd* | DHS | *nd* | *nd* | *nd* | *nd* | *nd* | *nd* | *nd* | 3 | |
| 312 | ZC3H12A.NCCM-2 | ZC3H12A.NCCM-2 | *ZC3H12A* | DO48911 | female | 23 | SHH | 1 | 37866483 | C | T | IGR | 5,9 | *nd* | *nd* | *nd* | *nd* | *nd* | H3K | MeDIP-seq Raw Signal | *nd* | *nd* | *nd* | *nd* | *nd* | *nd* | *nd* | *nd* | *nd* | *nd* | 2 | |
| 313 | ZC3H12A.NCCM-3 | ZC3H12A.NCCM-3 | *ZC3H12A* | DO48888 | male | 20 | SHH | 1 | 37870536 | C | A | IGR | 4,9 | *nd* | *nd* | *nd* | DNa | *nd* | H3K | MeDIP-seq Raw Signal | *nd* | *nd* | DHS | *nd* | *nd* | *nd* | *nd* | *nd* | *nd* | *nd* | 4 | |
| 314 | ZC3H12A.NCCM-4 | ZC3H12A.NCCM-4 | *ZC3H12A* | DO35572 | female | 4 | SHH | 1 | 37883902 | G | A | IGR | 1,2 | *nd* | *nd* | *nd* | *nd* | *nd* | *nd* | MeDIP-seq Raw Signal | *nd* | *nd* | *nd* | *nd* | *nd* | *nd* | *nd* | *nd* | *nd* | *nd* | 1 | |
| 315 | ZC3H12A.NCCM-5 | ZC3H12A.NCCM-5 | *ZC3H12A* | DO48909 | female | 39 | SHH | 1 | 37915124 | G | T | IGR | 1,7 | *nd* | *nd* | *nd* | *nd* | GH0 | *nd* | MeDIP-seq Raw Signal | *nd* | *nd* | *nd* | *nd* | *nd* | *nd* | *nd* | *nd* | *nd* | *nd* | 2 | |
| 366 | ZNF683.NCCM-1 | UBXN11.NCCM-2,SH3BGRL3.NCCM-2,( | *ZNF683* | DO48889 | female | 42 | SHH | 1 | 26597730 | C | T | Intron | 1,3 | *nd* | *nd* | *nd* | *nd* | *nd* | *nd* | MeDIP-seq Raw Signal | RNA | *nd* | *nd* | *nd* | *nd* | *nd* | *nd* | *nd* | *nd* | *nd* | 2 | |
| 367 | ZNF683.NCCM-2 | UBXN11.NCCM-3,SH3BGRL3.NCCM-3,( | *ZNF683* | DO35654 | male | 16 | Group 3 | 1 | 26599878 | C | T | Intron | 1,2 | *nd* | *nd* | *nd* | *nd* | *nd* | H3K | MeDIP-seq Raw Signal | *nd* | *nd* | *nd* | *nd* | *nd* | *nd* | *nd* | *nd* | *nd* | *nd* | 2 | |
| 368 | ZNF683.NCCM-3 | UBXN11.NCCM-4,SH3BGRL3.NCCM-4,( | *ZNF683* | DO35704 | male | 3 | Group 3 | 1 | 26663554 | C | A | Intron | 1,7 | ORE | *nd* | *nd* | *nd* | *nd* | *nd* | MeDIP-seq Raw Signal | *nd* | *nd* | DHS | *nd* | *nd* | TFP. | *nd* | *nd* | *nd* | *nd* | 4 | |
| 369 | ZNF683.NCCM-4 | UBXN11.NCCM-5,SH3BGRL3.NCCM-5,( | *ZNF683* | DO48914 | female | 19 | SHH | 1 | 26678633 | C | A | IGR | 8,9 | *nd* | V--P | *nd* | DNa | GH0 | H3K | MeDIP-seq Raw Signal | *nd* | *nd* | *nd* | *nd* | *nd* | TFP. | *nd* | *nd* | *nd* | *nd* | 6 | |
| 370 | ZNF683.NCCM-5 | UBXN11.NCCM-6,CD52.NCCM-5,ZNF68 | *ZNF683* | DO48909 | female | 39 | SHH | 1 | 26715523 | G | T | IGR | 1,8 | *nd* | *nd* | *nd* | DNa | *nd* | *nd* | MeDIP-seq Raw Signal | *nd* | *nd* | DHS | *nd* | *nd* | TFP. | *nd* | *nd* | *nd* | *nd* | 4 | |
| 376 | PGLYRP3.NCCM-1 | PGLYRP3.NCCM-1,PGLYRP4.NCCM-1 | *PGLYRP3* | DO48909 | female | 39 | SHH | 1 | 153211766 | C | T | IGR | 3,0 | *nd* | *nd* | *nd* | DNa | *nd* | *nd* | MeDIP-seq Raw Signal | *nd* | *nd* | *nd* | *nd* | *nd* | *nd* | *nd* | *nd* | *nd* | *nd* | 2 | |
| 377 | PGLYRP3.NCCM-2 | PGLYRP3.NCCM-2,PGLYRP4.NCCM-2 | *PGLYRP3* | DO35704 | male | 3 | Group 3 | 1 | 153225416 | G | A | IGR | 3,4 | *nd* | *nd* | *nd* | *nd* | *nd* | *nd* | MeDIP-seq Raw Signal | *nd* | *nd* | *nd* | *nd* | *nd* | *nd* | *nd* | *nd* | *nd* | *nd* | 1 | |
| 378 | PGLYRP3.NCCM-3 | PGLYRP3.NCCM-3,PGLYRP4.NCCM-3 | *PGLYRP3* | DO48902 | female | 29 | SHH | 1 | 153318866 | C | A | Intron | 1,4 | *nd* | *nd* | *nd* | DNa | *nd* | *nd* | MeDIP-seq Raw Signal | *nd* | *nd* | DHS | *nd* | *nd* | TFP. | *nd* | *nd* | *nd* | *nd* | 4 | |
| 379 | PGLYRP3.NCCM-4 | PGLYRP3.NCCM-4,PGLYRP4.NCCM-4 | *PGLYRP3* | DO48909 | female | 39 | SHH | 1 | 153328122 | C | T | 5'Flank | 2,4 | ORE | *nd* | *nd* | DNa | GH0 | *nd* | MeDIP-seq Raw Signal | *nd* | *nd* | DHS | *nd* | Pror | TFP. | *nd* | *nd* | *nd* | *nd* | 7 | |
| 380 | PGLYRP3.NCCM-5 | PGLYRP3.NCCM-5,PGLYRP4.NCCM-5 | *PGLYRP3* | DO48966 | female | 3 | Group 4 | 1 | 153359242 | G | C | IGR | 1,7 | ORE | *nd* | *nd* | DNa | GH0 | *nd* | MeDIP-seq Raw Signal | *nd* | *nd* | DHS | *nd* | *nd* | TFP. | *nd* | *nd* | *nd* | *nd* | 6 | |
| [illegible] | [illegible] | [illegible] | [illegible] | [illegible] | [illegible] | [illegible] | [illegible] | [illegible] | [illegible] | [illegible] | [illegible] | [illegible] | [illegible] | [illegible] | [illegible] | [illegible] | [illegible] | [illegible] | [illegible] | [illegible] | [illegible] | [illegible] | [illegible] | [illegible] | [illegible] | [illegible] | [illegible] | [illegible] | [illegible] | [illegible] | [illegible] | [illegible] |

**Key for table 7.**

| Col | Name | Description |
|---|---|---|
| A | Index | Sequential indexing for the current dataset |
| B | Mutation Name | Name of the mutation, formed by prefixing gene + consecutive |
| C | Mutation Name (Shared Loci separated by ',' and colored) | If present, other shared loci are shown as well. Otherwise nam |
| D | Top NCCM Gene | Genes, whose regulatory potential is examined |
| E | ICGC Donor ID | ICGC Sample ID |
| F | Molecular classification for the tumor | Can be one of the four categories listed in the key |
| G | Gender | Gender/Sex for the patient |
| H | Age at diagnosis | Age at which the disease was diagnosed. Colored cells represe |
| I | Chrom# | Chromosome number |
| J | Posi. | Position(s) where SPM/SIM is observed |
| K | Ref. | Reference allele |
| L | Alt. | Alternate allele |
| M | Mutation Classification | Type of change for the SPM/SIM. |
| N | phyloP | phyloP scores. |
| O | ORegAnno | SPM/SIM mapped to a regulatory region, which come from a c |
| P | Cosnv. TFBS | SPM/SIM mapped mapped to a conserved transcription factor |
| Q | CpG Islands | CpG islands are typically common near transcription start sites |
| R | DNAse Clusters | Digital DNaseI Hypersensitivity Clusters from ENCODE -- DNase |
| S | Genhancers | GeneHancer is a database of human regulatory elements (enh |
| T | H3K4me3 | H3K4Me3 track shows a histone mark associated with promot |
| U | MeDIP RawSignal | MeDIP-seq uses immunoprecipitation to extract the methylate |
| V | RNA RawSignal | Transcription Levels Assayed by RNA-seq. |
| W | Vista Enhancers | Enhancer sequences are regulatory DNA sequences that, wher |
| X | DHS | Dnase I hypersensitive sites derived from the ENCODE project. |
| Y | DRM | Distal regulatory module enhancers. These regulate the expres |
| Z | Promoter | Promoter annotations derived from GENCODE v16. |
| AA | TF | Transcription factor binding sites derived from the ENCODE pr |
| AB | TSS | Transcription start sites derived by taking the 100bp intervals i |
| AC | Ultra-conserved site | Sites that are highly conserved across many phylogenetic bran |
| AD | Ultra-sensitive site | Sites that are under higher selective pressure than most other |
| AE | UTR | UTR annotations derived from GENCODE v16. |
| AF | No. of regulatory annotations for the NCCM | Total number of regulatory annotations for the NCCM |

table 8: Genes in MB with ≥ 5 NCCMs in the target area (±100

| Hugo Gene | NCCM Counts | Rates/100 kbp (Cells in yellow have NCCMs ≥ 1.8 & < 2.0) | Genes with NCCMs ≥ 2/100 kbp? Y/N |
|---|---|---|---|
| DMD | 19 | 0,78 | No |
| ARHGAP15 | 16 | 1,83 | No |
| AUTS2 | 16 | 1,15 | No |
| IL1RAPL1 | 16 | 1,02 | No |
| CADM2 | 15 | 1,14 | No |
| C10orf11 | 15 | 1,29 | No |
| DLG2 | 15 | 0,63 | No |
| PARD3B | 14 | 1,10 | No |
| MECOM | 14 | 1,80 | No |
| TENM2 | 14 | 1,20 | No |
| SOX6 | 14 | 1,44 | No |
| ENOX1 | 14 | 1,82 | No |
| PTPRT | 14 | 1,07 | No |
| LSAMP | 13 | 0,54 | No |
| KCND2 | 13 | 1,93 | No |
| NPAS3 | 13 | 1,22 | No |
| DCC | 13 | 0,94 | No |
| LRP1B | 12 | 0,57 | No |
| EYS | 12 | 0,55 | No |
| NEGR1 | 11 | 1,01 | No |
| DPP10 | 11 | 0,69 | No |
| ROBO2 | 11 | 0,57 | No |
| GRID2 | 11 | 0,66 | No |
| SLIT3 | 11 | 1,32 | No |
| HDAC9 | 11 | 0,99 | No |
| IMMP2L | 11 | 1,00 | No |
| CNTNAP2 | 11 | 0,44 | No |
| CNTN5 | 11 | 0,72 | No |
| OPCML | 11 | 0,84 | No |
| HS6ST3 | 11 | 1,17 | No |
| NRXN3 | 11 | 0,61 | No |
| RBFOX1 | 11 | 0,58 | No |
| MACROD2 | 11 | 0,49 | No |
| CSMD2 | 10 | 1,19 | No |
| DAB1 | 10 | 0,57 | No |
| ESRRG | 10 | 1,20 | No |
| SLC4A10 | 10 | 1,32 | No |
| TBC1D5 | 10 | 0,67 | No |
| CACNA2D3 | 10 | 0,87 | No |
| FHIT | 10 | 0,59 | No |
| ZBTB20 | 10 | 0,99 | No |
| NLGN1 | 10 | 0,92 | No |
| CCSER1 | 10 | 0,60 | No |
| TENM3 | 10 | 1,18 | No |
| EBF1 | 10 | 1,66 | No |
| PTPRD | 10 | 0,40 | No |
| DENND1A | 10 | 1,34 | No |
| PRKG1 | 10 | 0,66 | No |
| RCN1 | 10 | 2,03 | Yes |
| TENM4 | 10 | 1,02 | No |
| SYT1 | 10 | 1,27 | No |
| ANKS1B | 10 | 0,69 | No |
| GPC6 | 10 | 0,73 | No |
| RYR3 | 10 | 1,35 | No |
| MSI2 | 10 | 1,61 | No |
| ZNF536 | 10 | 1,47 | No |
| LPHN2 | 9 | 1,02 | No |
| FOXP1 | 9 | 1,09 | No |
| ROBO1 | 9 | 0,66 | No |
| LIMCH1 | 9 | 1,68 | No |
| CDH12 | 9 | 0,69 | No |
| ARL15 | 9 | 1,44 | No |
| MEF2C | 9 | 2,34 | Yes |
| FAM172A | 9 | 1,30 | No |
| EFNA5 | 9 | 1,82 | No |
| SGCD | 9 | 0,82 | No |
| OFCC1 | 9 | 1,11 | No |
| DOCK4 | 9 | 1,33 | No |
| ZFHX4 | 9 | 2,40 | Yes |
| CSMD3 | 9 | 0,64 | No |
| ZNF462 | 9 | 2,63 | Yes |
| PCDH15 | 9 | 0,45 | No |
| CTNNA3 | 9 | 0,45 | No |
| NRG3 | 9 | 0,69 | No |
| NAV2 | 9 | 0,93 | No |
| LRRC4C | 9 | 0,58 | No |
| KIRREL3 | 9 | 1,16 | No |
| CACNA1C | 9 | 0,98 | No |
| TRHDE | 9 | 1,16 | No |
| SEMA6D | 9 | 1,14 | No |
| SKAP1 | 9 | 1,82 | No |
| AGBL4 | 8 | 0,47 | No |
| ST6GALNAC3 | 8 | 1,05 | No |
| DPYD | 8 | 0,77 | No |
| BRE | 8 | 1,24 | No |
| NRXN1 | 8 | 0,61 | No |
| SPTBN1 | 8 | 1,97 | No |
| CTNNA2 | 8 | 0,48 | No |
| IFIH1 | 8 | 3,22 | Yes |
| NABP1 | 8 | 3,67 | Yes |
| CADPS | 8 | 1,19 | No |
| FIP1L1 | 8 | 0,72 | No |
| LPHN3 | 8 | 0,75 | No |
| SLC4A4 | 8 | 1,39 | No |
| SLC10A7 | 8 | 1,71 | No |
| SUGCT | 8 | 0,87 | No |
| GRM8 | 8 | 0,79 | No |
| DPP6 | 8 | 0,62 | No |
| SGCZ | 8 | 0,59 | No |
| NRG1 | 8 | 0,60 | No |
| SAMD12 | 8 | 1,27 | No |
| PRUNE2 | 8 | 1,65 | No |
| ASTN2 | 8 | 0,67 | No |
| PBX3 | 8 | 1,91 | No |
| NEBL | 8 | 1,35 | No |
| GRIP1 | 8 | 1,22 | No |
| PCDH9 | 8 | 0,71 | No |
| DACH1 | 8 | 1,28 | No |
| TTC6 | 8 | 1,25 | No |
| MDGA2 | 8 | 0,78 | No |
| ZFHX3 | 8 | 1,72 | No |
| ANKFN1 | 8 | 1,34 | No |
| BCAS3 | 8 | 0,88 | No |
| TM4SF2 | 8 | 0,52 | No |
| DIAPH2 | 8 | 0,72 | No |
| PRDM16 | 7 | 1,24 | No |
| GRIK3 | 7 | 1,61 | No |
| PDE4B | 7 | 0,90 | No |
| RP5-1052I5.2 | 7 | 1,87 | No |
| NCKAP5 | 7 | 0,64 | No |
| UPP2 | 7 | 1,53 | No |
| ERBB4 | 7 | 0,52 | No |
| CNTN4 | 7 | 0,61 | No |
| ZNF385D | 7 | 0,61 | No |
| THRB | 7 | 1,21 | No |
| RARB | 7 | 1,12 | No |
| ERC2 | 7 | 0,61 | No |
| SYNPR | 7 | 1,19 | No |
| SLC9A9 | 7 | 0,90 | No |
| IQCJ-SCHIP1 | 7 | 0,62 | No |
| EIF2B5 | 7 | 0,94 | No |
| LPP | 7 | 0,75 | No |
| PPARGC1A | 7 | 2,02 | Yes |
| COL25A1 | 7 | 1,02 | No |
| MAST4 | 7 | 0,91 | No |
| MCTP1 | 7 | 0,90 | No |
| ANKS1A | 7 | 1,76 | No |
| RUNX2 | 7 | 1,31 | No |
| BAI3 | 7 | 0,74 | No |
| MAP3K7 | 7 | 2,58 | Yes |
| HS3ST5 | 7 | 1,44 | No |
| EPB41L2 | 7 | 1,67 | No |
| ESR1 | 7 | 1,04 | No |
| PARK2 | 7 | 0,44 | No |
| SDK1 | 7 | 0,60 | No |
| CALN1 | 7 | 0,81 | No |
| COL1A2 | 7 | 3,03 | Yes |
| CASD1 | 7 | 2,85 | Yes |
| FOXP2 | 7 | 0,88 | No |
| EXOC4 | 7 | 0,69 | No |
| EBF2 | 7 | 1,74 | No |
| VPS13B | 7 | 0,67 | No |
| PCSK5 | 7 | 1,05 | No |
| FSD1L | 7 | 2,31 | Yes |
| MAPKAP1 | 7 | 1,50 | No |
| ARMC4 | 7 | 1,66 | No |
| ZNF365 | 7 | 1,41 | No |
| TCF7L2 | 7 | 1,68 | No |
| NELL1 | 7 | 0,63 | No |
| ELP4 | 7 | 1,48 | No |
| PAX6 | 7 | 3,02 | Yes |
| GRIA4 | 7 | 1,24 | No |
| CADM1 | 7 | 1,31 | No |
| SOX5 | 7 | 1,13 | No |
| TPH2 | 7 | 1,57 | No |
| NUAK1 | 7 | 2,55 | Yes |
| GPC5 | 7 | 0,42 | No |
| GPHN | 7 | 0,80 | No |
| FMN1 | 7 | 1,12 | No |
| RORA | 7 | 0,74 | No |
| PRKCB | 7 | 1,20 | No |
| ABCC12 | 7 | 2,60 | Yes |
| NTN1 | 7 | 1,66 | No |
| PTPRM | 7 | 0,68 | No |
| ZNF521 | 7 | 1,44 | No |
| FHOD3 | 7 | 1,03 | No |
| RUNX1 | 7 | 0,49 | No |
| UBXN11 | 6 | 2,56 | Yes |
| WASF2 | 6 | 2,11 | Yes |
| AHDC1 | 6 | 2,27 | Yes |
| FGR | 6 | 2,71 | Yes |
| ZMYM4 | 6 | 1,72 | No |
| COL24A1 | 6 | 0,96 | No |
| HS2ST1 | 6 | 1,43 | No |
| SORT1 | 6 | 2,10 | Yes |
| PBX1 | 6 | 1,13 | No |
| DNM3 | 6 | 0,77 | No |
| TNR | 6 | 0,96 | No |
| ASTN1 | 6 | 1,19 | No |
| TGFB2 | 6 | 2,02 | Yes |
| SDCCAG8 | 6 | 1,36 | No |
| GALNT14 | 6 | 1,35 | No |
| EML6 | 6 | 1,36 | No |
| CCDC85A | 6 | 1,50 | No |
| ZEB2 | 6 | 1,79 | No |
| RBMS1 | 6 | 1,44 | No |
| GCA | 6 | 2,38 | Yes |
| OLA1 | 6 | 1,60 | No |
| PLCL1 | 6 | 0,62 | No |
| SPAG16 | 6 | 0,45 | No |
| PID1 | 6 | 0,97 | No |
| DIS3L2 | 6 | 1,04 | No |
| GRM7 | 6 | 0,51 | No |
| PPARG | 6 | 1,74 | No |
| ZCWPW2 | 6 | 1,55 | No |
| RBMS3 | 6 | 0,65 | No |
| GTF2E1 | 6 | 2,51 | Yes |
| EPHB1 | 6 | 0,70 | No |
| RSRC1 | 6 | 0,94 | No |
| FGF12 | 6 | 0,72 | No |
| KCNIP4 | 6 | 0,42 | No |
| EPHA5 | 6 | 1,10 | No |
| FRAS1 | 6 | 0,89 | No |
| SLC7A11 | 6 | 2,17 | Yes |
| HAND2 | 6 | 2,93 | Yes |
| ADAMTS16 | 6 | 1,59 | No |
| LIFR | 6 | 1,82 | No |
| PDE4D | 6 | 0,34 | No |
| PCSK1 | 6 | 2,48 | Yes |
| FBXL17 | 6 | 0,83 | No |
| DIAPH1 | 6 | 2,00 | No |
| ARHGAP26 | 6 | 0,91 | No |
| CDKAL1 | 6 | 0,67 | No |
| KCNQ5 | 6 | 0,78 | No |
| BACH2 | 6 | 1,06 | No |
| MTHFD1L | 6 | 1,39 | No |
| ARID1B | 6 | 0,96 | No |
| RPA3_AltSplice | 6 | 1,07 | No |
| ETV1 | 6 | 2,01 | Yes |
| PDE1C | 6 | 0,80 | No |
| ELMO1 | 6 | 0,76 | No |
| POU6F2 | 6 | 0,84 | No |
| GLI3 | 6 | 1,27 | No |
| MAGI2 | 6 | 0,37 | No |
| DLX6 | 6 | 2,93 | Yes |
| RELN | 6 | 0,85 | No |
| CHCHD3 | 6 | 1,21 | No |
| TMEM178B | 6 | 0,99 | No |
| STK3 | 6 | 0,81 | No |
| RP11-127H5.1 | 6 | 0,58 | No |
| EXT1 | 6 | 1,16 | No |
| BNC2 | 6 | 0,91 | No |
| MLLT3 | 6 | 1,25 | No |
| TLE4 | 6 | 1,70 | No |
| SLC44A1 | 6 | 1,53 | No |
| RP11-508N12.4 | 6 | 2,99 | Yes |
| CRB2 | 6 | 2,72 | Yes |
| ST8SIA6 | 6 | 1,79 | No |
| ANK3 | 6 | 0,67 | No |
| HK1 | 6 | 1,83 | No |
| HPSE2 | 6 | 0,62 | No |
| GFRA1 | 6 | 1,45 | No |
| MGMT | 6 | 1,20 | No |
| TRPC6 | 6 | 0,97 | No |
| NTM | 6 | 0,51 | No |
| ANO2 | 6 | 0,98 | No |
| GRIN2B | 6 | 0,94 | No |
| HMGA2 | 6 | 1,76 | No |
| PTPRQ | 6 | 1,29 | No |
| TMEM132C | 6 | 0,94 | No |
| SAMD4A | 6 | 1,41 | No |
| RAD51B | 6 | 0,54 | No |
| KIAA0247 | 6 | 1,98 | No |
| ALDH1A2 | 6 | 0,81 | No |
| FTO | 6 | 0,97 | No |
| CDH8 | 6 | 1,02 | No |
| CNTNAP4 | 6 | 1,26 | No |
| FAM222B | 6 | 2,02 | Yes |
| NF1 | 6 | 1,25 | No |
| ASIC2 | 6 | 0,44 | No |
| NFE2L1 | 6 | 2,85 | Yes |
| CBX1 | 6 | 2,60 | Yes |
| SNX11 | 6 | 2,75 | Yes |
| AXIN2 | 6 | 2,60 | Yes |
| DLGAP1 | 6 | 0,52 | No |
| KCTD1 | 6 | 1,51 | No |
| SETBP1 | 6 | 1,03 | No |
| NFIX | 6 | 1,99 | No |
| NACC1 | 6 | 2,71 | Yes |
| TASP1 | 6 | 1,05 | No |
| GSS | 6 | 2,66 | Yes |
| DLGAP4 | 6 | 1,31 | No |
| RBFOX2 | 6 | 1,23 | No |
| CACNG2 | 6 | 1,77 | No |
| AMMECR1 | 6 | 1,35 | No |
| TENM1 | 6 | 0,77 | No |
| GPC3 | 6 | 0,93 | No |
| CAMTA1 | 5 | 0,42 | No |
| KAZN | 5 | 0,70 | No |
| CEP85 | 5 | 2,06 | Yes |
| SH3BGRL3 | 5 | 2,47 | Yes |
| CD52 | 5 | 2,47 | Yes |
| AIM1L | 5 | 2,18 | Yes |
| ZNF683 | 5 | 2,37 | Yes |
| HMGB4 | 5 | 2,48 | Yes |
| C1orf94 | 5 | 2,00 | No |
| ZC3H12A | 5 | 2,40 | Yes |
| MACF1 | 5 | 0,86 | No |
| HIVEP3 | 5 | 0,69 | No |
| RNF220 | 5 | 1,13 | No |
| TRABD2B | 5 | 1,15 | No |
| SNX7 | 5 | 1,68 | No |
| KCNA2 | 5 | 2,12 | Yes |
| KCNA3 | 5 | 2,48 | Yes |
| PGLYRP3 | 5 | 2,36 | Yes |
| PGLYRP4 | 5 | 2,30 | Yes |
| BRINP2 | 5 | 1,62 | No |
| CR1 | 5 | 1,48 | No |
| RRP15 | 5 | 1,99 | No |
| ZBTB18 | 5 | 2,44 | Yes |
| NBAS | 5 | 0,85 | No |
| AC008271.1 | 5 | 1,99 | No |
| FOSL2 | 5 | 2,23 | Yes |
| C2orf91 | 5 | 2,29 | Yes |
| THADA | 5 | 0,80 | No |
| SRBD1 | 5 | 1,19 | No |
| FSHR | 5 | 1,28 | No |
| EHBP1 | 5 | 0,88 | No |
| INO80B | 5 | 2,45 | Yes |
| WBP1 | 5 | 2,49 | Yes |
| MOGS | 5 | 2,48 | Yes |
| MRPL53 | 5 | 2,49 | Yes |
| CCDC142 | 5 | 2,39 | Yes |
| TTC31 | 5 | 2,41 | Yes |
| GLI2 | 5 | 1,11 | No |
| THSD7B | 5 | 0,45 | No |
| PSMD14 | 5 | 1,67 | No |
| FAP | 5 | 1,84 | No |
| KCNH7 | 5 | 0,75 | No |
| ZNF385B | 5 | 0,81 | No |
| IKZF2 | 5 | 1,42 | No |
| EPHA4 | 5 | 1,43 | No |
| UBE2E2 | 5 | 0,85 | No |
| ITGA9 | 5 | 0,88 | No |
| LRTM1 | 5 | 2,02 | Yes |
| COL8A1 | 5 | 1,40 | No |
| ZPLD1 | 5 | 0,86 | No |
| RABL3 | 5 | 1,96 | No |
| CPNE4 | 5 | 0,53 | No |
| TMEM108 | 5 | 0,90 | No |
| ZBTB38 | 5 | 1,55 | No |
| MBNL1 | 5 | 1,20 | No |
| KCNAB1 | 5 | 0,71 | No |
| C3orf55 | 5 | 1,50 | No |
| TNIK | 5 | 0,84 | No |
| NAALADL2 | 5 | 0,32 | No |
| WHSC1 | 5 | 1,62 | No |
| MAN2B2 | 5 | 2,04 | Yes |
| PCDH7 | 5 | 0,80 | No |
| SCFD2 | 5 | 0,72 | No |
| UGT2A3 | 5 | 2,27 | Yes |
| ADAMTS3 | 5 | 1,03 | No |
| C4orf22 | 5 | 0,61 | No |
| WDFY3 | 5 | 1,03 | No |
| ARHGAP24 | 5 | 0,69 | No |
| MAPK10 | 5 | 0,64 | No |
| FAM13A | 5 | 0,86 | No |
| ATOH1 | 5 | 2,50 | Yes |
| UNC5C | 5 | 0,86 | No |
| PPP3CA | 5 | 0,96 | No |
| BANK1 | 5 | 0,58 | No |
| ANK2 | 5 | 0,66 | No |
| KIAA1109 | 5 | 1,27 | No |
| LSM6 | 5 | 2,24 | Yes |
| RNF175 | 5 | 2,02 | Yes |
| CDH18 | 5 | 0,38 | No |
| PDZD2 | 5 | 0,75 | No |
| HCN1 | 5 | 0,79 | No |
| ISL1 | 5 | 2,37 | Yes |
| CWC27 | 5 | 1,11 | No |
| AP3B1 | 5 | 1,02 | No |
| ATG10 | 5 | 1,00 | No |
| ANKRD32 | 5 | 1,57 | No |
| CDKL3 | 5 | 1,38 | No |
| HDAC3 | 5 | 2,33 | Yes |
| DOCK2 | 5 | 0,78 | No |
| GMDS | 5 | 0,61 | No |
| FARS2 | 5 | 0,71 | No |
| RNF144B | 5 | 1,67 | No |
| TCP11 | 5 | 2,18 | Yes |
| ZNF76 | 5 | 2,15 | Yes |
| SUPT3H | 5 | 0,65 | No |
| KHDRBS2 | 5 | 0,62 | No |
| FAM46A | 5 | 1,09 | No |
| GRIK2 | 5 | 0,58 | No |
| SOBP | 5 | 1,37 | No |
| SCML4 | 5 | 1,58 | No |
| SLC35F1 | 5 | 0,82 | No |
| MAN1A1 | 5 | 1,35 | No |
| NKAIN2 | 5 | 0,41 | No |
| AKAP12 | 5 | 1,61 | No |
| CREB5 | 5 | 0,69 | No |
| BMPER | 5 | 1,11 | No |
| WBSCR17 | 5 | 0,64 | No |
| SEMA3A | 5 | 0,68 | No |
| SEMA3D | 5 | 1,29 | No |
| PDK4 | 5 | 2,36 | Yes |
| DLX5 | 5 | 2,45 | Yes |
| SLC26A5 | 5 | 1,72 | No |
| TFEC | 5 | 1,18 | No |
| CADPS2 | 5 | 0,65 | No |
| SND1 | 5 | 0,78 | No |
| BPGM | 5 | 2,16 | Yes |
| CALD1 | 5 | 1,18 | No |
| FAM180A | 5 | 2,28 | Yes |
| CSMD1 | 5 | 0,22 | No |
| TUSC3 | 5 | 0,91 | No |
| RBPMS | 5 | 1,30 | No |
| ST18 | 5 | 0,92 | No |
| CPA6 | 5 | 0,96 | No |
| EYA1 | 5 | 1,38 | No |
| RALYL | 5 | 0,53 | No |
| FBXO32 | 5 | 2,06 | Yes |
| GLIS3 | 5 | 0,69 | No |
| FOCAD | 5 | 0,94 | No |
| PGM5 | 5 | 1,25 | No |
| PTCH1 | 5 | 1,90 | No |
| PAPPA | 5 | 1,13 | No |
| STRBP | 5 | 1,40 | No |
| CELF2 | 5 | 0,95 | No |
| ITGA8 | 5 | 1,24 | No |
| FAM188A | 5 | 1,78 | No |
| MKX | 5 | 1,84 | No |
| BICC1 | 5 | 0,97 | No |
| SUPV3L1 | 5 | 2,21 | Yes |
| HKDC1 | 5 | 2,05 | Yes |
| SLIT1 | 5 | 1,30 | No |
| ARHGAP19-SLIT1 | 5 | 1,03 | No |
| SORCS3 | 5 | 0,61 | No |
| VTI1A | 5 | 0,88 | No |
| CTBP2 | 5 | 1,35 | No |
| CNGA4 | 5 | 2,40 | Yes |
| OR5G3 | 5 | 2,50 | Yes |
| FAM168A | 5 | 1,28 | No |
| SIAE | 5 | 1,92 | No |
| SPA17 | 5 | 2,24 | Yes |
| PKNOX2 | 5 | 1,08 | No |
| SLC2A13 | 5 | 0,91 | No |
| NCKAP5L | 5 | 2,14 | Yes |
| BCDIN3D | 5 | 2,45 | Yes |
| FAM19A2 | 5 | 0,65 | No |
| RP11-366L20.2 | 5 | 2,17 | Yes |
| TMTC2 | 5 | 0,77 | No |
| MGAT4C | 5 | 0,47 | No |
| POC1B | 5 | 1,66 | No |
| FRY | 5 | 1,10 | No |
| DCLK1 | 5 | 0,90 | No |
| CCDC122 | 5 | 1,98 | No |
| CLYBL | 5 | 1,03 | No |
| PSMB11 | 5 | 2,49 | Yes |
| CDH24 | 5 | 2,41 | Yes |
| ACIN1 | 5 | 2,15 | Yes |
| C14orf119 | 5 | 2,45 | Yes |
| C14orf164 | 5 | 1,74 | No |
| PRKD1 | 5 | 0,62 | No |
| BMP4 | 5 | 2,41 | Yes |
| MNAT1 | 5 | 1,15 | No |
| TRMT5 | 5 | 2,40 | Yes |
| SLC38A6 | 5 | 1,67 | No |
| SPTB | 5 | 1,53 | No |
| CHURC1-FNTB | 5 | 1,45 | No |
| FNTB | 5 | 1,44 | No |
| FUT8 | 5 | 0,94 | No |
| DPF3 | 5 | 1,06 | No |
| CEP128 | 5 | 0,74 | No |
| VRK1 | 5 | 1,50 | No |
| DKFZP547L112 | 5 | 2,50 | Yes |
| AGBL1 | 5 | 0,46 | No |
| SNX29 | 5 | 0,63 | No |
| KIAA0556 | 5 | 1,18 | No |
| CTD-2144E22.5 | 5 | 2,50 | Yes |
| ABCC11 | 5 | 1,81 | No |
| IRX5 | 5 | 2,47 | Yes |
| PRMT7 | 5 | 2,04 | Yes |
| SMPD3 | 5 | 1,74 | No |
| C16orf47 | 5 | 2,30 | Yes |
| RAP1GAP2 | 5 | 1,10 | No |
| SUPT6H | 5 | 2,13 | Yes |
| RPL23A | 5 | 2,45 | Yes |
| TLCD1 | 5 | 2,47 | Yes |
| NEK8 | 5 | 2,32 | Yes |
| TRAF4 | 5 | 2,43 | Yes |
| TMEM132E | 5 | 1,96 | No |
| ACACA | 5 | 0,97 | No |
| STARD3 | 5 | 2,24 | Yes |
| TCAP | 5 | 2,48 | Yes |
| PNMT | 5 | 2,48 | Yes |
| PGAP3 | 5 | 2,25 | Yes |
| HOXB1 | 5 | 2,48 | Yes |
| HOXB2 | 5 | 2,45 | Yes |
| HOXB3 | 5 | 1,96 | No |
| ZNF652 | 5 | 1,84 | No |
| RP11-81K2.1 | 5 | 1,63 | No |
| TAC4 | 5 | 2,39 | Yes |
| SMURF2 | 5 | 1,59 | No |
| CTD-2535L24.2 | 5 | 1,66 | No |
| CHST9 | 5 | 1,07 | No |
| MOCOS | 5 | 1,77 | No |
| C18orf25 | 5 | 1,71 | No |
| RNF165 | 5 | 1,49 | No |
| DYM | 5 | 0,81 | No |
| DAND5 | 5 | 2,39 | Yes |
| LYL1 | 5 | 2,47 | Yes |
| TRMT1 | 5 | 2,38 | Yes |
| STX10 | 5 | 2,43 | Yes |
| IER2 | 5 | 2,45 | Yes |
| CACNA1A | 5 | 0,82 | No |
| PLCB1 | 5 | 0,49 | No |
| NKX2-2 | 5 | 2,47 | Yes |
| MYH7B | 5 | 2,26 | Yes |
| TRPC4AP | 5 | 1,74 | No |
| CDH4 | 5 | 0,56 | No |
| TPTE | 5 | 1,56 | No |
| TMPRSS15 | 5 | 1,21 | No |
| APP | 5 | 1,02 | No |
| GRIK1 | 5 | 0,83 | No |
| TIAM1 | 5 | 0,79 | No |
| TTC28 | 5 | 0,56 | No |
| CNKSR2 | 5 | 1,05 | No |
| POLA1 | 5 | 1,00 | No |
| HDAC8 | 5 | 1,13 | No |
| IL1RAPL2 | 5 | 0,36 | No |
| COL4A5 | 5 | 1,10 | No |
| ZIC3 | 5 | 2,38 | Yes |
| AFF2 | 5 | 0,72 | No |

**table 9: Functional enrichment/over-representation analysis of MB NCCM genes (≥ 5, and ≥ 2 NCCMs per 100 *kbp*)**

| Source Database | ID | Pathway/Ontology | *p*-value (Adjusted) | Total no. of genes in Pathway/Ontology | No. of query genes overlapping | Genes |
|---|---|---|---|---|---|---|
| GO:Biological Process | GO:0007399 | nervous system development | 1,325×10^-26 | 2449 | 158 | DMD,AUTS2,IL1RAPL1,TENM2,SOX6,LSAMP,DCC, |
| GO:Biological Process | GO:0048699 | generation of neurons | 1,684×10^-22 | 1559 | 115 | DMD,AUTS2,IL1RAPL1,TENM2,DCC,NEGR1,ROBO |
| GO:Biological Process | GO:0030182 | neuron differentiation | 1,593×10^-21 | 1412 | 107 | DMD,AUTS2,IL1RAPL1,TENM2,DCC,NEGR1,ROBO |
| GO:Biological Process | GO:0022008 | neurogenesis | 1,797×10^-21 | 1675 | 118 | DMD,AUTS2,IL1RAPL1,TENM2,SOX6,DCC,NEGR1, |
| GO:Biological Process | GO:0009653 | anatomical structure morphogenesis | 8,062×10^-19 | 2797 | 155 | DMD,ARHGAP15,AUTS2,IL1RAPL1,SOX6,DCC,ROB |
| GO:Biological Process | GO:0048666 | neuron development | 2,162×10^-18 | 1149 | 90 | DMD,AUTS2,IL1RAPL1,TENM2,DCC,NEGR1,ROBO |
| GO:Biological Process | GO:0007275 | multicellular organism development | 2,536×10^-17 | 5613 | 240 | DMD,AUTS2,IL1RAPL1,MECOM,TENM2,SOX6,LSA |
| GO:Biological Process | GO:0048468 | cell development | 4,528×10^-17 | 2100 | 126 | DMD,AUTS2,IL1RAPL1,TENM2,DCC,NEGR1,ROBO |
| GO:Biological Process | GO:0048731 | system development | 1,094×10^-16 | 5052 | 222 | DMD,AUTS2,IL1RAPL1,MECOM,TENM2,SOX6,LSA |
| GO:Biological Process | GO:0032989 | cellular component morphogenesis | 2,132×10^-16 | 791 | 70 | DMD,AUTS2,IL1RAPL1,DCC,ROBO2,SLIT3,CNTNA |
| GO:Cellular Compone | GO:0030054 | cell junction | 5,039×10^-16 | 2105 | 120 | DMD,IL1RAPL1,CADM2,DLG2,PARD3B,TENM2,PT |
| GO:Cellular Compone | GO:0045202 | synapse | 6,042×10^-16 | 1349 | 91 | DMD,IL1RAPL1,CADM2,DLG2,TENM2,PTPRT,KCNI |
| GO:Biological Process | GO:0048856 | anatomical structure development | 8,101×10^-16 | 6106 | 250 | DMD,ARHGAP15,AUTS2,IL1RAPL1,MECOM,TENM |
| GO:Biological Process | GO:0007417 | central nervous system development | 1,148×10^-15 | 1032 | 80 | DMD,SOX6,DCC,ROBO2,GRID2,IMMP2L,CNTNAP2 |
| GO:Biological Process | GO:0032990 | cell part morphogenesis | 1,763×10^-15 | 699 | 64 | DMD,AUTS2,IL1RAPL1,DCC,ROBO2,SLIT3,CNTNA |
| GO:Biological Process | GO:0120039 | plasma membrane bounded cell projection morphogenesis | 7,057×10^-15 | 678 | 62 | DMD,AUTS2,IL1RAPL1,DCC,ROBO2,SLIT3,CNTNA |
| GO:Biological Process | GO:0048858 | cell projection morphogenesis | 9,435×10^-15 | 682 | 62 | DMD,AUTS2,IL1RAPL1,DCC,ROBO2,SLIT3,CNTNA |
| GO:Biological Process | GO:0048812 | neuron projection morphogenesis | 1,091×10^-14 | 664 | 61 | DMD,AUTS2,IL1RAPL1,DCC,ROBO2,SLIT3,CNTNA |
| GO:Biological Process | GO:0000902 | cell morphogenesis | 1,982×10^-14 | 1037 | 78 | DMD,ARHGAP15,AUTS2,IL1RAPL1,DCC,ROBO2,SL |
| GO:Biological Process | GO:0007420 | brain development | 7,974×10^-14 | 754 | 64 | DMD,ROBO2,GRID2,IMMP2L,CNTNAP2,MACROD |
| GO:Biological Process | GO:0048667 | cell morphogenesis involved in neuron differentiation | 1,690×10^-13 | 602 | 56 | AUTS2,IL1RAPL1,DCC,ROBO2,SLIT3,NRXN3,DAB1, |
| GO:Biological Process | GO:0060322 | head development | 3,412×10^-13 | 798 | 65 | DMD,ROBO2,GRID2,IMMP2L,CNTNAP2,MACROD |
| GO:Biological Process | GO:0032502 | developmental process | 3,476×10^-13 | 6628 | 257 | DMD,ARHGAP15,AUTS2,IL1RAPL1,MECOM,TENM |
| GO:Cellular Compone | GO:0043005 | neuron projection | 7,271×10^-13 | 1380 | 86 | DMD,AUTS2,IL1RAPL1,CADM2,DLG2,TENM2,KCN |
| GO:Biological Process | GO:0007411 | axon guidance | 1,067×10^-12 | 283 | 37 | DCC,ROBO2,SLIT3,NRXN3,DAB1,ROBO1,EFNA5,SE |
| GO:Biological Process | GO:0097485 | neuron projection guidance | 1,198×10^-12 | 284 | 37 | DCC,ROBO2,SLIT3,NRXN3,DAB1,ROBO1,EFNA5,SE |
| GO:Biological Process | GO:0032501 | multicellular organismal process | 1,547×10^-12 | 7946 | 290 | DMD,AUTS2,IL1RAPL1,MECOM,TENM2,SOX6,LSA |
| GO:Cellular Compone | GO:0097060 | synaptic membrane | 7,859×10^-12 | 384 | 40 | DMD,IL1RAPL1,DLG2,TENM2,PTPRT,KCND2,DCC, |
| GO:Biological Process | GO:0031175 | neuron projection development | 8,265×10^-12 | 1012 | 72 | DMD,AUTS2,IL1RAPL1,DCC,NEGR1,ROBO2,GRID2 |
| GO:Biological Process | GO:0048513 | animal organ development | 1,380×10^-11 | 3672 | 166 | DMD,MECOM,SOX6,DCC,ROBO2,GRID2,SLIT3,HD |
| GO:Biological Process | GO:0048869 | cellular developmental process | 1,698×10^-11 | 4428 | 189 | DMD,AUTS2,IL1RAPL1,MECOM,TENM2,SOX6,DC |
| GO:Biological Process | GO:0030154 | cell differentiation | 2,806×10^-11 | 4350 | 186 | DMD,AUTS2,IL1RAPL1,MECOM,TENM2,SOX6,DC |
| GO:Cellular Compone | GO:0098978 | glutamatergic synapse | 4,460×10^-11 | 349 | 37 | IL1RAPL1,PTPRT,KCND2,GRID2,NLGN1,PTPRD,SYT |
| GO:Biological Process | GO:0007409 | axonogenesis | 1,568×10^-10 | 478 | 45 | AUTS2,DCC,ROBO2,SLIT3,NRXN3,DAB1,ROBO1,E |
| GO:Biological Process | GO:0034330 | cell junction organization | 4,321×10^-10 | 722 | 56 | IL1RAPL1,CADM2,DLG2,PTPRT,ROBO2,GRID2,CN |
| GO:Biological Process | GO:0000904 | cell morphogenesis involved in differentiation | 5,694×10^-10 | 749 | 57 | AUTS2,IL1RAPL1,DCC,ROBO2,SLIT3,NRXN3,DAB1, |
| GO:Cellular Compone | GO:0030424 | axon | 1,043×10^-09 | 660 | 50 | DMD,AUTS2,IL1RAPL1,CADM2,DLG2,TENM2,DCC, |
| GO:Biological Process | GO:0050808 | synapse organization | 1,248×10^-09 | 428 | 41 | IL1RAPL1,DLG2,PTPRT,ROBO2,GRID2,CNTN5,NLG |
| GO:Biological Process | GO:0061564 | axon development | 3,778×10^-09 | 523 | 45 | AUTS2,DCC,ROBO2,SLIT3,NRXN3,DAB1,ROBO1,E |
| GO:Cellular Compone | GO:0098794 | postsynapse | 3,952×10^-09 | 639 | 48 | DMD,IL1RAPL1,DLG2,TENM2,PTPRT,KCND2,DCC, |
| GO:Cellular Compone | GO:0098984 | neuron to neuron synapse | 4,081×10^-09 | 366 | 35 | DMD,DLG2,PTPRT,KCND2,DCC,GRID2,DAB1,NLGN |
| GO:Biological Process | GO:0009887 | animal organ morphogenesis | 4,357×10^-09 | 1073 | 69 | SOX6,ROBO2,SLIT3,SLC4A10,TENM3,GPC6,ROBO |
| GO:Cellular Compone | GO:0042995 | cell projection | 6,774×10^-09 | 2330 | 110 | DMD,AUTS2,IL1RAPL1,CADM2,DLG2,TENM2,KCN |
| GO:Cellular Compone | GO:0045211 | postsynaptic membrane | 7,496×10^-09 | 280 | 30 | DMD,IL1RAPL1,DLG2,TENM2,PTPRT,KCND2,DCC, |
| GO:Cellular Compone | GO:0036477 | somatodendritic compartment | 1,050×10^-08 | 871 | 57 | DMD,IL1RAPL1,DLG2,TENM2,KCND2,GRID2,CNTN |
| GO:Cellular Compone | GO:0032279 | asymmetric synapse | 1,079×10^-08 | 340 | 33 | DMD,DLG2,PTPRT,KCND2,DCC,GRID2,DAB1,NLGN |
| GO:Cellular Compone | GO:0120025 | plasma membrane bounded cell projection | 1,149×10^-08 | 2229 | 106 | DMD,AUTS2,IL1RAPL1,CADM2,DLG2,TENM2,KCN |
| GO:Biological Process | GO:0098742 | cell-cell adhesion via plasma-membrane adhesion molecules | 1,875×10^-08 | 277 | 31 | IL1RAPL1,TENM2,PTPRT,ROBO2,GRID2,DAB1,NLG |
| GO:Cellular Compone | GO:0014069 | postsynaptic density | 3,029×10^-08 | 334 | 32 | DMD,DLG2,PTPRT,KCND2,DCC,GRID2,DAB1,NLGN |
| GO:Cellular Compone | GO:0099572 | postsynaptic specialization | 4,246×10^-08 | 358 | 33 | DMD,DLG2,PTPRT,KCND2,DCC,GRID2,DAB1,NLGN |
| GO:Biological Process | GO:0030900 | forebrain development | 7,160×10^-08 | 385 | 36 | DMD,ROBO2,CNTNAP2,DAB1,SLC4A10,PRKG1,RO |
| GO:Biological Process | GO:0009888 | tissue development | 7,991×10^-08 | 2107 | 104 | DMD,SOX6,EYS,ROBO2,HDAC9,RBFOX1,TENM4,G |
| GO:Biological Process | GO:0007155 | cell adhesion | 9,518×10^-08 | 1492 | 82 | DMD,IL1RAPL1,CADM2,DLG2,PARD3B,TENM2,PT |
| GO:Biological Process | GO:0022610 | biological adhesion | 1,203×10^-07 | 1499 | 82 | DMD,IL1RAPL1,CADM2,DLG2,PARD3B,TENM2,PT |
| GO:Biological Process | GO:0006928 | movement of cell or subcellular component | 3,647×10^-07 | 2311 | 109 | DMD,AUTS2,DLG2,PTPRT,DCC,ROBO2,SLIT3,HDA |
| GO:Biological Process | GO:0050804 | modulation of chemical synaptic transmission | 4,153×10^-07 | 430 | 37 | DCC,GRID2,SLC4A10,NLGN1,PTPRD,SYT1,MEF2C, |
| GO:Biological Process | GO:0099177 | regulation of trans-synaptic signaling | 4,437×10^-07 | 431 | 37 | DCC,GRID2,SLC4A10,NLGN1,PTPRD,SYT1,MEF2C, |
| GO:Biological Process | GO:0050919 | negative chemotaxis | 4,565×10^-07 | 48 | 13 | ROBO2,SLIT3,ROBO1,EFNA5,NRG3,SEMA6D,NRG |
| GO:Biological Process | GO:0120036 | plasma membrane bounded cell projection organization | 4,764×10^-07 | 1569 | 83 | DMD,AUTS2,IL1RAPL1,TENM2,DCC,NEGR1,ROBO |
| GO:Cellular Compone | GO:0034703 | cation channel complex | 1,014×10^-06 | 227 | 24 | DLG2,KCND2,DPP10,CNTNAP2,CACNA2D3,NLGN |
| GO:Biological Process | GO:0099537 | trans-synaptic signaling | 1,140×10^-06 | 713 | 49 | IL1RAPL1,DLG2,KCND2,DCC,GRID2,SLC4A10,NLG |
| GO:Biological Process | GO:0099536 | synaptic signaling | 1,216×10^-06 | 738 | 50 | IL1RAPL1,DLG2,KCND2,DCC,GRID2,SLC4A10,NLG |
| GO:Cellular Compone | GO:0098590 | plasma membrane region | 1,604×10^-06 | 1240 | 66 | DMD,IL1RAPL1,DLG2,PARD3B,TENM2,PTPRT,KCN |
| GO:Biological Process | GO:0007267 | cell-cell signaling | 1,612×10^-06 | 1722 | 87 | IL1RAPL1,DLG2,KCND2,DCC,GRID2,SLC4A10,NLG |
| GO:Biological Process | GO:0030030 | cell projection organization | 1,627×10^-06 | 1609 | 83 | DMD,AUTS2,IL1RAPL1,TENM2,DCC,NEGR1,ROBO |
| GO:Biological Process | GO:0021537 | telencephalon development | 1,698×10^-06 | 256 | 27 | DMD,ROBO2,CNTNAP2,DAB1,ROBO1,NRG3,KIRR |
| GO:Biological Process | GO:0035107 | appendage morphogenesis | 1,722×10^-06 | 141 | 20 | CACNA1C,RARB,RUNX2,PCSK5,FMN1,PBX1,TGFB |
| GO:Biological Process | GO:0035108 | limb morphogenesis | 1,722×10^-06 | 141 | 20 | CACNA1C,RARB,RUNX2,PCSK5,FMN1,PBX1,TGFB |
| GO:Biological Process | GO:0006935 | chemotaxis | 1,851×10^-06 | 653 | 46 | DCC,ROBO2,SLIT3,NRXN3,DAB1,ROBO1,EFNA5,D |
| GO:Biological Process | GO:0007610 | behavior | 2,004×10^-06 | 586 | 43 | KCND2,NEGR1,CNTNAP2,NRXN3,DAB1,SLC4A10, |
| GO:Biological Process | GO:0042330 | taxis | 2,145×10^-06 | 656 | 46 | DCC,ROBO2,SLIT3,NRXN3,DAB1,ROBO1,EFNA5,D |
| GO:Biological Process | GO:0098916 | anterograde trans-synaptic signaling | 2,380×10^-06 | 705 | 48 | DLG2,KCND2,DCC,GRID2,SLC4A10,NLGN1,PTPRD, |
| GO:Biological Process | GO:0007268 | chemical synaptic transmission | 2,380×10^-06 | 705 | 48 | DLG2,KCND2,DCC,GRID2,SLC4A10,NLGN1,PTPRD, |
| GO:Biological Process | GO:0040011 | locomotion | 2,491×10^-06 | 2028 | 97 | AUTS2,PTPRT,DCC,ROBO2,SLIT3,HDAC9,NRXN3,D |
| GO:Biological Process | GO:0016358 | dendrite development | 2,607×10^-06 | 243 | 26 | IL1RAPL1,DCC,DAB1,NLGN1,PTPRD,PRKG1,MEF2 |
| GO:Cellular Compone | GO:0099634 | postsynaptic specialization membrane | 2,659×10^-06 | 120 | 17 | DLG2,PTPRT,KCND2,DCC,GRID2,NLGN1,LRRC4C,C |
| GO:Cellular Compone | GO:0030425 | dendrite | 4,169×10^-06 | 638 | 42 | IL1RAPL1,TENM2,KCND2,GRID2,CNTNAP2,SLC4A |
| GO:Biological Process | GO:0007507 | heart development | 4,293×10^-06 | 578 | 42 | SOX6,ROBO2,SLIT3,HDAC9,TENM4,ROBO1,MEF2 |
| GO:Cellular Compone | GO:0097447 | dendritic tree | 4,561×10^-06 | 640 | 42 | IL1RAPL1,TENM2,KCND2,GRID2,CNTNAP2,SLC4A |
| GO:Biological Process | GO:0098609 | cell-cell adhesion | 5,219×10^-06 | 894 | 55 | IL1RAPL1,DLG2,TENM2,PTPRT,NEGR1,ROBO2,GR |
| GO:Biological Process | GO:0050793 | regulation of developmental process | 5,377×10^-06 | 2605 | 115 | DMD,ARHGAP15,IL1RAPL1,SOX6,DCC,ROBO2,GR |
| GO:Molecular Functi | GO:0000978 | RNA polymerase II cis-regulatory region sequence-specific DNA bindi | 5,977×10^-06 | 64 | 1190 | MECOM,SOX6,ESRRG,ZBTB20,EBF1,ZNF536,FOXP |
| GO:Biological Process | GO:0035249 | synaptic transmission, glutamatergic | 6,081×10^-06 | 94 | 16 | GRID2,NLGN1,SYT1,MEF2C,NRXN1,GRM8,GRIK3, |
| GO:Biological Process | GO:0007157 | heterophilic cell-cell adhesion via plasma membrane cell adhesion m | 8,338×10^-06 | 49 | 12 | IL1RAPL1,TENM2,GRID2,NLGN1,TENM3,PTPRD,TE |
| GO:Molecular Functi | GO:0000987 | cis-regulatory region sequence-specific DNA binding | 1,091×10^-05 | 64 | 1209 | MECOM,SOX6,ESRRG,ZBTB20,EBF1,ZNF536,FOXP |
| GO:Biological Process | GO:0021545 | cranial nerve development | 1,369×10^-05 | 51 | 12 | DMD,NAV2,EPHB1,GLI3,EXT1,KCNA2,ISL1,SEMA3 |
| GO:Cellular Compone | GO:0098793 | presynapse | 1,432×10^-05 | 521 | 36 | DMD,CNTN5,NRXN3,SLC4A10,NLGN1,PTPRD,DEN |
| GO:Biological Process | GO:0048736 | appendage development | 1,587×10^-05 | 176 | 21 | CACNA1C,RARB,RUNX2,PCSK5,FMN1,PBX1,TGFB |
| GO:Biological Process | GO:0060173 | limb development | 1,587×10^-05 | 176 | 21 | CACNA1C,RARB,RUNX2,PCSK5,FMN1,PBX1,TGFB |
| GO:Cellular Compone | GO:0034702 | ion channel complex | 1,639×10^-05 | 302 | 26 | DLG2,KCND2,DPP10,GRID2,CNTNAP2,CACNA2D3, |
| GO:Biological Process | GO:0050807 | regulation of synapse organization | 1,895×10^-05 | 212 | 23 | IL1RAPL1,PTPRT,ROBO2,GRID2,NLGN1,PTPRD,GP |
| GO:Biological Process | GO:0051239 | regulation of multicellular organismal process | 2,325×10^-05 | 2861 | 121 | DMD,IL1RAPL1,SOX6,KCND2,DCC,ROBO2,GRID2, |
| GO:Biological Process | GO:0045595 | regulation of cell differentiation | 2,689×10^-05 | 1679 | 82 | DMD,IL1RAPL1,SOX6,DCC,ROBO2,HDAC9,RBFOX1 |
| GO:Biological Process | GO:0022603 | regulation of anatomical structure morphogenesis | 2,787×10^-05 | 1042 | 59 | ARHGAP15,IL1RAPL1,DCC,ROBO2,DAB1,NLGN1,P |
| GO:Cellular Compone | GO:0098839 | postsynaptic density membrane | 2,932×10^-05 | 94 | 14 | DLG2,PTPRT,DCC,GRID2,LRRC4C,CACNA1C,NRG1, |
| GO:Biological Process | GO:0045664 | regulation of neuron differentiation | 3,775×10^-05 | 202 | 22 | DMD,DAB1,NLGN1,ZNF536,MEF2C,ZFHX3,CNTN4 |
| GO:Cellular Compone | GO:0043025 | neuronal cell body | 3,830×10^-05 | 518 | 35 | DMD,DLG2,KCND2,CNTNAP2,DAB1,SLC4A10,DEN |
| REACTOME | R-HSA-1266738 | Developmental Biology Homo sapiens | 4,043×10^-05 | 786 | 48 | ROBO2;PSMB11;HDAC3;PSMD14;SEMA3A;CACN |
| GO:Biological Process | GO:0007389 | pattern specification process | 4,155×10^-05 | 442 | 34 | ROBO2,ROBO1,MEF2C,NRG3,PBX3,ERBB4,PCSK5, |
| GO:Biological Process | GO:0048589 | developmental growth | 4,268×10^-05 | 649 | 43 | DMD,AUTS2,DCC,EYS,SLIT3,SLC4A10,TENM4,SYT1 |
| GO:Biological Process | GO:0051960 | regulation of nervous system development | 4,394×10^-05 | 443 | 34 | IL1RAPL1,DCC,ROBO2,GRID2,DAB1,NLGN1,PTPR |
| GO:Biological Process | GO:0050803 | regulation of synapse structure or activity | 4,975×10^-05 | 223 | 23 | IL1RAPL1,PTPRT,ROBO2,GRID2,NLGN1,PTPRD,GP |
| GO:Biological Process | GO:0072359 | circulatory system development | 5,036×10^-05 | 1194 | 64 | SOX6,ROBO2,SLIT3,HDAC9,IMMP2L,NRXN3,TEN |
| GO:Biological Process | GO:0021953 | central nervous system neuron differentiation | 5,178×10^-05 | 188 | 21 | DCC,ROBO2,GRID2,SLC4A10,ROBO1,AGBL4,NRX |
| GO:Biological Process | GO:0031344 | regulation of cell projection organization | 5,826×10^-05 | 656 | 43 | DMD,AUTS2,IL1RAPL1,TENM2,DCC,NEGR1,ROBO |
| GO:Biological Process | GO:0001764 | neuron migration | 6,752×10^-05 | 157 | 19 | AUTS2,DCC,DAB1,PRKG1,MEF2C,NRG3,KIRREL3, |
| GO:Cellular Compone | GO:1902495 | transmembrane transporter complex | 7,032×10^-05 | 325 | 26 | DLG2,KCND2,DPP10,GRID2,CNTNAP2,CACNA2D3, |
| GO:Biological Process | GO:0120035 | regulation of plasma membrane bounded cell projection organizatior | 7,614×10^-05 | 638 | 42 | DMD,AUTS2,IL1RAPL1,TENM2,DCC,NEGR1,ROBO |
| GO:Biological Process | GO:0048729 | tissue morphogenesis | 9,400×10^-05 | 667 | 43 | ROBO2,GPC6,FOXP1,ROBO1,MEF2C,NRG1,ASTN2 |
| GO:Cellular Compone | GO:0044297 | cell body | 1,159×10^-04 | 592 | 37 | DMD,DLG2,KCND2,CNTNAP2,DAB1,SLC4A10,DEN |
| GO:Biological Process | GO:0007416 | synapse assembly | 1,241×10^-04 | 180 | 20 | IL1RAPL1,ROBO2,GRID2,CNTN5,NLGN1,PTPRD,G |
| GO:Cellular Compone | GO:0031226 | intrinsic component of plasma membrane | 1,360×10^-04 | 1726 | 77 | DLG2,TENM2,PTPRT,KCND2,DCC,DPP10,GRID2,C |
| GO:Molecular Functi | GO:0008066 | glutamate receptor activity | 1,409×10^-04 | 8 | 27 | GRID2,GRM8,GRIK3,GRIA4,GRM7,GRIN2B,GRIK2, |
| GO:Biological Process | GO:0060560 | developmental growth involved in morphogenesis | 1,438×10^-04 | 236 | 23 | AUTS2,DCC,SLIT3,SYT1,SEMA6D,ESR1,PARK2,FM |
| GO:Cellular Compone | GO:1990351 | transporter complex | 1,682×10^-04 | 340 | 26 | DLG2,KCND2,DPP10,GRID2,CNTNAP2,CACNA2D3, |
| GO:Biological Process | GO:0048863 | stem cell differentiation | 1,793×10^-04 | 258 | 24 | SOX6,MSI2,MEF2C,FAM172A,SEMA6D,NRG1,ERB |
| GO:Cellular Compone | GO:0043204 | perikaryon | 1,919×10^-04 | 160 | 17 | DLG2,KCND2,CNTNAP2,SLC4A10,CACNA1C,ASTN |
| GO:Biological Process | GO:0035113 | embryonic appendage morphogenesis | 1,977×10^-04 | 119 | 16 | CACNA1C,RARB,RUNX2,PBX1,TGFB2,FRAS1,HAND |
| GO:Biological Process | GO:0030326 | embryonic limb morphogenesis | 1,977×10^-04 | 119 | 16 | CACNA1C,RARB,RUNX2,PBX1,TGFB2,FRAS1,HAND |
| GO:Biological Process | GO:0021675 | nerve development | 2,205×10^-04 | 77 | 13 | DMD,NAV2,EPHB1,GLI3,EXT1,KCNA2,ISL1,SEMA3 |
| GO:Cellular Compone | GO:0099240 | intrinsic component of synaptic membrane | 2,289×10^-04 | 162 | 17 | PTPRT,KCND2,DCC,GRID2,CNTN5,NLGN1,PTPRD,L |
| GO:Biological Process | GO:0060914 | heart formation | 2,989×10^-04 | 33 | 9 | ROBO2,ROBO1,MEF2C,HAND2,EXT1,AXIN2,ISL1,E |
| GO:Biological Process | GO:0007215 | glutamate receptor signaling pathway | 3,009×10^-04 | 54 | 11 | GRID2,GRM8,GRIK3,GRIA4,GRM7,GRIN2B,GRIK2, |
| GO:Biological Process | GO:0007158 | neuron cell-cell adhesion | 3,259×10^-04 | 17 | 7 | NRXN3,NLGN1,NRXN1,ASTN2,CNTN4,TNR,ASTN1 |
| GO:Cellular Compone | GO:0071944 | cell periphery | 3,285×10^-04 | 6169 | 204 | DMD,IL1RAPL1,CADM2,DLG2,PARD3B,TENM2,EN |
| KEGG | KEGG:04360 | Axon guidance | 3,350×10^-04 | 181 | 18 | EPHA5;ROBO2;EPHA4;TRPC6;DCC;SEMA3D;SEMA |
| GO:Cellular Compone | GO:0005911 | cell-cell junction | 3,407×10^-04 | 494 | 32 | PARD3B,CNTNAP2,CDH12,CTNNA3,KIRREL3,SKAP |
| GO:Cellular Compone | GO:0034705 | potassium channel complex | 4,212×10^-04 | 100 | 13 | DLG2,KCND2,DPP10,CNTNAP2,DPP6,GRIK3,KCNI |
| GO:Biological Process | GO:0034765 | regulation of ion transmembrane transport | 4,671×10^-04 | 583 | 38 | DMD,DLG2,KCND2,DPP10,CACNA2D3,NLGN1,ME |
| GO:Biological Process | GO:0035235 | ionotropic glutamate receptor signaling pathway | 5,517×10^-04 | 26 | 8 | GRID2,GRIK3,GRIA4,GRIN2B,GRIK2,APP,GRIK1,TI |
| GO:Biological Process | GO:1905114 | cell surface receptor signaling pathway involved in cell-cell signaling | 5,595×10^-04 | 636 | 40 | GRID2,NLGN1,GPC6,MEF2C,NRXN1,MAP3K7,ESR |
| GO:Biological Process | GO:0034762 | regulation of transmembrane transport | 5,800×10^-04 | 588 | 38 | DMD,DLG2,KCND2,DPP10,CACNA2D3,NLGN1,ME |
| GO:Biological Process | GO:0048813 | dendrite morphogenesis | 5,913×10^-04 | 145 | 17 | IL1RAPL1,NLGN1,PTPRD,CTNNA2,ZNF365,DNM3, |
| GO:Biological Process | GO:0045944 | positive regulation of transcription by RNA polymerase II | 6,748×10^-04 | 1197 | 61 | AUTS2,MECOM,ESRRG,MEF2C,ZNF462,SKAP1,PB |
| GO:Biological Process | GO:0051963 | regulation of synapse assembly | 6,934×10^-04 | 99 | 14 | IL1RAPL1,ROBO2,GRID2,NLGN1,PTPRD,GPC6,ME |
| GO:Biological Process | GO:0010975 | regulation of neuron projection development | 7,128×10^-04 | 451 | 32 | DMD,IL1RAPL1,DCC,NEGR1,ROBO2,GRID2,DAB1, |
| GO:Biological Process | GO:0008038 | neuron recognition | 7,472×10^-04 | 47 | 10 | ROBO2,CNTNAP2,OPCML,ROBO1,CNTN4,EXT1,N |
| GO:Biological Process | GO:0048598 | embryonic morphogenesis | 7,815×10^-04 | 595 | 38 | TENM4,MEF2C,CACNA1C,RARB,RUNX2,EXOC4,PA |
| GO:Biological Process | GO:0003128 | heart field specification | 8,074×10^-04 | 19 | 7 | ROBO2,ROBO1,MEF2C,EXT1,AXIN2,ISL1,BMP4 |
| GO:Biological Process | GO:0051966 | regulation of synaptic transmission, glutamatergic | 8,123×10^-04 | 72 | 12 | NLGN1,SYT1,MEF2C,NRXN1,GRM8,GRIK3,TNR,GR |
| GO:Biological Process | GO:1901888 | regulation of cell junction assembly | 8,365×10^-04 | 202 | 20 | IL1RAPL1,ROBO2,GRID2,NLGN1,PTPRD,GPC6,LIM |
| GO:Biological Process | GO:2000310 | regulation of NMDA receptor activity | 8,800×10^-04 | 37 | 9 | DLG2,NLGN1,MEF2C,NRXN1,PPARGC1A,GRIA4,RE |
| GO:Biological Process | GO:0060284 | regulation of cell development | 8,877×10^-04 | 502 | 34 | IL1RAPL1,DCC,ROBO2,HDAC9,DAB1,PTPRD,TENM |
| GO:Biological Process | GO:0060429 | epithelium development | 1,018×10^-03 | 1325 | 65 | DMD,ROBO2,GPC6,FOXP1,ROBO1,MEF2C,NRG1, |
| GO:Biological Process | GO:0008015 | blood circulation | 1,032×10^-03 | 553 | 36 | DMD,KCND2,IMMP2L,CACNA2D3,PRKG1,RYR3,SG |
| KEGG | KEGG:04724 | Glutamatergic synapse | 1,099×10^-03 | 114 | 13 | PRKCB;GRIK3;CACNA1A;GRIK1;CACNA1C;GRIK2; |
| GO:Biological Process | GO:0099173 | postsynapse organization | 1,117×10^-03 | 169 | 18 | IL1RAPL1,DLG2,GRID2,NLGN1,PTPRD,NRXN1,ZNF |
| GO:Molecular Functi | GO:0000977 | RNA polymerase II transcription regulatory region sequence-specific | 1,163×10^-03 | 65 | 1408 | MECOM,SOX6,NPAS3,ESRRG,ZBTB20,EBF1,ZNF53 |
| GO:Biological Process | GO:0034329 | cell junction assembly | 1,165×10^-03 | 438 | 31 | IL1RAPL1,ROBO2,GRID2,CNTN5,NLGN1,PTPRD,G |
| GO:Cellular Compone | GO:0005886 | plasma membrane | 1,259×10^-03 | 5681 | 188 | DMD,IL1RAPL1,CADM2,DLG2,PARD3B,TENM2,EN |
| GO:Biological Process | GO:0021510 | spinal cord development | 1,290×10^-03 | 104 | 14 | SOX6,DCC,ROBO2,DAB1,PBX3,MDGA2,PAX6,GLI3, |
| KEGG | KEGG:05412 | Arrhythmogenic right ventricular cardiomyopathy (ARVC) | 1,391×10^-03 | 72 | 10 | TCF7L2;SGCD;CACNA2D3;ITGA8;DMD;CTNNA3;C |
| GO:Biological Process | GO:0021602 | cranial nerve morphogenesis | 1,416×10^-03 | 29 | 8 | DMD,EPHB1,GLI3,KCNA2,SEMA3A,HOXB1,HOXB2 |
| GO:Biological Process | GO:0010646 | regulation of cell communication | 1,436×10^-03 | 3543 | 135 | DMD,ARHGAP15,AUTS2,DLG2,MECOM,PTPRT,DC |
| GO:Biological Process | GO:0014033 | neural crest cell differentiation | 1,455×10^-03 | 90 | 13 | MEF2C,FAM172A,SEMA6D,NRG1,ERBB4,HAND2, |
| GO:Molecular Functi | GO:0001067 | regulatory region nucleic acid binding | 1,516×10^-03 | 68 | 1509 | MECOM,SOX6,NPAS3,ESRRG,ZBTB20,EBF1,ZNF53 |
| GO:Biological Process | GO:0003013 | circulatory system process | 1,582×10^-03 | 637 | 39 | DMD,KCND2,IMMP2L,CACNA2D3,PRKG1,RYR3,SG |
| GO:Cellular Compone | GO:0099699 | integral component of synaptic membrane | 1,591×10^-03 | 148 | 15 | PTPRT,KCND2,DCC,GRID2,NLGN1,PTPRD,LRRC4C, |
| GO:Biological Process | GO:0060537 | muscle tissue development | 1,613×10^-03 | 399 | 29 | DMD,SOX6,HDAC9,RBFOX1,TENM4,MEF2C,SGCD, |
| GO:Biological Process | GO:0060485 | mesenchyme development | 1,673×10^-03 | 291 | 24 | ROBO2,ROBO1,MEF2C,FAM172A,SEMA6D,NRG1, |
| GO:Cellular Compone | GO:0044304 | main axon | 1,762×10^-03 | 66 | 10 | DLG2,ROBO2,CNTNAP2,SPTBN1,IQCJ-SCHIP1,ANK |
| GO:Biological Process | GO:0007423 | sensory organ development | 1,875×10^-03 | 567 | 36 | TENM3,RCN1,PCDH15,CACNA1C,PBX3,THRB,RAR |
| GO:Biological Process | GO:0001822 | kidney development | 1,894×10^-03 | 293 | 24 | ROBO2,MEF2C,KIRREL3,RARB,PCSK5,FMN1,PBX1, |
| GO:Biological Process | GO:0021554 | optic nerve development | 2,079×10^-03 | 14 | 6 | NAV2,EPHB1,GLI3,EXT1,KCNA2,NKX2-2 |
| GO:Biological Process | GO:1903508 | positive regulation of nucleic acid-templated transcription | 2,105×10^-03 | 1617 | 74 | AUTS2,MECOM,NPAS3,ESRRG,MEF2C,ZNF462,SK |
| GO:Biological Process | GO:0045893 | positive regulation of transcription, DNA-templated | 2,105×10^-03 | 1617 | 74 | AUTS2,MECOM,NPAS3,ESRRG,MEF2C,ZNF462,SK |
| GO:Biological Process | GO:1902680 | positive regulation of RNA biosynthetic process | 2,155×10^-03 | 1618 | 74 | AUTS2,MECOM,NPAS3,ESRRG,MEF2C,ZNF462,SK |
| GO:Biological Process | GO:0045597 | positive regulation of cell differentiation | 2,158×10^-03 | 879 | 48 | DMD,IL1RAPL1,SOX6,ROBO2,DAB1,PTPRD,TENM |
| GO:Biological Process | GO:0003002 | regionalization | 2,165×10^-03 | 338 | 26 | ROBO2,ROBO1,MEF2C,PBX3,PCSK5,PAX6,PBX1,G |
| REACTOME | R-HSA-422475 | Axon guidance Homo sapiens | 2,350×10^-03 | 515 | 32 | ROBO2;PSMB11;PSMD14;SEMA3A;CACNA1C;EF |
| REACTOME | R-HSA-418990 | Adherens junctions interactions Homo sapiens | 2,350×10^-03 | 31 | 7 | CDH4;CADM1;CADM2;CDH12;CDH24;CDH8;CDH |
| REACTOME | R-HSA-2022928 | HS-GAG biosynthesis Homo sapiens | 2,350×10^-03 | 31 | 7 | EXT1;GPC3;GPC5;HS3ST5;GPC6;HS2ST1;HS6ST3 |
| GO:Biological Process | GO:0023051 | regulation of signaling | 2,525×10^-03 | 3578 | 135 | DMD,ARHGAP15,AUTS2,DLG2,MECOM,PTPRT,DC |
| GO:Biological Process | GO:0048645 | animal organ formation | 2,557×10^-03 | 66 | 11 | ROBO2,ROBO1,MEF2C,HAND2,GLI3,EXT1,AXIN2, |
| GO:Biological Process | GO:0048568 | embryonic organ development | 2,675×10^-03 | 432 | 30 | MEF2C,PBX3,RARB,RUNX2,PCSK5,PAX6,NTN1,PB |
| GO:Biological Process | GO:0065008 | regulation of biological quality | 2,807×10^-03 | 4131 | 151 | DMD,ARHGAP15,IL1RAPL1,PTPRT,KCND2,DCC,RO |
| GO:Molecular Functi | GO:0000976 | transcription regulatory region sequence-specific DNA binding | 2,858×10^-03 | 67 | 1507 | MECOM,SOX6,NPAS3,ESRRG,ZBTB20,EBF1,ZNF53 |
| GO:Biological Process | GO:0050767 | regulation of neurogenesis | 2,994×10^-03 | 366 | 27 | IL1RAPL1,DCC,ROBO2,DAB1,PTPRD,TENM4,ROB |
| GO:Molecular Functi | GO:1990837 | sequence-specific double-stranded DNA binding | 2,999×10^-03 | 69 | 1570 | MECOM,SOX6,NPAS3,ESRRG,ZBTB20,EBF1,ZNF53 |
| GO:Molecular Functi | GO:0003690 | double-stranded DNA binding | 3,134×10^-03 | 72 | 1664 | MECOM,SOX6,NPAS3,ESRRG,ZBTB20,EBF1,ZNF53 |
| GO:Molecular Functi | GO:0004970 | ionotropic glutamate receptor activity | 3,159×10^-03 | 6 | 19 | GRID2,GRIK3,GRIA4,GRIN2B,GRIK2,GRIK1 |
| GO:Biological Process | GO:0072001 | renal system development | 3,256×10^-03 | 302 | 24 | ROBO2,MEF2C,KIRREL3,RARB,PCSK5,FMN1,PBX1, |
| GO:Biological Process | GO:0097120 | receptor localization to synapse | 3,511×10^-03 | 55 | 10 | DLG2,NLGN1,ANKS1B,GPC6,NRXN1,GRIP1,GPHN, |
| GO:Biological Process | GO:0090596 | sensory organ morphogenesis | 3,672×10^-03 | 262 | 22 | TENM3,THRB,RARB,SDK1,PAX6,NTN1,PTPRM,EP |
| GO:Biological Process | GO:0035295 | tube development | 3,799×10^-03 | 1146 | 57 | ROBO2,HDAC9,NRXN3,DAB1,FOXP1,ROBO1,MEF |
| GO:Biological Process | GO:0002009 | morphogenesis of an epithelium | 3,866×10^-03 | 560 | 35 | GPC6,FOXP1,MEF2C,ASTN2,ESR1,FMN1,NTN1,PB |
| GO:Biological Process | GO:0003008 | system process | 3,933×10^-03 | 2293 | 95 | DMD,KCND2,EYS,GRID2,IMMP2L,CNTNAP2,CNTN |
| GO:Biological Process | GO:0001709 | cell fate determination | 4,274×10^-03 | 44 | 9 | MEF2C,EBF2,PAX6,POU6F2,ATOH1,ISL1,PTCH1,B |
| GO:Molecular Functi | GO:0000981 | DNA-binding transcription factor activity, RNA polymerase II-specific | 4,353×10^-03 | 62 | 1375 | MECOM,SOX6,NPAS3,ESRRG,ZBTB20,EBF1,ZNF53 |
| GO:Biological Process | GO:0016477 | cell migration | 4,391×10^-03 | 1649 | 74 | AUTS2,PTPRT,DCC,HDAC9,DAB1,PRKG1,GPC6,FO |
| GO:Biological Process | GO:0007156 | homophilic cell adhesion via plasma membrane adhesion molecules | 4,438×10^-03 | 167 | 17 | PTPRT,ROBO2,TENM3,ROBO1,CDH12,PCDH15,KI |
| GO:Biological Process | GO:0099601 | regulation of neurotransmitter receptor activity | 4,519×10^-03 | 84 | 12 | DLG2,NLGN1,MEF2C,NRXN1,PPARGC1A,GRIA4,R |
| GO:Molecular Functi | GO:0140110 | transcription regulator activity | 4,792×10^-03 | 80 | 1935 | MECOM,SOX6,NPAS3,HDAC9,ESRRG,ZBTB20,EBF |
| GO:Molecular Functi | GO:0005515 | protein binding | 4,843×10^-03 | 412 | 14767 | DMD,AUTS2,IL1RAPL1,DLG2,PARD3B,MECOM,TE |
| GO:Biological Process | GO:0051254 | positive regulation of RNA metabolic process | 4,861×10^-03 | 1714 | 76 | AUTS2,MECOM,NPAS3,ESRRG,MEF2C,ZNF462,SK |
| GO:Molecular Functi | GO:0003700 | DNA-binding transcription factor activity | 4,950×10^-03 | 64 | 1441 | MECOM,SOX6,NPAS3,ESRRG,ZBTB20,EBF1,ZNF53 |
| GO:Cellular Compone | GO:0099060 | integral component of postsynaptic specialization membrane | 5,019×10^-03 | 74 | 10 | PTPRT,KCND2,DCC,GRID2,NLGN1,LRRC4C,NRG1,E |
| GO:Biological Process | GO:0045165 | cell fate commitment | 5,031×10^-03 | 267 | 22 | SOX6,TENM4,MEF2C,NRG1,ERBB4,RUNX2,EBF2,T |
| GO:Cellular Compone | GO:0008076 | voltage-gated potassium channel complex | 5,037×10^-03 | 90 | 11 | DLG2,KCND2,DPP10,CNTNAP2,DPP6,KCNIP4,KCN |
| GO:Biological Process | GO:2000026 | regulation of multicellular organismal development | 5,131×10^-03 | 1447 | 67 | IL1RAPL1,SOX6,DCC,ROBO2,GRID2,DAB1,NLGN1, |
| GO:Biological Process | GO:0021517 | ventral spinal cord development | 5,222×10^-03 | 45 | 9 | DAB1,MDGA2,PAX6,GLI3,RELN,GLI2,ISL1,PTCH1, |
| GO:Biological Process | GO:0007626 | locomotory behavior | 5,777×10^-03 | 189 | 18 | KCND2,NEGR1,DAB1,SLC4A10,NAV2,NRG1,PBX3, |
| GO:Biological Process | GO:0040007 | growth | 6,094×10^-03 | 967 | 50 | DMD,AUTS2,DCC,EYS,SLIT3,SLC4A10,TENM4,SYT1 |
| GO:Biological Process | GO:0051240 | positive regulation of multicellular organismal process | 6,206×10^-03 | 1455 | 67 | IL1RAPL1,SOX6,ROBO2,GRID2,ESRRG,ZBTB20,NL |
| GO:Biological Process | GO:0048562 | embryonic organ morphogenesis | 6,340×10^-03 | 292 | 23 | MEF2C,RARB,RUNX2,PAX6,NTN1,HAND2,MTHFD |
| GO:Biological Process | GO:0044057 | regulation of system process | 6,401×10^-03 | 623 | 37 | DMD,KCND2,CACNA2D3,NLGN1,PRKG1,TENM4,R |
| GO:Cellular Compone | GO:0008328 | ionotropic glutamate receptor complex | 6,456×10^-03 | 47 | 8 | DLG2,GRID2,NLGN1,GRIK3,GRIA4,GRIN2B,CACNG |
| GO:Biological Process | GO:0051094 | positive regulation of developmental process | 6,783×10^-03 | 1341 | 63 | DMD,IL1RAPL1,SOX6,ROBO2,GRID2,DAB1,NLGN1 |
| GO:Biological Process | GO:0001655 | urogenital system development | 6,823×10^-03 | 337 | 25 | ROBO2,MEF2C,KIRREL3,RARB,ESR1,PCSK5,FMN1, |
| GO:Biological Process | GO:0050905 | neuromuscular process | 6,998×10^-03 | 103 | 13 | GRID2,RBFOX1,PCDH15,NRXN1,CTNNA2,ANKFN1 |
| GO:Biological Process | GO:0072073 | kidney epithelium development | 7,371×10^-03 | 137 | 15 | ROBO2,MEF2C,RARB,FMN1,PBX1,ADAMTS16,GL |
| GO:Molecular Functi | GO:0008013 | beta-catenin binding | 7,409×10^-03 | 11 | 87 | PTPRT,CTNNA3,CTNNA2,GRIP1,ESR1,PARK2,TCF7 |
| GO:Biological Process | GO:0051674 | localization of cell | 7,840×10^-03 | 1828 | 79 | AUTS2,PTPRT,DCC,HDAC9,DAB1,PRKG1,GPC6,FO |
| GO:Biological Process | GO:0048870 | cell motility | 7,840×10^-03 | 1828 | 79 | AUTS2,PTPRT,DCC,HDAC9,DAB1,PRKG1,GPC6,FO |
| GO:Biological Process | GO:0001501 | skeletal system development | 7,987×10^-03 | 504 | 32 | SOX6,FOXP1,MEF2C,SLC10A7,RARB,RUNX2,COL1 |
| GO:Biological Process | GO:0010604 | positive regulation of macromolecule metabolic process | 8,013×10^-03 | 3483 | 130 | AUTS2,MECOM,NPAS3,DAB1,ESRRG,TBC1D5,ZBT |
| GO:Biological Process | GO:0048532 | anatomical structure arrangement | 8,022×10^-03 | 17 | 6 | DMD,DAB1,KCNA2,SEMA3A,HOXB1,HOXB2 |
| GO:Cellular Compone | GO:0098948 | intrinsic component of postsynaptic specialization membrane | 8,040×10^-03 | 78 | 10 | PTPRT,KCND2,DCC,GRID2,NLGN1,LRRC4C,NRG1,E |
| GO:Molecular Functi | GO:0046872 | metal ion binding | 8,074×10^-03 | 148 | 4260 | DMD,MECOM,TENM2,KCND2,LRP1B,EYS,SLIT3,H |
| GO:Biological Process | GO:0051093 | negative regulation of developmental process | 8,152×10^-03 | 977 | 50 | DMD,SOX6,DCC,ROBO2,DAB1,NLGN1,ZNF536,EF |
| GO:Cellular Compone | GO:0042734 | presynaptic membrane | 8,409×10^-03 | 150 | 14 | CNTN5,PTPRD,DENND1A,SYT1,NRXN1,NRG1,GRI |
| GO:Biological Process | GO:0009790 | embryo development | 8,671×10^-03 | 1035 | 52 | RCN1,TENM4,MEF2C,CACNA1C,AGBL4,PBX3,ERB |
| GO:Biological Process | GO:0051962 | positive regulation of nervous system development | 8,683×10^-03 | 276 | 22 | IL1RAPL1,ROBO2,GRID2,NLGN1,PTPRD,TENM4,R |
| GO:Cellular Compone | GO:0098878 | neurotransmitter receptor complex | 8,862×10^-03 | 49 | 8 | DLG2,GRID2,NLGN1,GRIK3,GRIA4,GRIN2B,CACNG |
| GO:Cellular Compone | GO:0005887 | integral component of plasma membrane | 9,003×10^-03 | 1645 | 68 | DLG2,TENM2,PTPRT,KCND2,DCC,DPP10,GRID2,C |
| GO:Biological Process | GO:0010092 | specification of animal organ identity | 1,063×10^-02 | 37 | 8 | ROBO2,ROBO1,MEF2C,GLI3,EXT1,AXIN2,ISL1,BM |
| GO:Biological Process | GO:0016043 | cellular component organization | 1,073×10^-02 | 6566 | 216 | DMD,AUTS2,IL1RAPL1,CADM2,DLG2,PARD3B,ME |
| GO:Cellular Compone | GO:0098797 | plasma membrane protein complex | 1,083×10^-02 | 694 | 36 | DMD,DLG2,KCND2,DPP10,GRID2,CNTNAP2,TBC1 |
| GO:Cellular Compone | GO:0099055 | integral component of postsynaptic membrane | 1,147×10^-02 | 116 | 12 | PTPRT,KCND2,DCC,GRID2,NLGN1,LRRC4C,NRG1,E |
| REACTOME | R-HSA-1638091 | Heparan sulfate\|heparin (HS-GAG) metabolism Homo sapiens | 1,152×10^-02 | 54 | 8 | EXT1;HPSE2;GPC3;GPC5;HS3ST5;GPC6;HS2ST1;H |
| GO:Cellular Compone | GO:0099061 | integral component of postsynaptic density membrane | 1,198×10^-02 | 51 | 8 | PTPRT,DCC,GRID2,LRRC4C,NRG1,ERBB4,CACNG2, |
| GO:Biological Process | GO:0002062 | chondrocyte differentiation | 1,198×10^-02 | 108 | 13 | SOX6,MEF2C,RARB,RUNX2,SOX5,RUNX1,GLI3,EXT |
| REACTOME | R-HSA-373752 | Netrin-1 signaling Homo sapiens | 1,252×10^-02 | 42 | 7 | TRPC6;DCC;SLIT1;SLIT3;UNC5C;NTN1;ROBO1 |
| GO:Biological Process | GO:0042391 | regulation of membrane potential | 1,253×10^-02 | 442 | 29 | DMD,KCND2,GRID2,NLGN1,MEF2C,CTNNA3,CAC |
| GO:Molecular Functi | GO:0043565 | sequence-specific DNA binding | 1,315×10^-02 | 70 | 1673 | MECOM,SOX6,NPAS3,ESRRG,ZBTB20,EBF1,ZNF53 |
| GO:Biological Process | GO:0021604 | cranial nerve structural organization | 1,318×10^-02 | 11 | 5 | DMD,KCNA2,SEMA3A,HOXB1,HOXB2 |
| GO:Biological Process | GO:0045935 | positive regulation of nucleobase-containing compound metabolic pr | 1,388×10^-02 | 1887 | 80 | AUTS2,MECOM,NPAS3,ESRRG,ZBTB20,MEF2C,ZN |
| GO:Biological Process | GO:0006357 | regulation of transcription by RNA polymerase II | 1,390×10^-02 | 2555 | 101 | AUTS2,MECOM,TENM2,SOX6,NPAS3,HDAC9,ESRR |
| GO:Biological Process | GO:0045596 | negative regulation of cell differentiation | 1,413×10^-02 | 697 | 39 | DMD,SOX6,DCC,DAB1,ZNF536,EFNA5,SEMA6D,Z |
| GO:Biological Process | GO:0051173 | positive regulation of nitrogen compound metabolic process | 1,416×10^-02 | 3083 | 117 | AUTS2,MECOM,NPAS3,DAB1,ESRRG,ZBTB20,ROB |
| GO:Cellular Compone | GO:0008091 | spectrin | 1,469×10^-02 | 9 | 4 | SPTBN1,EPB41L2,PRKCB,SPTB |
| GO:Biological Process | GO:0048663 | neuron fate commitment | 1,479×10^-02 | 64 | 10 | NRG1,PAX6,ZNF521,GLI3,GLI2,ATOH1,ISL1,EYA1,E |
| REACTOME | R-HSA-112316 | Neuronal System Homo sapiens | 1,578×10^-02 | 301 | 20 | KCND2;SYT1;PRKCB;KCNH7;KCNA2;CACNA2D3;G |
| GO:Molecular Functi | GO:0043169 | cation binding | 1,629×10^-02 | 149 | 4351 | DMD,MECOM,TENM2,KCND2,LRP1B,EYS,SLIT3,H |
| GO:Biological Process | GO:0048646 | anatomical structure formation involved in morphogenesis | 1,642×10^-02 | 1202 | 57 | ROBO2,GRID2,HDAC9,NRXN3,TENM4,ROBO1,ME |
| REACTOME | R-HSA-1500931 | Cell-Cell communication Homo sapiens | 1,654×10^-02 | 131 | 12 | KIRREL3;CDH4;CADM1;CADM2;DCC;MAGI2;CDH |
| GO:Molecular Functi | GO:0050839 | cell adhesion molecule binding | 1,682×10^-02 | 31 | 546 | DMD,TENM2,PTPRT,NRXN3,NLGN1,TENM3,PTPR |
| GO:Biological Process | GO:0048522 | positive regulation of cellular process | 1,695×10^-02 | 5757 | 193 | DMD,AUTS2,IL1RAPL1,MECOM,TENM2,SOX6,NP |
| GO:Biological Process | GO:0060047 | heart contraction | 1,729×10^-02 | 288 | 22 | DMD,KCND2,CACNA2D3,RYR3,SGCD,CTNNA3,CA |
| GO:Biological Process | GO:0014706 | striated muscle tissue development | 1,875×10^-02 | 380 | 26 | DMD,SOX6,HDAC9,RBFOX1,TENM4,MEF2C,SGCD, |
| GO:Cellular Compone | GO:0098936 | intrinsic component of postsynaptic membrane | 1,891×10^-02 | 122 | 12 | PTPRT,KCND2,DCC,GRID2,NLGN1,LRRC4C,NRG1,E |
| GO:Biological Process | GO:0048705 | skeletal system morphogenesis | 1,912×10^-02 | 226 | 19 | SOX6,MEF2C,RARB,RUNX2,SOX5,FMN1,FGR,MTH |
| GO:Cellular Compone | GO:0099146 | intrinsic component of postsynaptic density membrane | 2,098×10^-02 | 55 | 8 | PTPRT,DCC,GRID2,LRRC4C,NRG1,ERBB4,CACNG2, |
| GO:Biological Process | GO:0071625 | vocalization behavior | 2,353×10^-02 | 20 | 6 | CNTNAP2,NRXN3,NRXN1,FOXP2,GLI3,EXT1 |
| GO:Cellular Compone | GO:0070161 | anchoring junction | 2,396×10^-02 | 836 | 40 | DMD,PARD3B,CNTNAP2,CDH12,CTNNA3,KIRREL3 |
| GO:Biological Process | GO:0031345 | negative regulation of cell projection organization | 2,397×10^-02 | 189 | 17 | DCC,DAB1,NLGN1,SEMA6D,NRXN1,ZNF365,NTN1 |
| GO:Cellular Compone | GO:0044224 | juxtaparanode region of axon | 2,399×10^-02 | 10 | 4 | DLG2,CNTNAP2,KCNA2,KCNAB1 |
| GO:Biological Process | GO:1990138 | neuron projection extension | 2,484×10^-02 | 170 | 16 | AUTS2,SLIT3,SYT1,SEMA6D,PARK2,NTN1,TNR,MA |
| GO:Biological Process | GO:0007167 | enzyme linked receptor protein signaling pathway | 2,492×10^-02 | 1103 | 53 | PTPRT,PTPRD,ANKS1B,ROBO1,EFNA5,NRG3,NRX |
| GO:Biological Process | GO:2001257 | regulation of cation channel activity | 2,571×10^-02 | 190 | 17 | DMD,DLG2,NLGN1,MEF2C,NRXN1,PDE4B,PPARG |
| GO:Biological Process | GO:0006366 | transcription by RNA polymerase II | 2,585×10^-02 | 2723 | 105 | AUTS2,MECOM,TENM2,SOX6,NPAS3,HDAC9,ESRR |
| GO:Cellular Compone | GO:0030018 | Z disc | 2,592×10^-02 | 126 | 12 | DMD,RYR3,CACNA1C,NEBL,PDE4B,FHOD3,ANK3, |
| GO:Biological Process | GO:0035418 | protein localization to synapse | 2,594×10^-02 | 83 | 11 | DLG2,NLGN1,ANKS1B,GPC6,NRXN1,GRIP1,GPHN, |
| GO:Biological Process | GO:0048864 | stem cell development | 2,917×10^-02 | 84 | 11 | MSI2,FAM172A,SEMA6D,NRG1,ERBB4,HAND2,AL |
| GO:Biological Process | GO:0003015 | heart process | 2,974×10^-02 | 298 | 22 | DMD,KCND2,CACNA2D3,RYR3,SGCD,CTNNA3,CA |
| GO:Biological Process | GO:0048762 | mesenchymal cell differentiation | 3,150×10^-02 | 234 | 19 | MEF2C,FAM172A,SEMA6D,NRG1,ERBB4,TCF7L2, |
| REACTOME | R-HSA-4641265 | Repression of WNT target genes Homo sapiens | 3,171×10^-02 | 14 | 4 | TLE4;TCF7L2;CTBP2;AXIN2 |
| GO:Biological Process | GO:0009893 | positive regulation of metabolic process | 3,203×10^-02 | 3783 | 136 | AUTS2,MECOM,NPAS3,DAB1,ESRRG,TBC1D5,ZBT |
| GO:Biological Process | GO:0021772 | olfactory bulb development | 3,225×10^-02 | 31 | 7 | ROBO2,ROBO1,ERBB4,EXT1,SEMA3A,DLX5,SLIT1 |
| GO:Cellular Compone | GO:0000785 | chromatin | 3,464×10^-02 | 1526 | 62 | SOX6,NPAS3,ESRRG,EBF1,FOXP1,MEF2C,ZFHX4,P |
| GO:Biological Process | GO:0007166 | cell surface receptor signaling pathway | 3,535×10^-02 | 3210 | 119 | PTPRT,DCC,ROBO2,GRID2,SLIT3,DAB1,NLGN1,PT |
| REACTOME | R-HSA-6785631 | ERBB2 Regulates Cell Motility Homo sapiens | 3,811×10^-02 | 15 | 4 | DIAPH1;NRG3;ERBB4;NRG1 |
| REACTOME | R-HSA-5576894 | Phase 1 - inactivation of fast Na+ channels Homo sapiens | 3,819×10^-02 | 27 | 5 | KCND2;KCNIP4;CACNA2D3;CACNA1C;CACNG2 |
| REACTOME | R-HSA-5635851 | GLI proteins bind promoters of Hh responsive genes to promote transcription Homo sapiens | 3,819×10^-02 | 7 | 3 | PTCH1;GLI3;GLI2 |
| GO:Molecular Functi | GO:0022836 | gated channel activity | 3,942×10^-02 | 22 | 341 | KCND2,GRID2,CACNA2D3,RYR3,CACNA1C,GRIK3, |
| GO:Cellular Compone | GO:0031225 | anchored component of membrane | 4,256×10^-02 | 174 | 14 | LSAMP,NEGR1,CNTN5,OPCML,GPC6,EFNA5,MDG |
| GO:Biological Process | GO:0010557 | positive regulation of macromolecule biosynthetic process | 4,260×10^-02 | 1851 | 77 | AUTS2,MECOM,NPAS3,ESRRG,MEF2C,ZNF462,SK |
| GO:Biological Process | GO:0071840 | cellular component organization or biogenesis | 4,269×10^-02 | 6760 | 218 | DMD,AUTS2,IL1RAPL1,CADM2,DLG2,PARD3B,ME |
| GO:Biological Process | GO:0050770 | regulation of axonogenesis | 4,603×10^-02 | 159 | 15 | DCC,ROBO2,DAB1,ROBO1,LRRC4C,SEMA6D,NTN1 |
| REACTOME | R-HSA-1296072 | Voltage gated Potassium channels Homo sapiens | 4,894×10^-02 | 43 | 6 | KCND2;KCNH7;KCNA2;KCNA3;KCNAB1;KCNQ5 |
| GO:Biological Process | GO:0060443 | mammary gland morphogenesis | 4,957×10^-02 | 45 | 8 | NRG3,ESR1,NTN1,GLI3,GLI2,CSMD1,PTCH1,BMP4 |

**table 10: Coding mutations in the top 114 NCCM genes**

| #SAMPLE | Hugo_Symbol | CHROM | POS | REF | ALT | BED_START | BED_STOP | phyloP_valu | Variant_Cla | OREGANNO_ID | OREGANNO_Va | Variant_Cla |
|---|---|---|---|---|---|---|---|---|---|---|---|---|
| DO35538_PBC | FOSL2 | 2 | 28627104 | G | A | 28627103 | 28627104 | 8.796 | Missense_Mu | NOINFO | NOINFO | coding |
| DO35552_PBC | COL1A2 | 7 | 94038652 | G | A | 94038651 | 94038652 | 8.796 | Missense_Mu | NOINFO | NOINFO | coding |
| DO48890_PBC | COL1A2 | 7 | 94040249 | table | A | 94040248 | 94040249 | 7.026 | Missense_Mu | NOINFO | NOINFO | coding |
| DO35616_PBC | ZBTB18 | 1 | 244218364 | T | A | 244218363 | 244218364 | 6.318 | Missense_Mu | OREG0007882 | Outcome=NEG | coding |
| DO48906_PBC | ZBTB18 | 1 | 244217247 | TTTCCAC | T | 244217246 | 244217253 | 8.903 | In_Frame_De | NOINFO | NOINFO | coding |
| DO35621_PBC | PPARGC1A | 4 | 23830044 | A | G | 23830043 | 23830044 | -0.113 | Missense_Mu | NOINFO | NOINFO | coding |
| DO35625_PBC | AHDC1 | 1 | 27876103 | A | AGCGAAAGTAG | 27876102 | 27876103 | 6.299 | Frame_Shift | NOINFO | NOINFO | coding |
| DO35625_PBC | SUPV3L1 | 10 | 70962729 | G | A | 70962728 | 70962729 | 6.96 | Missense_Mu | NOINFO | NOINFO | coding |
| DO35676_PBC | CRB2 | 9 | 126128248 | C | T | 126128247 | 126128248 | 0.302 | Silent | NOINFO | NOINFO | coding |
| DO35723_PBC | ZNF462 | 9 | 109687545 | G | A | 109687544 | 109687545 | 8.796 | Missense_Mu | NOINFO | NOINFO | coding |
| DO35556_PBC | ZNF462 | 9 | 109686956 | C | T | 109686955 | 109686956 | 5.924 | Nonsense_Mu | NOINFO | NOINFO | coding |
| DO35739_PBC | HOXB2 | 17 | 46622734 | G | T | 46622733 | 46622734 | 0.506 | De_novo_Sta | OREG1269828 | Type=TRANSC | coding |
| DO35841_PBC | PRMT7 | 16 | 68386164 | T | G | 68386163 | 68386164 | 6.36 | Missense_Mu | OREG1194303 | Type=TRANSC | coding |
| DO48900_PBC | CTD-2144E22 | 16 | 34257197 | G | A | 34257196 | 34257197 | 0.313 | Silent | NOINFO | NOINFO | coding |
| DO48902_PBC | CTD-2144E22 | 16 | 34257180 | C | A | 34257179 | 34257180 | 0.31 | Missense_Mu | NOINFO | NOINFO | coding |
| DO48905_PBC | SUPT6H | 17 | 27022378 | G | A | 27022377 | 27022378 | 8.771 | Missense_Mu | OREG1507433 | Type=TRANSC | coding |
| DO48907_PBC | ABCC12 | 16 | 48149453 | C | T | 48149452 | 48149453 | 0.246 | Missense_Mu | NOINFO | NOINFO | coding |
| DO48914_PBC | PAX6 | 11 | 31823284 | T | TAGTCTC | 31823283 | 31823284 | 6.354 | In_Frame_In | OREG1833377 | Type=TRANSC | coding |
| DO48917_PBC | IFIH1 | 2 | 163144732 | G | A | 163144731 | 163144732 | 2.277 | Silent | NOINFO | NOINFO | coding |
| DO48919_PBC | GTF2E1 | 3 | 120500104 | G | A | 120500103 | 120500104 | -6.922 | Silent | OREG1807824 | Type=TRANSC | coding |
| DO48930_PBC | FGR | 1 | 27943703 | C | G | 27943702 | 27943703 | 8.885 | Splice_Site | NOINFO | NOINFO | coding |
| DO48965_PBC | AIM1L | 1 | 26658028 | C | T | 26658027 | 26658028 | 8.811 | Missense_Mu | NOINFO | NOINFO | coding |
| DO48968_PBC | ZFHX4 | 8 | 77768412 | C | T | 77768411 | 77768412 | 2.417 | Silent | NOINFO | NOINFO | coding |
| DO48971_PBC | ZFHX4 | 8 | 77761891 | G | T | 77761890 | 77761891 | 1.782 | Silent | NOINFO | NOINFO | coding |

table 11: NCCM rates in the significantly mutated genes of the MB cohort

| Gene | NCMs | NCCMs | NCCM Rate |
|---|---|---|---|
| MYCN | 31 | 4 | 1.9524 |
| LHX1 | 12 | 4 | 1.936 |
| PTCH1 | 19 | 5 | 1.8975 |
| OTX2 | 15 | 3 | 1.4376 |
| PRDM6 | 23 | 4 | 1.3204 |
| SYNCRIP | 16 | 3 | 1.2843 |
| GLI2 | 40 | 5 | 1.1055 |
| TBR1 | 6 | 2 | 0.98 |
| MYC | 25 | 2 | 0.9774 |
| KBTBD4 | 11 | 2 | 0.9748 |
| FAT1 | 28 | 3 | 0.9231 |
| MED12 | 20 | 2 | 0.9202 |
| SUFU | 18 | 3 | 0.9144 |
| KMT2D | 17 | 2 | 0.8922 |
| GFI1B | 14 | 2 | 0.8184 |
| CTNNB1 | 15 | 2 | 0.7614 |
| BAI3 | 83 | 7 | 0.7378 |
| CDK6 | 27 | 3 | 0.6972 |
| KDM6A | 28 | 3 | 0.6897 |
| PIK3CA | 18 | 2 | 0.687 |
| TCF4 | 45 | 4 | 0.6308 |
| ATM | 18 | 2 | 0.5934 |
| NSD1 | 23 | 2 | 0.557 |
| FBXW7 | 35 | 2 | 0.486 |
| CTDNEP1 | 17 | 1 | 0.4811 |
| GFI1 | 18 | 1 | 0.4748 |
| ZMYM3 | 18 | 1 | 0.4745 |
| DDX3X | 22 | 1 | 0.4365 |
| FMR1 | 30 | 1 | 0.4213 |
| PRKAR1A | 16 | 1 | 0.4197 |
| BRCA2 | 12 | 1 | 0.3659 |
| PTEN | 14 | 1 | 0.3245 |
| ZIC1 | 12 | 1 | 0.3169 |
| BCOR | 18 | 1 | 0.3105 |
| APC | 19 | 1 | 0.3032 |
| EPHA7 | 35 | 1 | 0.2656 |
| CHD7 | 21 | 1 | 0.264 |
| TP53 | 8 | 0 | 0.0 |
| GSE1 | 20 | 0 | 0.0 |
| SMARCA4 | 19 | 0 | 0.0 |
| TERT | 28 | 0 | 0.0 |
| KDM3B | 11 | 0 | 0.0 |
| SMO | 11 | 0 | 0.0 |
| CSNK2B | 10 | 0 | 0.0 |
| FLG | 16 | 0 | 0.0 |
| ARID1A | 16 | 0 | 0.0 |
| IDH1 | 15 | 0 | 0.0 |
| ARID2 | 18 | 0 | 0.0 |
| CREBBP | 19 | 0 | 0.0 |
| KMT2C | 23 | 0 | 0.0 |
| LDB1 | 15 | 0 | 0.0 |

**table 12: NCCM counts for genes in previously implicated MB pathways, and those well-known to be mutated in MB**

| Pathways | Genes Implicated | Total NCCMs | WNT | SHH | Group 3 | Group 4 |
|---|---|---|---|---|---|---|
| SHH Signalling | *Shh* | 1 | - | - | 1 (P) | - |
| | *Ptch1* | 5 | - | 2 (A) + 1(P) | 1 (P) | 1 (P) |
| | *Smo* | 0 | - | - | - | - |
| | *Gli3* | 9 | - | 1(A) | 4 (P) | 4 (P) |
| | *Gli2* | 3 | 1 (P) | 2 (A) | - | 4 (P) |
| | *Gli1* | 1 | - | - | 1 (P) | - |
| | *Noggin* | 2 | - | 2 (A) | - | - |
| | *SuFu* | 3 | - | 1(A) | 1 (P) | 1 (P) |
| | *KIF7* | 1 | - | 1(A) | - | - |
| WNT Signalling | *WNT* | 3 | - | 3 (A) | - | - |
| | *Frizzled* | 3 | - | 2 (A) + 1(P) | - | - |
| | *Beta catenin* | 6 | - | 1 (P) | 3 (P) | 1 (A) + 1(P) |
| | *Axin2* | 7 | - | - | 2 (P) | 5 (P) |
| | *APC* | 1 | - | - | 1 (P) | - |
| | *ROR1* | 1 | - | 1 (A) | - | - |
| | *ROR2* | 2 | - | 2 (A) | - | - |
| | *CK1 (delta)* | 0 | - | - | - | - |
| | *Lrp5* | 1 | - | - | - | 1 (P) |
| | *Lrp6* | 0 | - | - | - | - |
| | *DVL1* | 1 | - | - | 1 (P) | - |
| | *DVL2* | 3 | - | 2 (A) | 1 (P) | - |
| MYC Regulation | *mTOR* | 1 | - | 1 (A) | - | - |
| | *MYC* | 2 | - | - | 1 (P) | - |
| | *MYCN* | 4 | - | 1 (A) | - | 4 (A) |
| | *MYCL1* | 1 | - | 1 (A) | - | - |
| | *MYCBP* | 3 | - | - | - | 2 (P) + 1(A) |
| | *NMI (n-myc interactor)* | 4 | - | - | 1 (P) | 3 (P) |
| | *ARF1* | 2 | - | 1 (A) | 1 (P) | - |
| | *MAX* | 1 | - | - | 1 (P) | - |
| | *GSK3B* | 3 | - | 1 (A) | - | 2 (P) |
| Genes known to be prominently mutated in MB | *DVL1* | 1 | | | 1 (P) | |
| | *GFI1* | 1 | - | 1(A) | - | - |
| | *GFI1B* | 2 | - | 1(A) | 1 (P) | - |
| | *KBTBD4* | 2 | - | 2(A) | - | - |
| | *KDM6A* | 3 | - | 1 (P) + 1(A) | - | 1 (P) |
| | *LPR5* | 1 | | | | 1 (P) |
| | *PRDM6* | 4 | - | 1 (P) | 3(P) | - |
| | *PRKCD* | 1 | - | 1(A) | - | - |
| | *TRAAP* | 2 | | 1 (A) | | 1 (A) |
| | *ZMYM3* | 3 | 1 (P) | 1(A) | 1 (P) | - |

**A --> Adult Cohort**

**P --> Pediatric Cohort**

table 13: sTRAP analysis for NCCMs associated with *BMP4* and *HOXB*-locus genes

| Mutation Name | op NCCM Ger | Sample ID | Gender | Age at diagnosis | Molecular classification for the tumor | om# | Posi. | Ref. | Alt. | Mutation Classification | loP | Matrix ID (Duplicated Matrices shaded in blue) | Matrix Name | Wildtype Bound Sites. (The footprint view shows the bound sites in red uppercase). | Mutant Bound Sites (The footprint view shows the bound sites in red uppercase). | Difference log(p) for two sequences | Location (s) (Strand) | Wt Weight Score(s) | Mut Weight Score(s) | Wt sequence *p*-value | Mut sequence *p*-value |
|---|---|---|---|---|---|---|---|---|---|---|---|---|---|---|---|---|---|---|---|---|---|
| BMP4.NCCM-3 | *BMP4* | DO35553 | female | 13 | Group3 | 14 | 54414824 | T | A | IGR | 6,3 | MA0442.1 | **SOX10** | - | caatttttattggtttccACAAAGctgctccttttggagga | -1,57 | 19-24(-) | - | **6,75** | 0,67 | <0.018 |
| | | | | | | | | | | | | MA0122.1 | **Nkx3-2** | caatttttattggttTCCACTAAGctgctccttttggagga | - | 1,55 | 16-24(-) | **6,52** | - | **0,02** | 0,64 |
| BMP4.NCCM-2 | *BMP4* | DO35564 | male | 8 | Group4 | 14 | 54369539 | C | T | IGR | 1,3 | MA0027.1 | **En1** | - | ttttagaaaaGAAAATTATTTgactacataatcagacgtta | -1,54 | 11-21(-) | - | **5,51** | 0,05 | 0,00 |
| | | | | | | | | | | | | MA0068.1 | **Pax4** | ttttagaaaaGAAAATTATTCGACTACATAATCAGACGTTa | - | 0,777 | 11-40(+) | **6,73** | - | **0,05** | 0,29 |
| BMP4.NCCM-1 | *BMP4* | DO35575 | male | 6 | Group4 | 14 | 54351945 | C | T | IGR | 4,7 | MA0075.1 | **Prrx2** | - | ttgattaaaagttttctTAATTAactggaatgtttccttag | -1,73 | 18-22(-);19-23(+) | - | **6.71;6.71** | 0,17 | <0.003 |
| | | | | | | | | | | | | MA0132.1 | **Pdx1** | - | ttgattaaaagttttcTTAATTAActggaatgtttccttag | -1,13 | 17-22(+);19-24(-) | - | **6.05;6.05** | 0,22 | 0,02 |
| | | | | | | | | | | | | MA0125.1 | **Nobox** | - | ttgattaaaagttttCTTAATTAACtggaatgtttccttag | -0,964 | 16-23(-);18-25(+) | - | **5.62;6.40** | 0,20 | 0,02 |
| | | | | | | | | | | | | MA0153.1 | **HNF1B** | - | ttgattaaaagttTTCTTAATTAACtggaatgtttccttag | -0,885 | 14-25(+) | - | **7,35** | 0,10 | 0,01 |
| | | | | | | | | | | | | MA0063.1 | **Nkx2-5** | - | ttgattaaaagttttcTTAATTAActggaatgtttccttag | -0,817 | 17-23(+);18-24(-) | - | **5.53;5.53** | 0,18 | 0,03 |
| BMP4.NCCM-5 | *BMP4* | DO35857 | male | 3 | Group3 | 14 | 54512866 | C | CT | IGR | 6,0 | MA0142.1 | **Pou5f1** | - | gacagacgtttttatttgcaCTTTTTTTTTCAAATctatcca | -1,08 | 21-35(+) | - | **8,14** | 0,13 | 0,01 |
| | | | | | | | | | | | | MA0143.1 | **Sox2** | - | gacagacgtttttatttgcACTTTTTTTTTCAAAtctatcca | -0,763 | 20-34(+) | - | **6,40** | 0,24 | 0,04 |
| BMP4.NCCM-4 | *BMP4* | DO48922 | male | 10 | Group3 | 14 | 54421098 | G | A | Intron | 4,6 | MA0043.1 | **HLF** | tcagatcttaatgtggccgaGGTTTTACAACTcccgacccg | - | 0,546 | 21-32(+) | **5,31** | - | **0,05** | 0,17 |
| | | | | | | | | | | | | MA0131.1 | **MIZF** | - | tcagatcttaatgtgGCCGAAGTTTtacaactcccgacccg | -0,337 | 16-25(-) | - | **5,16** | 0,08 | 0,04 |
| HOXB1.NCCM-1 | *HOXB1/B2* | DO35547 | male | 3 | SHH | 17 | 46545510 | C | T | lincRNA | 2,4 | MA0101.1 | **REL** | - | gcaccatttcttaTGGGTTTTTCtgaaccctgcccatatct | -1,08 | 14-23(+) | | **6,70** | 0,46 | 0,04 |
| | | | | | | | | | | | | MA0137.2 | **STAT1** | gcaccatttcttatgGGTTTCTCTGAACCCtgcccatatct | - | 0,676 | 16-30(+) | **5,01** | - | 0,07 | 0,32 |
| HOXB1.NCCM-2 | *HOXB1/B2* | DO48957 | male | 5 | Group3 | 17 | 46640029 | A | G | Intron | 6,3 | MA0132.1 | **Pdx1** | atgataaaatcAATTAAATTACggccacttagggcccagcc | atgataaaatcAATTAAattgcggccacttagggcccagcc | 0,29 | 12-17(-) | **6,05** | **6,05** | 0,02 | 0,04 |
| HOXB1.NCCM-3 | *HOXB1/B2* | DO35581 | male | 3 | Group3 | 17 | 46655880 | G | T | 5'UTR | 8,7 | MA0042.1 | **FOXI1** | gaccccactccagccaaAGAGGTTTATTTccccttcttccc | gaccccactccagccaaAGATGTTTATTTccccttcttccc | -1,35 | 18-29(+) | 5,6547 | **10,48** | 0,10 | 0,00 |
| | | | | | | | | | | | | MA0040.1 | **Foxq1** | gaccccactccagccaaAGAGGTTTATTtccccttcttccc | gaccccactccagccaaAGATGTTTATTtccccttcttccc | -1,26 | 18-28(+) | 5,03101 | **9,14** | 0,10 | 0,01 |
| | | | | | | | | | | | | MA0066.1 | **PPARG** | gaccccactccagccaAAGAGGTTTATTTCCCCTTCttccc | - | 1,11 | 17-36(-) | **3,98** | - | 0,02 | 0,20 |
| | | | | | | | | | | | | MA0031.1 | **FOXD1** | - | gaccccactccagccaaagATGTTTATttccccttcttccc | -0,908 | 20-27(-) | - | **6,86** | 0,27 | 0,03 |
| | | | | | | | | | | | | MA0041.1 | **FOXD3** | - | gaccccactccagccaaAGATGTTTATTTccccttcttccc | -0,598 | 18-29(+) | | **6,30** | 0,30 | 0,08 |
| | | | | | | | | | | | | MA0030.1 | **FOXF2** | - | gaccccactccagccaaaGATGTTTATTTCCCcttcttccc | -0,365 | 19-32(-) | - | **5,97** | 0,09 | 0,04 |
| HOXB1.NCCM-5 | *HOXB1/B2* | DO35575 | male | 6 | Group4 | 17 | 46703946 | G | A | 5'Flank | 2,3 | MA0069.1 | **Pax6** | - | tccgcaccgcactccatATCAAGGATGGATTgttttatgct | -1,09 | 18-31(+) | | **5,11** | 0,25 | 0,02 |
| | | | | | | | | | | | | MA0142.1 | **Pou5f1** | - | tccgcaccgCACTCCATATCAAGGatggattgttttatgct | -0,679 | 10-24(-) | | **5,03** | 0,35 | 0,07 |
| | | | | | | | | | | | | MA0036.1 | **GATA2** | tccgcaccgcactccaTATCGaGGATGgattgttttatgct | tccgcaccgcactccaTATCAaGGATGgattgttttatgct | 0,326 | 17-21(-); 23-27(+) | **4.60; 4.42** | 4.24; 4.42 | 0,02 | 0,04 |

**table 14: oligos used for the EMSA experiments.**

| Allele name | Forward Oligo | Reverse Oligo |
|---|---|---|
| WT_BMP4 | TATTGGTTTCCACTAAGCTGCTCCTTTT | AAAAGGAGCAGCTTAGTGGAAACCAAT |
| MUT_BMP4 | TATTGGTTTCCACAAAGCTGCTCCTTTT | AAAAGGAGCAGCTTTGTGGAAACCAATA |
| WT_HOXB1/HOXB2_1 | TTTCTTATGGGTTTCTCTGAACCCTGCCC | GGGCAGGGTTCAGAGAAACCCATAAGAAA |
| MUT_HOXB1/HOXB2_1 | TTTCTTATGGGTTTTTCTGAACCCTGCCC | GGGCAGGGTTCAGAAAAACCCATAAGAAA |
| WT_HOXB1/HOXB2_2 | CCGCACTCCATATCGAGGATGGATTGTTT | AAACAATCCATCCTCGATATGGAGTGCGG |
| MUT_HOXB1/HOXB2_2 | CCGCACTCCATATCAAGGATGGATTGTTT | AAACAATCCATCCTTGATATGGAGTGCGG |
| WT_HOXB1/HOXB2_3 | ACTCCAGCCAAAGAGGTTTATTTCCCCTT | AAGGGGAAATAAACCTCTTTGGCTGGAGT |
| MUT_HOXB1/HOXB2_3 | ACTCCAGCCAAAGATGTTTATTTCCCCTT | AAGGGGAAATAAACATCTTTGGCTGGAGT |
